# Constitutive Photomorphogenesis Protein 1 homolog (COP1) sustains nuclear factor-4 alpha function in human hepatocyte models

**DOI:** 10.1101/2024.01.11.575239

**Authors:** Sébastien Soubeyrand, Paulina Lau, Ruth McPherson

**Affiliations:** Atherogenomics Laboratory, University of Ottawa Heart Institute, Ottawa, Canada; Department of Medicine, Ruddy Canadian Cardiovascular Genetics Centre, University of Ottawa Heart Institute, Ottawa, Canada

**Keywords:** COP1, HNF4A, hepatocarcinoma, hepatoblastoma, hepatocyte, HepG2, HuH-7, MTTP, TRIB1

## Abstract

Constitutive Photomorphogenesis Protein 1 homolog (COP1) is a conserved E3 ligase with key roles in several biological systems. Prior work in hepatocyte derived tumors categorized COP1 as an oncogene but its role in untransformed hepatocytes remains largely unexplored. Here we have investigated the role of COP1 in primary human hepatocytes as well as in two transformed hepatocyte models, HepG2 and HuH-7 cells. Contrary to a previous report, COP1 suppression via siRNA had no noticeable impact on HepG2 and HuH-7 proliferation and was associated with contrasting rather than congruent transcriptome changes. Clustering analyses identified patterns indicative of perturbed metabolism in primary hepatocytes and HepG2 cells whereas patterns pointed to cell proliferation impacts in HuH-7 cells. In HepG2 and primary hepatocytes, COP1 suppression reduced the expression important hepatic regulators and markers, which could be restored by the introduction of a siRNA resistant COP1 transgene. COP1 downregulation reduced hepatic nuclear factor-4 alpha (HNF4A) abundance and function, as assessed by lower abundance of key HNF4A targets and reduced APOB secretion. HNF4A restoration partially rescued COP1 silencing in HepG2 cells. This study identifies COP1 as a key regulator of hepatocyte function, in part via HNF4A. COP1 was required to maintain HNF4A abundance and function in primary hepatocytes and in HepG2 cells, but not in HuH-7 cells. Lastly, by demonstrating contrasting roles of COP1 in HuH-7 and HepG2 cells, our findings also challenge previous work linking COP1 to hepatic tumorigenesis.

## Introduction

The genome is predicted to encode over 600 RING (really interesting new gene) type E3 ligases that catalyze the las step in substrate ubiquitination^1^. Constitutive Photomorphogenic Protein 1 (COP1)/*RFWD2* is an essential, evolutionarily conserved, E3 ligase that plays a pivotal regulatory role by promoting ubiquitination and degradation of cognate cargos ^2–10^. This can be achieved via direct binding (e.g p53) or may require the presence of additional factors (e.g., *tribbles* proteins for CEBPA, ACC1) ^7,11^. Of relevance to liver function, COP1 suppresses lipogenesis and gluconeogenesis in hepatocytes by facilitating CEBPA and FOXO1 degradation ^10,12^.

In addition, COP1 has also been linked to oncogenesis. Hypomorphic *cop1* mice are susceptible to developing tumors ^5^. COP1 deficiency in mouse prostate leads to increased cell proliferation and hyperplasia ^3^. Moreover, COP1 induces acute myeloid leukemia in murine overexpression models ^13^. In humans with gastric carcinoma, low COP1 levels are associated with a poor prognosis and have been shown to increase proliferation in a gastric cancer cell line ^14^. However, previous studies have been inconsistent regarding the role of COP1 function in hepatic cancer, with tumor suppressor or oncogenic properties ^5,11,15,16^. HepG2 and HuH-7 cells, two well characterized liver tumor derived cell populations, have been reported to undergo profound growth inhibition following COP1 suppression ^15^. COP1 suppression was shown to elicit growth inhibition and apoptotic regression and, in a nude mouse model *COP1* siRNA, reduced xenograft growth ^15^

HepG2 and HuH-7 cells were established over 40 years ago. HuH-7 is an established cell line derived from a 57-year-old Japanese hepatoma whereas the HepG2 cell line was derived from the lobectomy specimen of a 15-year old Caucasian male ^17,18^. Importantly, while both cell lines have a neoplastic origin, HuH-7 are hepatocarcinoma cells (HCC) whereas HepG2 cells are hepatoblastoma (HBB), despite the latter being often erroneously referred to as hepatocarcinoma ^19^. HBB typically share features reminiscent of fetal liver, are less aggressive and respond better to chemotherapy than HCC ^20^. As expected for transformed cell lines, HuH-7 and HepG2 genomes are rearranged and mutated ^21–24^. Notwithstanding their transformed phenotypes, they are widely used as surrogate primary hepatocyte models. But this comes at a price, as these cells are known to differ substantially from plated hepatocytes. For instance, they display impaired lipoprotein secretion ^25,26^. The two cell lines also differ functionally from each other in numerous points including their dependence on glucose and lipogenic ability ^27,28^.

In this work, COP1 is investigated in HepG2 and HuH-7 cells as well as in primary hepatocytes to shed light on its role in both pathological and physiological liver settings. We report several discrepancies with previously published work on HepG2 and HuH-7 cells. Finally, we identify HNF4A as a functional target of COP1.

## Materials and Methods

### Cell culture

HepG2 and HuH-7 cells were maintained in 5 mM Glucose DMEM (Gibco) supplemented with 10% fetal bovine serum and antibiotic supplements (Gibco). HepG2 and HuH-7 cells were obtained from the American Type Culture Collection and the Japanese Collection of Research Biosources Cell Bank, respectively. All siRNAs were of Silencer™ Select grade (Ambion) to minimize off-targeting. HepG2 and HuH-7 cells were transfected at 30%-50% confluence in 0.5 ml media in 24 well plates using 10 nM siRNA (final) and 1 µl of RNAiMax. The siRNA transfection mix was prepared in 100 µl Optimem (Gibco). Treatment was continuous for 96 h. Cryopreserved primary hepatocytes (HMCPMS), media and media supplements were obtained from ThermoFisher Scientific. Two distinct lots, originating from 2 distinct donors, were used: lot Hu8306 (transcription array analysis of COP1 suppressed hepatocytes; 27 YO, BMI 29.5) and Hu8287 (replication with si33 and si91; 49 YO, BMI 19.6) and were thawed in thawing media and then plated in plating media for 6 h in 12-well plates (1 ml media per well) pre-coated with collagen I (1 vial was seeded in 10 and 14 wells, respectively). Media was changed to culture medium which was replaced every 24 h. Replicates (3) consisting of 2 treatments each (COP1 silencing oligonucleotides and NT1si) were obtained by staggering transfection by 24 h i.e. 4, 5 and 6 days after initial plating; 10 nM siRNA and 3 µl of RNAiMax in 100 ul Optimem were used per transfection of cells. Cell extracts were obtained by washing 3L×Lin PBS and either lysing for RNA (TriPure reagent, Roche) or protein extraction (50 mM Tris-HCl, 1% NP40, 0.15 M NaCl, pH 8) supplemented with phosphatase (PhoStop, Roche) and protease inhibitors (Complete, Roche).

### Expression constructs

COP1 and HNF4A were synthesized *in vitro* by BioBasic (Markham, Ontario) and assembled in the lentiviral PLVX plasmid. COP1 and HNF4A are tagged with C-terminal FLAG epitope and Hemagglutinin tags, respectively. COP1 has been codon optimized to be compatible with gene synthesis. Silent mutations over the siRNA targeted region (sequence in Supplementary Materials) were included to confer resistance to COP1 siRNA. Endogenous HNF4A is generated from 2 distinct promoters and is alternatively spliced. The variant used here corresponds to NP_000448.3/HNF4α1, the canonical and fully active form, and includes a C-terminal HA tag. Sequences are detailed in the supplemental materials section.

### Viral particle generation and stable cell line generation

Viral particles were generated by co-transfection of 293FT cells with the empty PLVX plasmid or PLVXCOP1(FLAG) with psPAX2 and pMD2.G viral particles harvested during the 16-72 h window post-transfection were concentrated with Lenti-X (Clontech) and stored at -80°C. Stable pools were generated using incremental doses of virus and puromycin resistance (3 µg/ml). Unless specified otherwise, cell populations selected with the lowest doses of virus compatible with puromycin resistance were retained.

### Western blotting

SDS-PAGE Gels (8% or 10%) were transferred to nitrocellulose membranes (Bio-Rad) using a Trans-blot apparatus (2.5 A constant, 6 min). Membranes were briefly ponceau stained to ensure even loading, fully destained with 3 x 1 min PBS over 3 min and blocked in Intercept buffer (LI-COR) for 1 h. Blots were then rinsed once in PBS/0.1% Tween and detection was performed in PBS/0.1% Tween for 16 h at antibody concentrations of 0.5-1 µg/ml. Secondary antibodies were from LI-COR (680 and 800) and were used as 1:20,000 dilutions. Four 30 second PBS rinses were performed between incubations. Acquisition was performed on an Odyssey Imager (LI-COR) using the default settings. All images shown were modified for contrast and intensity only (kappa=1) and were within the dynamic range of the imager. Antibodies used are listed in Supplementary Materials.

### Cell proliferation assays

Cells were assessed 96 h post incubation onset by the addition of 1/10 volume of alamarBlue (ThermoFisher Scientific) and a 1 h incubation. Fluorescence was measured in a Dynatek (Synergy) fluorometer, using 560 nm excitation and 590 nm emission, using AutoScale. Values from a cell-free well control was included and subtracted from the sample values.

### APOB enzyme-linked immunosorbent assay (ELISA)

APOB in the media of si33 and si91 was measured by ELISA (Mabtech). Briefly, media from the last 24 h of siRNA treatment was harvested, centrifuged at 1000 X g (2 min) to remove cell debris and frozen at -80 C until analyzed. Thawed media was diluted 10 X in PBS for assay to fit within the linear range of the APOB standard curve. Substrate (1-Step Ultra TMB-ELISA; ThermoFisher Scientific) was added for 10 min and reactions were stopped with the addition of 1M sulfuric acid (final). Absorbance (450 nm) was measured on a Dynatek fluorometer (Synergy). Values were corrected to RNA levels (harvested concurrently) and are expressed relative to the matching control siRNA; similar results were obtained without correction (i.e., using media values only).

### Real-time RNA quantification (RT-qPCR)

Total RNA was extracted from culture plates using Tri-Reagent (Roche) and isolated using Direct-Zol RNA miniprep kits (ZYMO RESEARCH). RNA (200 ng) was reverse transcribed in a 10 µl reaction using a Roche kit, using a 1:1 mixture of oligo dT and random hexamer primers (Roche). Quantification was performed on a LightCycler 480 (Roche). Target of interest values were expressed relative to the corresponding PPIA values using the ΔΔCt method. Oligonucleotide primers are detailed in the Supplementary Materials section.

### Transcription array analysis

RNA from primary hepatocytes and tumor derived liver cells (3 biological replicate sets, each consisting of a COP1 silenced and a NT1 control), was isolated 96 h post-silencing and sent to The Centre for Applied Genomics (The Hospital for Sick Children, Toronto, Ontario, Canada) for processing. A HuH-7 sample (COP1si replicate) failed bioanalyzer analysis and was discarded. Biotin-labeled cRNA was hybridized on HuGene-2_0-st arrays (Affymetrix) and results were processed with the Expression Console and analyzed with the Transcriptome Analysis Console from Affymetrix using default settings. False Discovery Rates (FDR) were determined using the Benjamini– Hochberg method. The data are available at the Gene Expression Omnibus (GSE206116).

### Bioinformatic analyses

Mapping of transcriptional consequences of COP1 suppression was performed using the WebGestalt interface (http://www.webgestalt.org/) ^29^. Over-representation analysis (ORA) and Gene Set Enrichment analyses (GSEA) were used. Affymetrix Gene IDs were mapped via WebGestalt. GSEA was performed using the complete transcriptome fold-change dataset. Nominal changes were used for ORA. Background list was “affy hugene 2.0 stv1”. Only FDR significant hits (q < 0.05) are shown. Impacted transcripts were mapped to Kyoto Encyclopedia of Genes and Genomes (KEGG), Gene Ontology (GO) and Reactome categories^30–33^. For Ingenuity Pathway Analysis (IPA) core analysis, molecules undergoing fold changes > 1.5 (ANOVA p < 0.05) as determined by the Transcriptome Analysis Console (Affymetrix) output were selected. The Ingenuity Knowledge Base (Genes + Endogenous Chemicals) was used as reference set. High confidence and experimentally observed data were included in the analyses. For IPA Upstream Regulator analysis, the analysis examines known targets of upstream regulators present in the dataset provided and compares the direction of the changes observed to the expected direction. Fisher’s exact test is used to measure statistical enrichment of impacted targets present in the sample list versus a database of regulator-targets.

### Statistical analyses

Statistical significance was tested using (paired) Student’s t-test for 2 groups or one-way ANOVA for multiple treatment groups, as indicated.

## Results

### COP1 suppression is well tolerated in HepG2 and HuH-7 hepatoma cells

While examining the role of COP1 in transformed liver models, we found that COP1 suppression (∼85% in HuH-7 and ∼70% in HepG2) had little impact on HepG2 and HuH-7 cell abundance (Fig 1A, B). This observation contrasts with previously published results, showing that a comparable COP1 suppression strongly impaired proliferation of both cell types ^15^. Western blotting for COP1 confirmed COP1 suppression at the protein level (Fig 1C). To ensure specificity, HuH-7 and HepG2 cells were treated with 2 additional siRNAs, neither of which showed toxicity despite comparable COP1 suppression (Fig S1). As suppression of COP1 mRNA expression was comparable across both studies, other sources of discrepancy were considered, including the use of different transfection reagents, Lipofectamine 2000 versus RNAiMAX, used in the current study. We reasoned that Lipofectamine 2000 may synergize with COP1 suppression to be cytotoxic. This was tested in HuH-7; however, findings were comparable to RNAiMax (Fig S2). While the reasons behind this discrepancy were not further investigated, COP1 suppression in the absence of significant cell death warranted a fresh assessment of COP1 function in these 2 cell lines.

**Figure 1.**
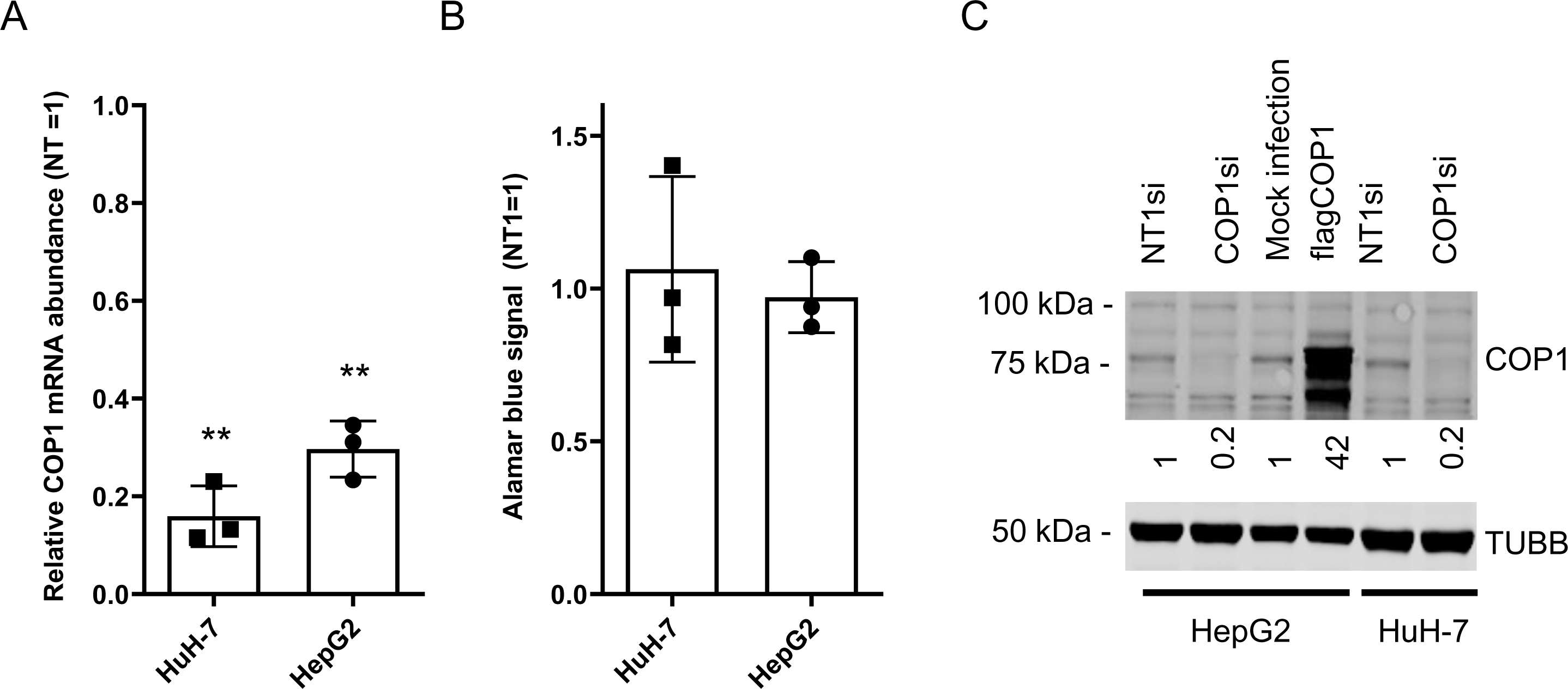
COP1 suppression in HepG2 and HuH-7 cells does not affect proliferation. HuH-7 and HepG2 cells were transfected with a COP1 targeting siRNA (COP1si) or a non-specific control (NT1) for 96 h prior to analysis. A, RNA isolation and quantification by qRT-PCR. B, cell abundance estimation by Alamar blue of siRNA treated cells. Three biological repeats (and means ± SD) are shown. Statistical significance of NT1si vs COP1si values was assessed using Students t-test. ** p<0.01. C, Western blot of HuH-7 and HepG2 cells treated with siRNA or transduced (for HepG2) with a flagCOP1 expressing virus for 96 h. Quantification representative of 3 biological repeats. TUBB, Tubulin beta chain.

### Over-representation analyses of transcriptome changes following COP1 suppression in transformed and primary hepatocytes

HepG2 and HuH-7 transcriptomes following COP1 silencing showed relatively little overlap, sharing only 423 nominally significant hits (out of 6057 (7% of nominal hits) and 2861 (15% nominal hits) for HepG2 and HuH-7, respectively), indicating divergent trajectories with respect to the role of COP1 (Fig S3). These hits were mapped to pathways (Reactome and the Kyoto Encyclopedia of Genes and Genomes (KEGG)) and Gene Ontology (GO) terms by over-representation analysis (ORA), which identified enrichment of terms linked to cell cycle regulation and DNA metabolism (Table S1). Comparison of nominally impacted transcripts within each cell type revealed dichotomous responses for HepG2 and HuH-7: transcriptome changes in HepG2 mainly mapped to metabolic processes while in HuH-7, changes were prominently consistent with cell cycle regulation terms (Table 1, Table S2).

**Table 1.**
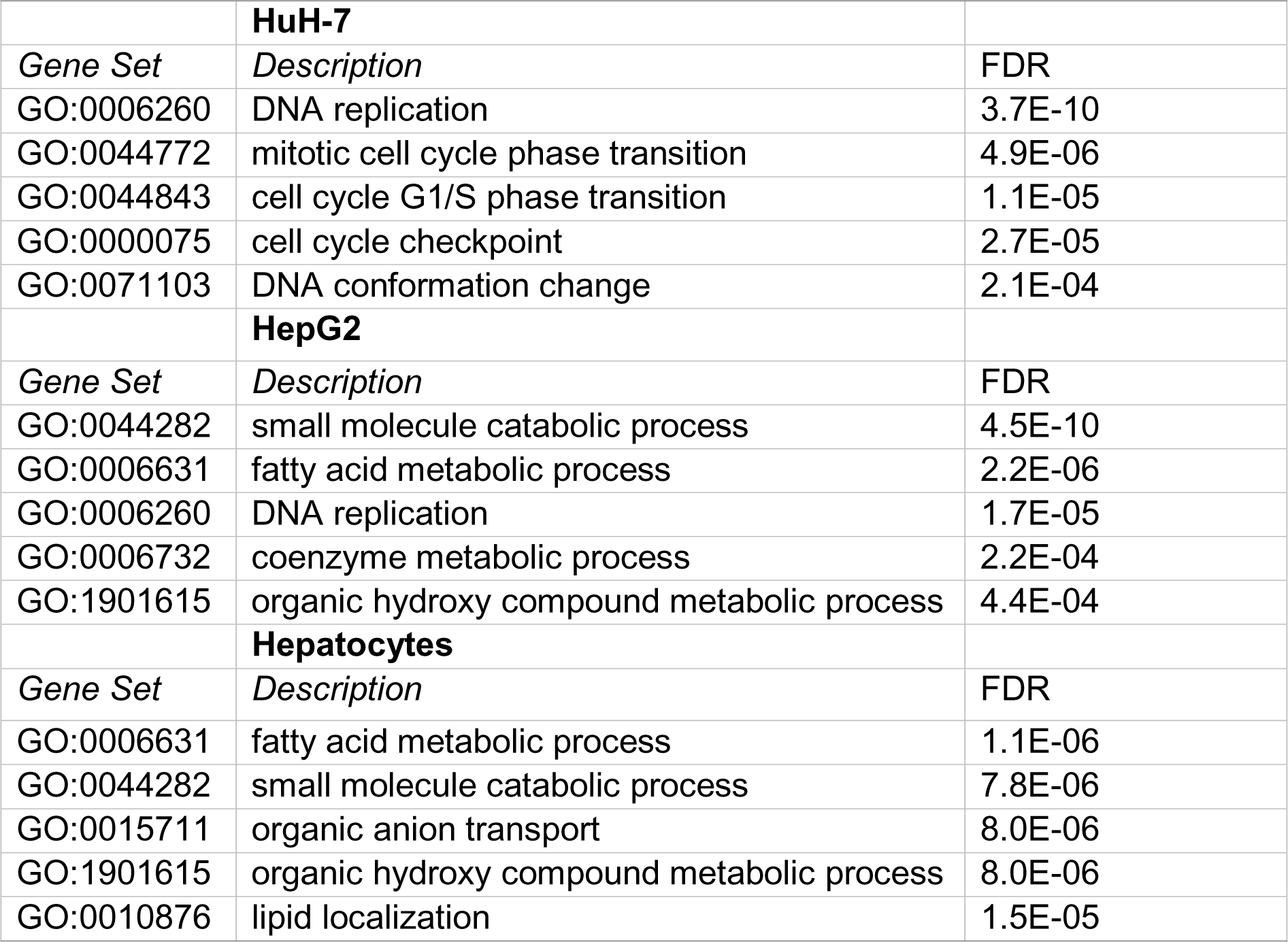
Over-representation analysis of transcripts nominally affected in COP1 silenced HepG2, HuH-7 and primary hepatocytes. Transcriptome data were mapped Gene Ontology terms (Biological_Process_noRedundant). The top five FDR significant hits are shown for each cell type. A complete table (including KEGG and Reactome sets) is included as Table S2.

These results pointed to divergent roles of COP1 in transformed hepatocyte models. To delineate the physiological role of COP1, COP1 was next targeted in primary human hepatocytes. Consistent with transformed models, siRNA mediated COP1 knock-down resulted in the suppression of COP1 at the mRNA (reduced to 28 ± 10% of control value, 95% C.I.) and protein levels (reduced to 25 ± 28% of control value, 95% C.I.) (Fig 2). The suppression resulted in 4262 nominally changed transcripts, of which only a small fraction was also impacted in HepG2 (848) and HuH-7 cells (299) (Fig S3). Over-representation analysis indicated that primary hepatocytes underwent transcriptome changes mapping primarily to metabolic processes (Table 1, Table S2). Mapping specificity was confirmed by transcript randomization (Table S3). To obtain a more granular view of these findings, ORA was repeated on nominal transcripts segregated by direction of effect. Enrichment analysis of reduced abundance transcripts revealed an extensive overlap between HepG2 and primary hepatocytes, with an enrichment of metabolic and lipid-related terms (Table S4). Moreover, the analysis uncovered opposite patterns in HepG2 versus HuH-7: proliferation related gene sets were upregulated in HepG2 but downregulated in HuH-7 (Table S4, pink highlights). By comparison, only one proliferation pathway (“epithelial cell proliferation”) was positively enriched in primary hepatocytes. Rather, COP1 suppression resulted in the upregulation of transcripts enriched in immunity related gene sets (“acute inflammatory response”, “lymphocyte mediated immunity”, etc.) suggesting that COP1 contributes to the regulation of the hepatocyte immune environment.

**Figure 2.**
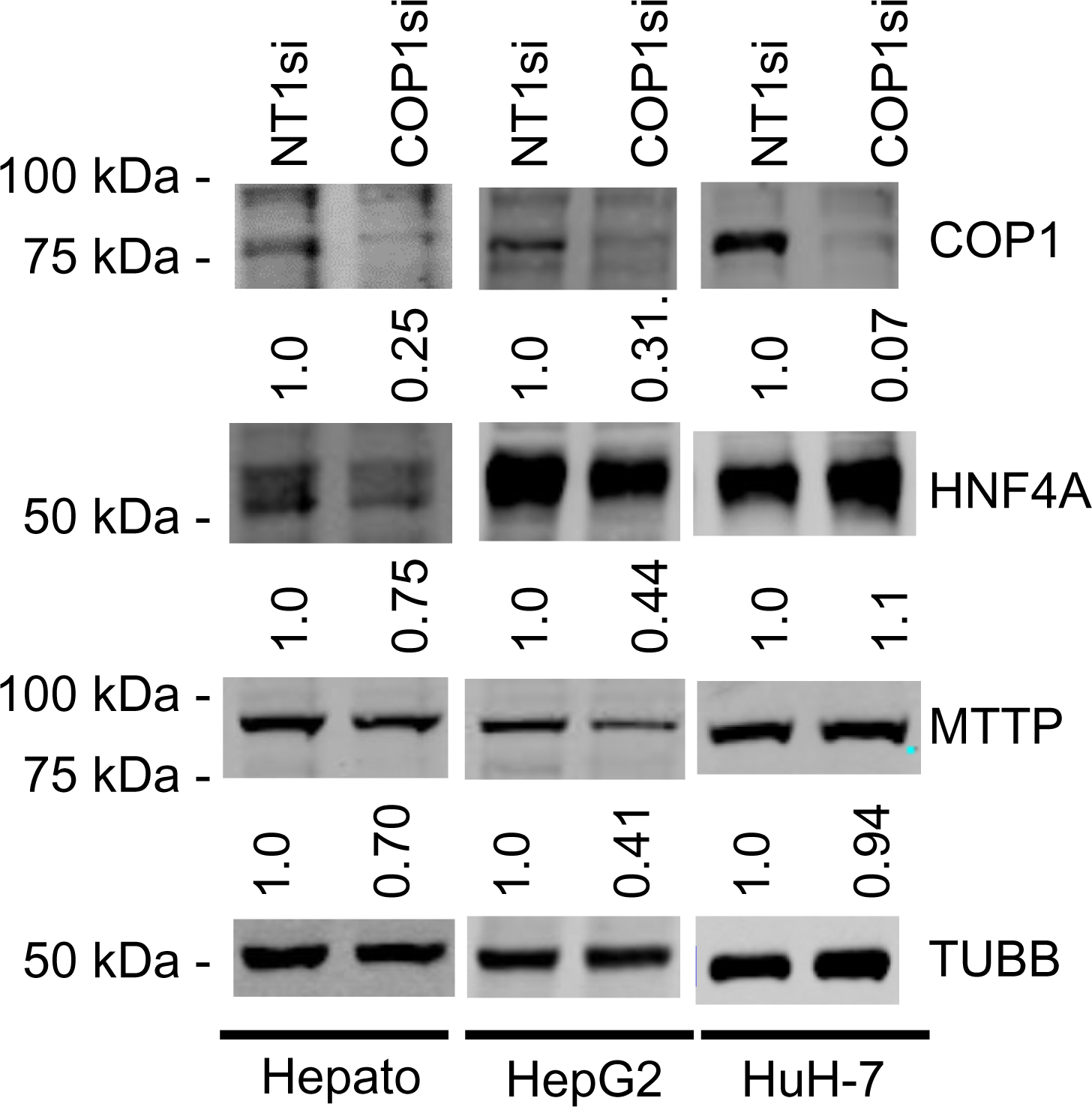
Western blots of COP1si treated cells. Primary hepatocytes (Hepato), HepG2 cells and HuH-7 cells were treated with COP1si or a Non-target control (NT1) for 96 h. Western blot is representative of 2 independent biological repeats for primary hepatocytes, and 4 for HepG2 and HuH-7 cells. In each cell type, the same blot was probed sequentially (COP1, MTTP, HNF4A, TUBB). Quantification is shown after correcting for TUBB (Tubulin beta chain) intensity and normalizing to the corresponding NT1si value.

### Gene Set Enrichment Analysis analyses of transcripts in COP1-suppressed transformed and primary hepatocytes

Transcriptome changes were then submitted to Gene Set Enrichment Analysis (GSEA). In contrast to ORA, GSEA leverages a ranking approach rather than nominal significance of individual hits to map transcripts of interest to gene sets ^34^. Importantly, the analysis provides directionality through a normalized effect size estimate. GSEA analyses were consistent with ORA findings. Findings in primary hepatocytes and HepG2 cells centered on impaired metabolic processes whereas terms related to inhibition in DNA replication and cell cycling were identified in HuH-7 (Table S5). In contrast to HuH-7 cells, transcripts mapping to DNA replication and proliferation were globally increased in HepG2 cells.

### Ingenuity pathway analysis points to HNF4A dysfunction in primary hepatocytes and HepG2 cells following COP1 knock-down

To further delineate the role of COP1, we then performed Ingenuity pathway analysis (IPA). Like GSEA, IPA leverages changes in direction and magnitude to infer functionality but it uses its own set of curated pathways (canonical pathways) and a different analysis pipeline. Mapping transcriptome changes to canonical pathways following COP1si was largely supportive of earlier analyses (Fig S4; Fig S5; Table S6). In addition, IPA can infer effects on master regulators (“Upstream Regulators”) in line with transcriptome perturbations. The analysis identified several targets of interest (Table S7). Changes in line with reduced HNF4A activity (“HNF4A”, “HNF4alpha dimer”) and HNF1A/B, two master regulators of liver function, were observed in HepG2 and primary hepatocytes but not in HuH-7 cells. Upon verification of the transcriptome-wide changes in the GSE206116 dataset, *HNF4A* abundance was reduced 2.04-fold in HepG2 (p = 0.0001) and trended towards a reduction (1.28-fold (p=0.11)) in hepatocytes following COP1 silencing. By contrast, HNF4A was possibly increased (1.24-fold, p=0.36) in HuH-7 cells.

### COP1 suppression results in reduced HNF4A and MTTP protein abundance in HepG2 cells and primary hepatocytes

Western blot analyses confirmed that reduced *HNF4A* transcript abundance was accompanied by lower protein abundance in both primary hepatocytes, HepG2 but not in HuH-7 cells (Figure 2). *MTTP* is a major regulator of very low density lipoprotein assembly and its expression requires HNF4A. Consistently, *COP1* silencing resulted in a statistically significant reduction of *MTTP* mRNA in both HepG2 and primary hepatocytes (3.5-fold and 1.69-fold, respectively, in GSE206116). Again, effects in HuH-7 cells differed with a modest upregulation (1.2-fold). Western blot analysis confirmed that MTTP protein abundance was concordantly reduced in hepatocytes and HepG2 cells, but unchanged in HuH-7 cells (Figure 2).

To ensure specificity, COP1 suppression was repeated in a distinct hepatocyte lot, using the other two siRNAs used earlier (si33 and si91). In addition to *HNF4A* and *MTTP*, *CEBPA,* another major liver transcription factor, reduced in GSE206116 (-1.8-fold, p = 7.8e-5), was also measured. COP1 suppression resulted in a trend toward reduced *HNF4A* expression, reduced CEBPA and MTTP abundance, replicating the earlier findings (Fig S6A). Importantly, compromised HNF4A function was ascertained by a reduction of Apolipoprotein B-100 abundance in the media in silenced cells (Fig S6B).

### Limited impact of higher COP1 levels on HepG2 cells

The consequences of increased *COP1* expression were examined next by transducing HepG2 cells with *COP1*. Silent mutations and a FLAG tag were introduced to render the transcript siRNA-resistant and to facilitate downstream experiments. *TRIB1*, which was upregulated in the transcriptome analysis and plays a role in sustaining HNF4A function was also included ^35,36^. Except for COP1 itself, moderate *COP1* overexpression did not alter transcripts tested, although *TRIB1* and *MTTP* were slightly increased (Fig S7) with no effects on protein levels other than that of COP1.

### Introduction of a siRNA resistant COP1 form partially normalizes transcript abundance

Since COP1 overexpression had limited downstream effects, we next verified that the siRNA-elicited changes were specific to COP1. COP1 silencing was repeated in pools of HepG2 cells expressing a siRNA resistant COP1. Introduction of the siRNA resistant COP1 (resulting in a 2-3-fold increase over basal level) provided a partial protection as assessed by a return towards baseline of *HNF4A, MTTP, APOC3 (*another prominent target of HFN4A) and*TRIB1* (Fig 3A, B). Changes at the protein level were confirmed by western blot for HNF4A, MTTP and COP1, for which suitable antibodies were available (Fig 3 C).

**Figure 3.**
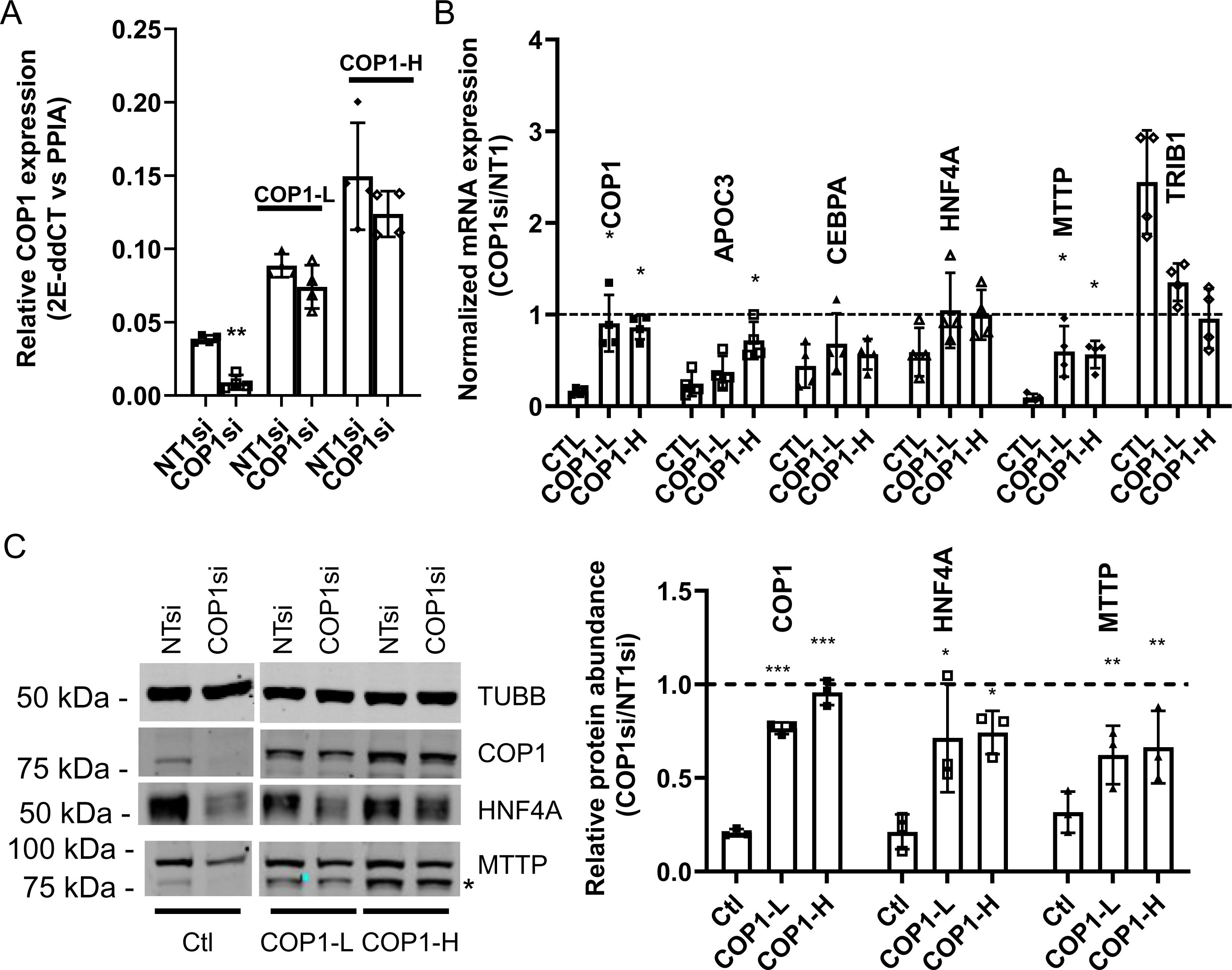
A siRNA resistant COP1 attenuates the impact of COP1si on hepatic markers. Pools of HepG2 cells, stably transduced with 2 doses (COP1-L(ow) and COP1-H(igh)) of a siRNA resistant form of (flag)COP1 or empty virus (viral dose matching the Wt-H; Ctl) were treated with COP1si or a control siRNA (NT1). A, relative expression of COP1 in transduced cells. Levels of *COP1* transcript were measured in COP1or NT1 silenced cells. For each transfection, statistical significance (NT vs COP1si) was measured using a Student’s paired t-tests. B, qPCR analyses of hepatic markers. Results shown are the COP1si values relative to the corresponding NT1 values. Statistical significance of the COP1 restoration was tested by 1-way repeated ANOVA followed by a *post-hoc* Dunnett’s test with the (PLVX) Ctl. Error bars represent standard deviations. C, representative Western blot analysis (left) and quantification of 3 biological repeats (right). Blot was probed sequentially with COP1 and MTTP; leftover COP1 signal is indicated by *. TUBB, Tubulin beta chain. To test for statistical significance, repeated measures 1-way ANOVAs on the relative (COP1/NT1) protein abundance values (followed by a *post-hoc* Dunnett’s comparison test with the control) were performed. Values shown correspond to biological replicates. Bars represent averages ± SD. Errors: *, p<0.05; **,p<0.01;***,<0.001. Bars represent the SD.

### HNF4A silencing partially phenocopies COP1 suppression in HepG2 cells

COP1 suppression reduced HNF4A protein levels in HepG2 and primary hepatocytes. Since HNF4A is a master regulator of hepatocyte function we hypothesized that the impacts of COP1 suppression might be through impaired *HNF4A* function ^37,38^. First, we confirmed that *HNF4A* suppression via siRNA mediated silencing phenocopies *COP1* suppression in HepG2 cells (Fig 4A). *HNF4A* suppression resulted in reduced *CEBPA*, *MTTP* and *APOC3* but increased *TRIB1*. Interestingly, there was evidence of a dose dependent relationship, with greater suppression leading to more pronounced effects. Changes in *HNF4A*, *COP1* and *MTTP* were further investigated and validated by western blot (Fig 4B). Unlike *COP1* suppression, which led to diminished *HNF4A* expression in HepG2 cells, *HNF4A* suppression had a modest impact on *COP1*, suggesting a unidirectional relationship whereby *COP1* regulates *HNF4A* function.

**Figure 4.**
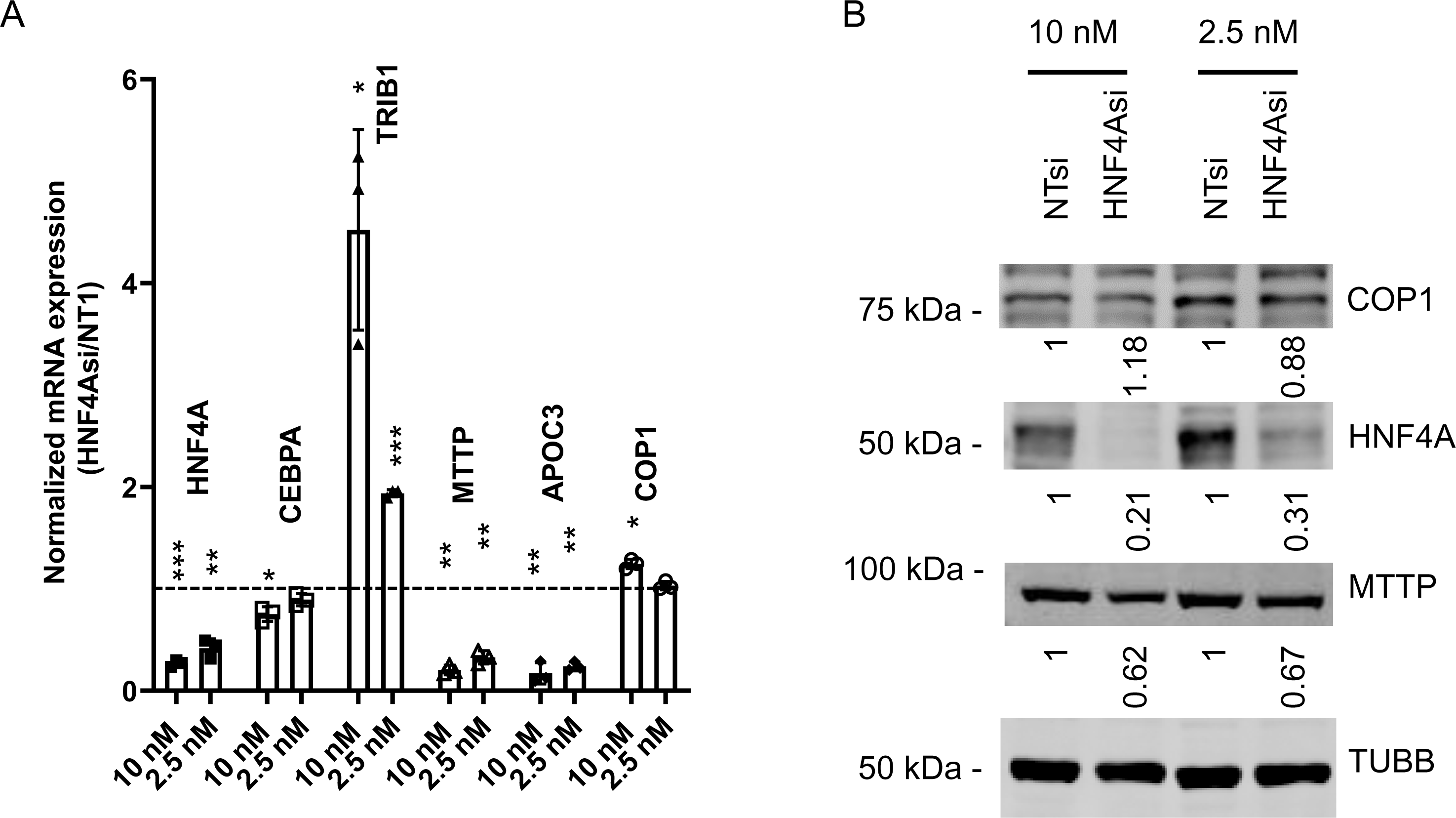
HNF4A suppression phenocopies COP1 suppression in HepG2 cells. HepG2 cells were transfected with 10 nM or 2.5 nM siRNA targeting HNF4A or the non-target control, NT1) for 96 h prior to harvest for RNA or protein. A, RT-PCR analysis of transcripts. Values were normalized internally to PPIA levels and are expressed relative to the corresponding NT1 values. Results from three biological replicates (average and SD) are shown. Statistical significance was tested for 3 experimental repeats against the NT1 value using Student’s t-test. *p<0.05, ** p< 0.01, *** p<0.001. B, Western blot analysis of corresponding samples. Values were corrected for TUBB (Tubulin beta chain) intensity and normalized to the matching NT1 control. Results shown are representative of 3 experimental repeats.

The consequences of HNF4A overexpression were tested next. The liver enriched HNF4A form, driven by the P1 promoter (corresponding to NP_000448.3), was transduced in naïve HepG2 cells and the contribution of increased HFN4A to cognate gene expression was examined. HNF4A overexpression (∼ 2-fold at the protein level) was sufficient to increase *MTTP* and *APOC3* abundance. By contrast, effects on *CEBPA*, *COP1* and *TRIB1* were modest, although reaching nominal statistical significance for *CEBPA* and *TRIB1* (Fig S8). These findings are consistent with the presence of at least 2 distinct COP1 regulatory axes in HepG2 cells: a COP1-HNF4A axis, regulating *APOC3* and *MTTP* expression, and the presence of another COP1-dependent but HNF4A independent (or less dependent) regulatory axes (*CEBPA*, *TRIB1*).

### HNF4A overexpression restores MTTP and APOC3 expression but not CEBPA in COP1 silenced HepG2 cells

Next, we determined the extent to which impaired HNF4A function contributed to the changes observed following COP1 suppression. The lack of impact of COP1 overexpression on HNF4A protein levels suggested that changes in HNF4A protein abundance in COP1 suppressed cells might stem from reduced *HNF4A* transcription rather than changes at the post-transcriptional level. We predicted that exogenously introduced HNF4A, driven by a heterologous promoter, might resist COP1 silencing and thus allow the contribution of HNF4A under COP1 suppressed conditions to be assessed. To test this hypothesis, *HNF4A* was stably transduced in HepG2 cells. This resulted in a ∼60% increase in *HNF4A* mRNA and ∼80% increase in HNF4A protein level (Fig 5). Importantly, at comparable *HNF4A* levels (i.e. [HNF4A OE + COP1si] vs [PLVX + NT1si]), *APOC3* and *MTTP* expression levels were similar, confirming that changes secondary to COP1si can be ascribed to reduced HNF4A abundance. By contrast, HNF4A transduction failed to normalize *CEBPA* and *TRIB1* abundance. Thus, COP1 regulates *TRIB1* and *CEPBA* in part via HNF4A-independent processes.

**Figure 5.**
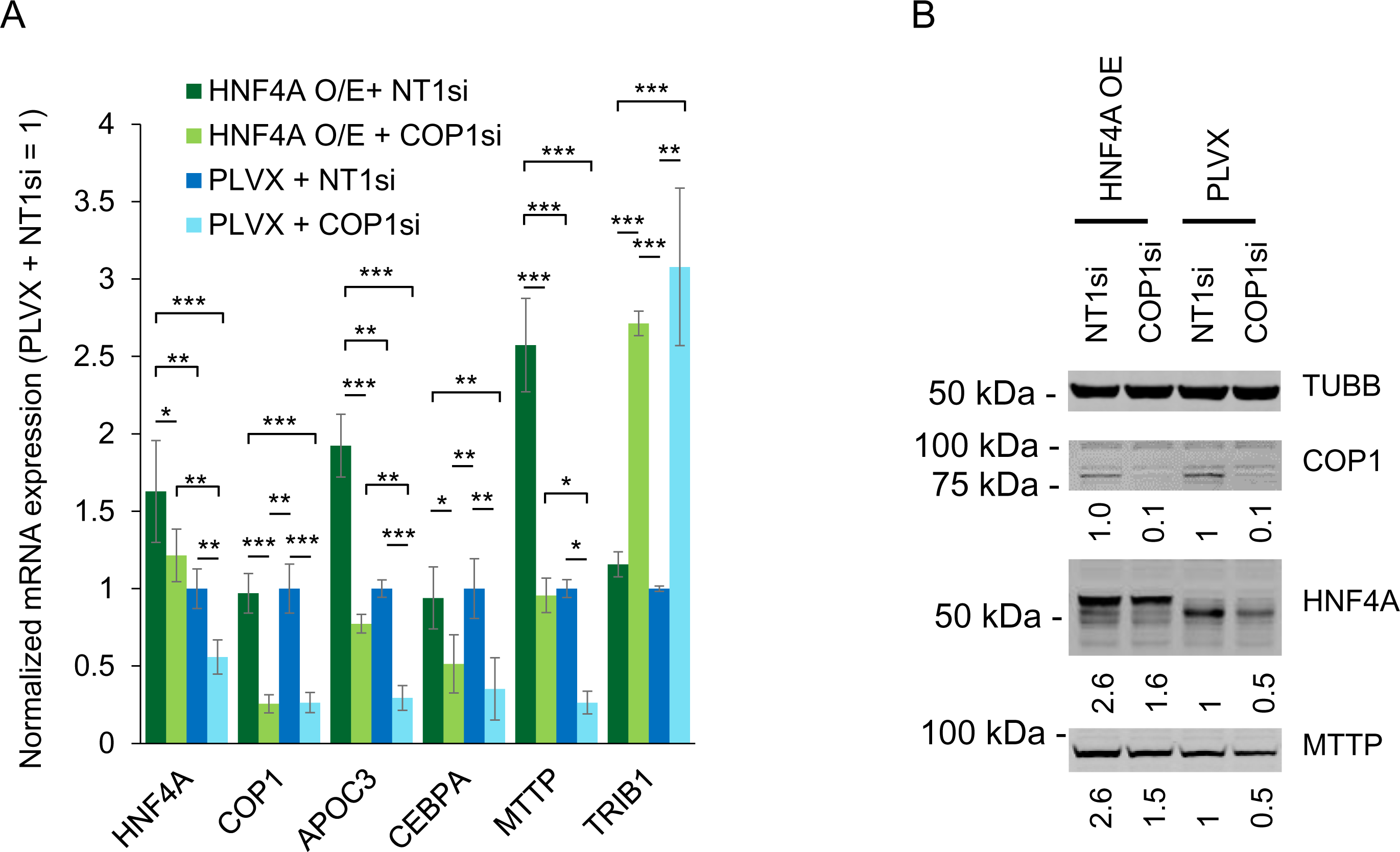
COP1 suppression in HepG2 cells can be partially compensated by HNF4A overexpression. Impact of HFN4A overexpression (O/E) on a selection of targets was assessed by qRT-PCR and Western botting. A, Transcript abundance assessed qRT-PCR. Values were internally normalized to PPIA expression and are shown relative to the matching mean [PLVX + NT1si] value. Bars represent the means of 3 independent biological repeats (± S.D.). Statistical significance was tested for each transcript using a repeated measures ANOVA followed by Tukey’s *post-hoc* tests. Only statistically significant differences are shown. *p<0.05, ** p< 0.01, *** p<0.001. B, Western blot representative of 3 biological repeats. Intensities were normalized to TUBB (Tubulin beta chain) and expressed relative to the [PLVX + NT1si] control.

### Identification of pathways co-regulated by *HNF4A* and *COP1*

To estimate the degree to which processes were co-regulated by HNF4A and COP1, expression profiles of COP1 and HNF4A suppressed cells were compared. Data from GSE15991, reporting transcriptome changes in *HNF4A* silenced HepG2, were extracted and compared with our COP1 silenced findings ^39^. *HNF4A* suppression led to perturbations of about a third of the gene transcripts. A comparison of the FDR significant transcripts affected by HNF4A and COP1 knockdowns revealed a common set of 215 enriched transcripts (out of 4785 and 515 unique gene symbols for HNF4Asi and COP1si samples, respectively). While this reveals that a large proportion of COP1 silenced transcripts (215/515) are also HNF4A targets, it also indicates that the majority of the HNF4A dependent transcripts are not regulated by COP1. Over-representation analyses of the overlapping transcripts identified 62 FDR significant Gene Ontology terms, ranging from inflammation to steroid metabolism and prioritized Reactome and KEGG gene sets linked to bile acid metabolism, peroxisome function and complement systems (Table 2, Table S8). These results suggest that HNF4A and COP1 jointly regulate a wide range of biological pathways central to liver function.

**Table 2.**
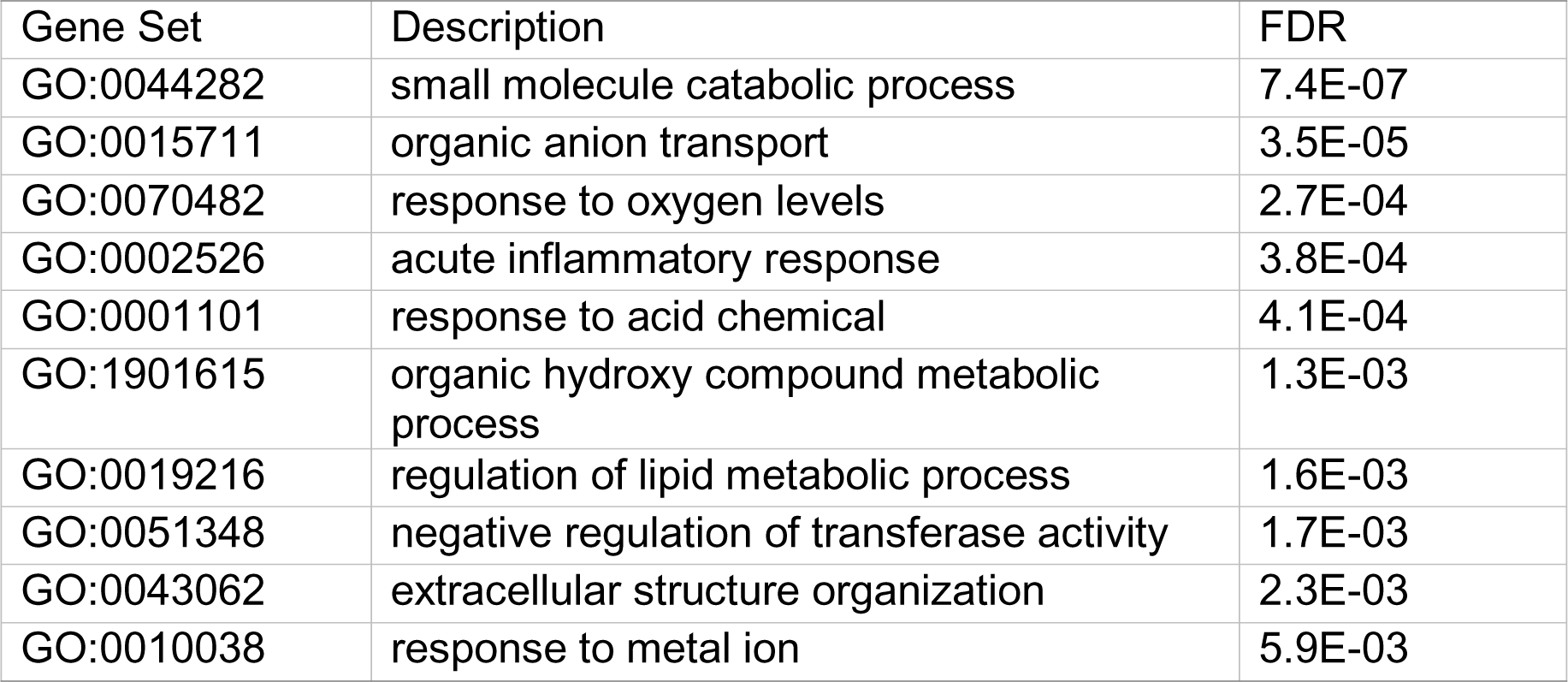
Over-representation analysis of (FDR) significant transcripts following HNF4A and COP1 silencing. Enriched Gene Ontology terms (Biological_Process_noRedundant) are shown, after simplification by Weighted Set Cover (Webgestalt). A complete table, including KEGG and Reactome sets, is included in Supplementary Tables (Table S7).

| Gene Set | Description | FDR |
| --- | --- | --- |
| GO:0044282 | small molecule catabolic process | 7.4E-07 |
| GO:0015711 | organic anion transport | 3.5E-05 |
| GO:0070482 | response to oxygen levels | 2.7E-04 |
| GO:0002526 | acute inflammatory response | 3.8E-04 |
| GO:0001101 | response to acid chemical | 4.1E-04 |
| GO:1901615 | organic hydroxy compound metabolic process | 1.3E-03 |
| GO:0019216 | regulation of lipid metabolic process | 1.6E-03 |
| GO:0051348 | negative regulation of transferase activity | 1.7E-03 |
| GO:0043062 | extracellular structure organization | 2.3E-03 |
| GO:0010038 | response to metal ion | 5.9E-03 |

## Discussion

Here we have explored COP1 function in both normal and transformed hepatocytes. Our transcriptomics findings in HepG2 and HuH-7 differ considerably from those previously published ^15^. In that report, 7 transcripts linked to p53 (including RFWD2/COP1) and a list of 78 genes that exhibited directionally coherent changes were reported to be impacted by COP1 silencing. No such pattern was evident from our data (Fig S9, Table S9). Reasons for this discrepancy are unclear. One possibility is that the previously reported changes resulted from apoptosis engagement, possibly through p53. Indeed, a separate report demonstrated (albeit in non-liver models) that COP1 could mediate p53 turnover, preventing p53 mediated cell death ^7^. Thus, COP1 downregulation combined with other stressors, rather than COP1 suppression *per se*, may activate p53 and lead to apoptotic death. By contrast, we did not find evidence that reduced COP1 level is cytotoxic in either cell model, consistent with lack, or insufficiency, of p53 engagement.

*COP1* silencing had significant but dichotomous effects on HepG2 and HuH-7. Whereas changes elicited by *COP1* suppression matched a prominently proliferative profile in HuH-7, perturbations in HepG2 revealed a metabolic portrait reminiscent of primary hepatocytes. Importantly, this work does not answer whether these different requirements for COP1 can be attributed to cell line singularities (e.g., specific mutations) or broader features (e.g., hepatoblastoma vs hepatocarcinoma). One limitation to these findings is that this 3-way transcriptome-wide comparison hinges on a single siRNA type, albeit designed to minimize off-targeting. While two additional siRNAs had similar impacts on a subset of targets of relevance to hepatocyte function, their transcriptome scale impacts were not investigated, and the extent of off-targeting remains unknown. However, it seems unlikely that off-targeting could account for the lack of cytotoxic impact of COP1 silencing observed here or the contrasted transcriptomic signatures.

Interestingly, transcripts could be mapped to the same terms and pathways despite HepG2 cells and hepatocytes sharing only a subset (∼8 %) of nominally impacted transcripts. Most were directionally consistent by GSEA (e.g., cellular ketone metabolic processes, (reduced) lipid homeostasis, (reduced) small molecule catabolic process (reduced) etc..), although exceptions were evident. For instance, IPA indicated that COP1 suppression had opposite effects on transcripts linked to cholesterol synthesis (e.g., “Superpathway of Cholesterol Biosynthesis”) in HepG2 and hepatocytes. Uniquely, COP1 suppression in hepatocytes resulted in increased abundance of transcripts that were mapped by ORA to gene sets linked to inflammation. Although how it contributes to inflammation is still unclear, several mediators of liver inflammation featured prominently, including IL1B (1.9-fold increase) and IL1R1 (1.6-fold increase), suggesting that COP1 plays an anti-inflammatory role. This is supported by IPA which predicts an activation of IL1 (IL1, IL1A and IL1B) signaling following COP1 knock-down (Table S6).

Leveraging siRNA experiments in HepG2 cells, HNF4A was identified as an important effector regulating a subset of COP1 targets. Importantly, COP1 restoration using a siRNA resistant expression construct could largely normalize HNF4A levels in HepG2 cells, ensuring siRNA specificity. Although restoration experiments were not performed in primary hepatocytes, our siRNA results indicated that COP1 is also instrumental in supporting HNF4A function therein. Comparison of FDR significant hits in HNF4Asi and COP1si samples identified unique and shared gene targets. Transcripts impacted by COP1 silencing represented a small subset (215/4785) of HNF4A responsive genes while a large fraction (215/515) of COP1si impacted genes were also sensitive to HNF4A silencing. This analysis was conservative, as several transcripts of potential biological importance were excluded because they did not reach FDR significance. Indeed, as the extent of HNF4A reduction associated with COP1 knock-down is less pronounced than that achieved via a targeted HNF4A approach, sufficient HNF4A may persist in COP1 silenced cells to sustain near normal or not FDR significant changes in the expression of target genes. A more permissive approach using nominally significant hits (not corrected for the number of targets investigated) suggests that 20-25% of the transcriptome requires both COP1 and HNF4A (*data not shown*).

In view of the essential role of HNF4A in hepatocyte function, future work will aim to clarify how COP1 increases HNF4A levels ^40,41^. The COP1-HNF4A relationship appears largely unidirectional as COP1 expression was unaffected by HNF4A targeting. Whether HNF4A can affect COP1 function in other ways remains untested. COP1 is an E3 ligase that targets cognate proteins for degradation and although HNF4A can be ubiquitylated, our data are inconsistent with COP1 directly promoting HNF4A degradation ^42,43^. For one, COP1 suppression led to decreased, rather than increased, HNF4A protein abundance. Moreover, COP1 overexpression did not reduce HNF4A protein level. Thus, the physical interaction we previously reported between HNF4A and TRIB1, a COP1 adaptor protein involved in CEBPA degradation, is unlikely to contribute to HNF4A degradation; indeed, TRIB1 was positively correlated with HNF4A abundance in that study ^36^. Rather, COP1 may promote the degradation or inactivation of HNF4A transcription repressors, such as SNAI1 or NF-kappaB ^44–46^. Of note, GSEA found NF-kappaB signaling to be increased in COP1 suppressed HepG2 cells (Table S5). Furthermore, changes consistent with increased “TNF” (i.e., TNFA) activity were identified by IPA in HepG2 and hepatocytes, consistent with activation of NF-kappaB signaling (Table S7). Future studies will examine this possibility.

In summary, we demonstrated that HepG2 and HuH-7 could proliferate overtly normally despite reduced COP1 expression. Moreover, COP1 was required to maintain prominent hepatocyte transcripts, including HNF4A, in primary hepatocytes and HepG2 cells, but not in HuH-7 cells. Thus, HepG2 cells represent a better model system to further clarify the role of COP1 in liver physiology.

## Acknowledgments

This work was funded by a Canadian Institutes of Health Research Foundation grant (FRN:154308; RM).

## Author contributions

S.S. designed, performed, and analyzed experiments. P.L. performed experiments.

R.M. and S.S. wrote the manuscript.

## Data availability

Expression datasets are deposited at the Gene Expression Omnibus (https://www.ncbi.nlm.nih.gov/geo/): Our COP1 suppression (HepG2, HuH-7 and Primary hepatocytes) data corresponds to GSE206116 whereas the HNF4A suppression dataset is available as GSE15991.

## Abbreviations

COP1: Constitutive Photomorphogenesis Protein 1 homolog
HNF4A: hepatic nuclear factor-4 alpha
GSEA: Gene Set Enrichment Analysis
KEGG: Kyoto Encyclopedia of Genes and Genomes
GO: Gene Ontology
ORA: over-representation analysis
IPA: Ingenuity Pathway Analysis.

## References

1. Zheng, N. & Shabek, N. Ubiquitin Ligases: Structure, Function, and Regulation. Annu. Rev. Biochem. 86, 129–157 (2017).

2. Ouyang, W. et al. Erk1/2 inactivation promotes a rapid redistribution of COP1 and degradation of COP1 substrates. Proc. Natl. Acad. Sci. U. S. A. 117, 4078–4087 (2020).

3. Vitari, A. C. et al. COP1 is a tumour suppressor that causes degradation of ETS transcription factors. Nature 474, 403–408 (2011).

4. Bianchi, E. et al. Characterization of human constitutive photomorphogenesis protein 1, a RING finger ubiquitin ligase that interacts with Jun transcription factors and modulates their transcriptional activity. J. Biol. Chem. 278, 19682– 19690 (2003).

5. Migliorini, D. et al. Cop1 constitutively regulates c-Jun protein stability and functions as a tumor suppressor in mice. J. Clin. Invest. 121, 1329–1343 (2011).

6. Qi, L. et al. TRB3 links the E3 ubiquitin ligase COP1 to lipid metabolism. Science 312, 1763–1766 (2006).

7. Dornan, D. et al. The ubiquitin ligase COP1 is a critical negative regulator of p53. Nature 429, 86–92 (2004).

8. Prudente, S. et al. Trib1 and Evi1 cooperate with Hoxa and Meis1 in myeloid leukemogenesis. Blood 282, 122–126 (2013).

9. Choi, H. H. et al. COP9 signalosome subunit 6 stabilizes COP1, which functions as an E3 ubiquitin ligase for 14-3-3σ. Oncogene 2011 3048 30, 4791–4801 (2011).

10. Kato, S., Ding, J., Pisck, E., Jhala, U. S. & Du, K. COP1 functions as a FoxO1 ubiquitin E3 ligase to regulate FoxO1-mediated gene expression. J. Biol. Chem. 283, 35464–35473 (2008).

11. Song, Y. et al. Role of the COP1 protein in cancer development and therapy. Semin. Cancer Biol. 67, 43–52 (2020).

12. Bauer, R. C. et al. Tribbles-1 regulates hepatic lipogenesis through posttranscriptional regulation of C/EBPα. J. Clin. Invest. 125, 3809–3818 (2015).

13. Yoshida, A., Kato, J. Y., Nakamae, I. & Yoneda-Kato, N. COP1 targets C/EBPalpha for degradation and induces acute myeloid leukemia via Trib1. Blood 122, 1750–1760 (2013).

14. Sawada, G. et al. Loss of COP1 expression determines poor prognosis in patients with gastric cancer. Oncol. Rep. 30, 1971–1975 (2013).

15. Lee, Y. H. et al. Definition of ubiquitination modulator COP1 as a novel therapeutic target in human hepatocellular carcinoma. Cancer Res. 70, 8264–8269 (2010).

16. Sanchez-Barcelo, E. J., Mediavilla, M. D., Vriend, J. & Reiter, R. J. Constitutive photomorphogenesis protein 1 (COP1) and COP9 signalosome, evolutionarily conserved photomorphogenic proteins as possible targets of melatonin. J. Pineal Res. 61, 41–51 (2016).

17. Nakabayashi, H., Taketa, K., Miyano, K., Yamane, T. & Sato, J. Growth of human hepatoma cells lines with differentiated functions in chemically defined medium. Cancer Res. 42, 3858–3863 (1982).

18. Aden, D. P., Fogel, A., Plotkin, S., Damjanov, I. & Knowles, B. B. Controlled synthesis of HBsAg in a differentiated human liver carcinoma-derived cell line. Nature 282, 615–616 (1979).

19. López-Terrada, D., Cheung, S. W., Finegold, M. J. & Knowles, B. B. Hep G2 is a hepatoblastoma-derived cell line. Hum. Pathol. 40, 1512–1515 (2009).

20. Ranganathan, S., Lopez-Terrada, D. & Alaggio, R. Hepatoblastoma and Pediatric Hepatocellular Carcinoma: An Update. Pediatr. Dev. Pathol. 23, 79–95 (2020).

21. Kasai, F., Hirayama, N., Ozawa, M., Satoh, · Motonobu & Kohara, A. HuH-7 reference genome profile: complex karyotype composed of massive loss of heterozygosity. Hum. Cell 31, 261–267 (2018).

22. Zhou, B. et al. Haplotype-resolved and integrated genome analysis of the cancer cell line HepG2. Nucleic Acids Res. 47, 3846–3861 (2019).

23. Arzumanian, V. A., Kiseleva, O. I. & Poverennaya, E. V. The curious case of the HepG2 cell line: 40 years of expertise. Int. J. Mol. Sci. 22, 13135 (2021).

24. Zhao, Y. et al. Genomic alterations across six hepatocellular carcinoma cell lines by panel-based sequencing. Transl. Cancer Res. 7, 231–239 (2018).

25. Meex, S. J., Andreo, U., Sparks, J. D. & Fisher, E. A. Huh-7 or HepG2 cells: which is the better model for studying human apolipoprotein-B100 assembly and secretion? 131. J.Lipid Res. 52, 152–158 (2011).

26. Tsai, J., Qiu, W., Kohen-Avramoglu, R. & Adeli, K. MEK-ERK inhibition corrects the defect in VLDL assembly in HepG2 cells: potential role of ERK in VLDL-ApoB100 particle assembly. Arterioscler.Thromb.Vasc.Biol. 27, 211–218 (2007).

27. Gunn, P. J., Green, C. J., Pramfalk, C. & Hodson, L. In vitro cellular models of human hepatic fatty acid metabolism: differences between Huh7 and HepG2 cell lines in human and fetal bovine culturing serum. Physiol. Rep. 5, (2017).

28. Berger, E., Vega, N., Weiss-Gayet, M. & Géloën, A. Gene Network Analysis of Glucose Linked Signaling Pathways and Their Role in Human Hepatocellular Carcinoma Cell Growth and Survival in HuH7 and HepG2 Cell Lines. Biomed Res. Int. 2015, 821761 (2015).

29. Liao, Y., Wang, J., Jaehnig, E. J., Shi, Z. & Zhang, B. WebGestalt 2019: gene set analysis toolkit with revamped UIs and APIs. Nucleic Acids Res. 47, W199–W205 (2019).

30. The Gene Ontology Consortium et al. The Gene Ontology knowledgebase in 2023. Genetics 224, iyad031 (2023).

31. Kanehisa, M. & Goto, S. KEGG: kyoto encyclopedia of genes and genomes. Nucleic Acids Res. 28, 27–30 (2000).

32. Kanehisa, M., Furumichi, M., Sato, Y., Kawashima, M. & Ishiguro-Watanabe, M. KEGG for taxonomy-based analysis of pathways and genomes. Nucleic Acids Res. 51, D587–D592 (2023).

33. Fabregat, A. et al. The Reactome Pathway Knowledgebase. Nucleic Acids Res. 46, D649–D655 (2017).

34. Subramanian, A. et al. Gene set enrichment analysis: a knowledge-based approach for interpreting genome-wide expression profiles. Proc. Natl. Acad. Sci. U. S. A. 102, 15545–50 (2005).

35. Soubeyrand, S., Martinuk, A., Naing, T., Lau, P. & McPherson, R. Role of Tribbles Pseudokinase 1 (TRIB1) in human hepatocyte metabolism. Biochim. Biophys. Acta 1862, 223–32 (2016).

36. Soubeyrand, S., Martinuk, A. & McPherson, R. TRIB1 is a positive regulator of hepatocyte nuclear factor 4-Alpha. Sci. Rep. 7, (2017).

37. Battle, M. A. et al. Hepatocyte nuclear factor 4alpha orchestrates expression of cell adhesion proteins during the epithelial transformation of the developing liver. Proc.Natl.Acad.Sci.U.S.A. 103, 8419–8424 (2006).

38. Hayhurst, G. P., Lee, Y. H., Lambert, G., Ward, J. M. & Gonzalez, F. J. Hepatocyte nuclear factor 4alpha (nuclear receptor 2A1) is essential for maintenance of hepatic gene expression and lipid homeostasis. Mol Cell Biol. 21, 1393–1403 (2001).

39. Wang, Z., Bishop, E. P. & Burke, P. A. Expression profile analysis of the inflammatory response regulated by hepatocyte nuclear factor 4α. BMC Genomics 12, 1–14 (2011).

40. Parviz, F. et al. Hepatocyte nuclear factor 4alpha controls the development of a hepatic epithelium and liver morphogenesis. Nat.Genet. 34, 292–296 (2003).

41. Santangelo, L. et al. The stable repression of mesenchymal program is required for hepatocyte identity: a novel role for hepatocyte nuclear factor 4alpha. Hepatology 53, 2063–2074 (2011).

42. Hong, Y. H., Varanasi, U. S., Yang, W. & Leff, T. AMP-activated protein kinase regulates HNF4α transcriptional activity by inhibiting dimer formation and decreasing protein stability. J. Biol. Chem. 278, 27495–27501 (2003).

43. Cai, W. Y. et al. Yes-associated protein/TEA domain family member and hepatocyte nuclear factor 4-alpha (HNF4α) repress reciprocally to regulate hepatocarcinogenesis in rats and mice. Hepatology 65, 1206–1221 (2017).

44. Simó, R. et al. Molecular Mechanism of TNFα-Induced Down-Regulation of SHBG Expression. Mol. Endocrinol. 26, 438–446 (2012).

45. Cicchini, C. et al. Snail controls differentiation of hepatocytes by repressing HNF4alpha expression. J Cell Physiol 209, 230–238 (2006).

46. Zhou, B. P. et al. Dual regulation of Snail by GSK-3beta-mediated phosphorylation in control of epithelial-mesenchymal transition. Nat. Cell Biol. 6, 931–940 (2004).

