## Supplementary figures for "Constitutive Photomorphogenesis Protein 1 homolog (COP1) sustains nuclear factor-4 alpha function in human hepatocyte models"

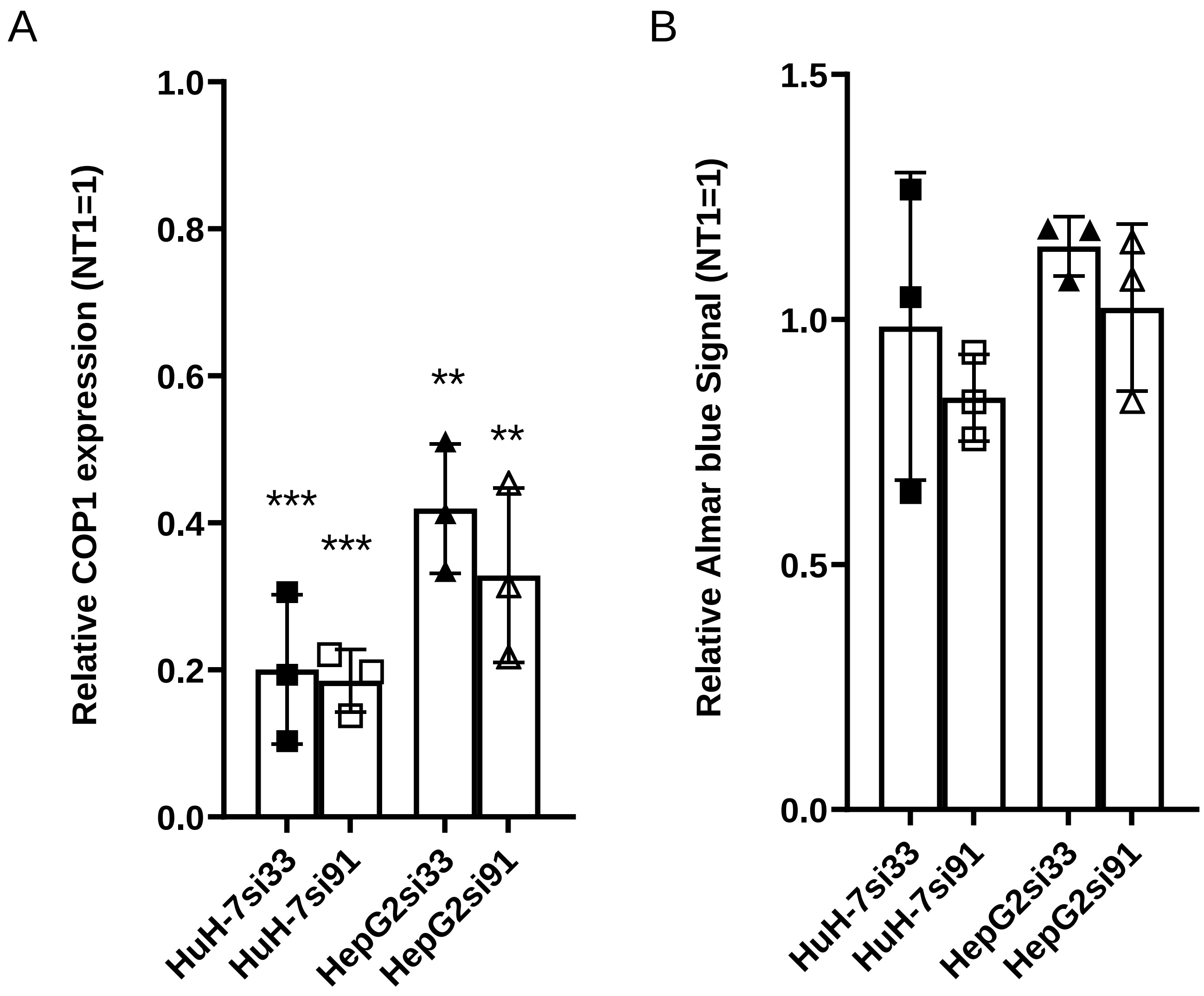


**Figure S1. COP1 suppression via distinct siRNA is associated with overtly normal cell growth in cell culture.** HepG2 and HuH-7 cells were treated with NT1, COP1si-33 or COP1si-91 for 96 h. Cell growth was estimated with Alamar blue (B) while mRNA was quantified by qRT-PCR (A). Results from three biological replicates (average and SD) are shown. Statistical significance of NT1 vs COP1 targeting siRNAs was tested using a Student's t-test. Errors: *, p<0.05; **,p<0.01;***,<0.001.

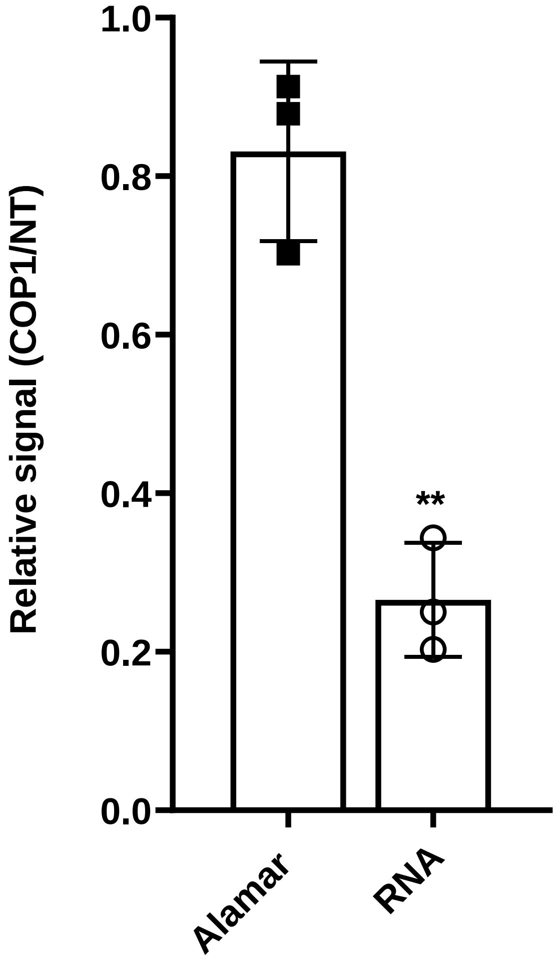


**Figure S2. Lipofectamine 2000 mediated COP1 suppression is not toxic in HuH-7 cells**. HuH-7 cells were treated for 96 h with COP1si and Lipofectamine 2000. Changes in proliferation (Alamar) and COP1 RNA levels (RNA) were then determined. Alamar signal reduction was not statistically significant (Student’s t-test). Results from three biological replicates (average and SD) are shown. **, $p<0.001$.

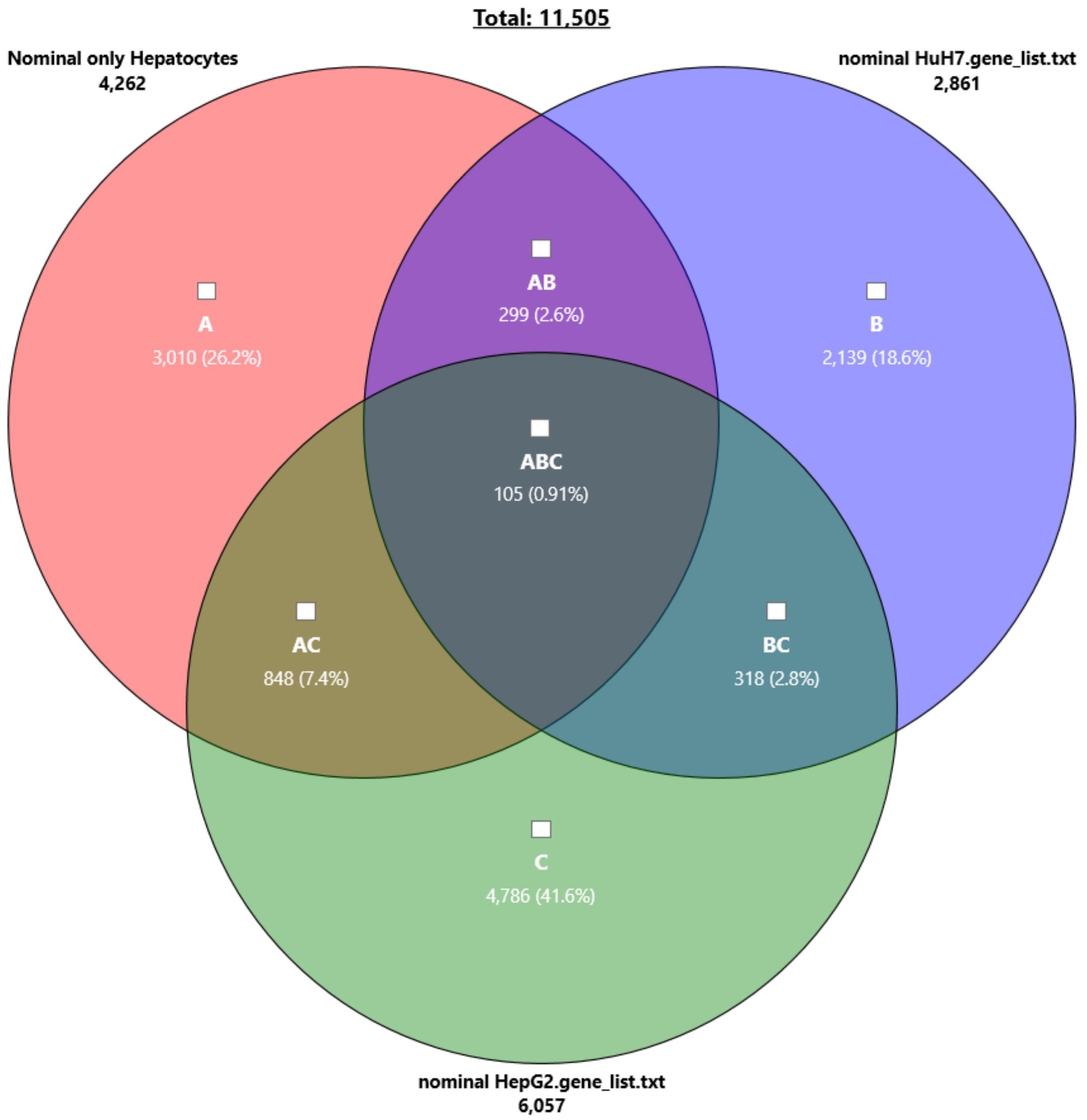


**Figure S3. Overlap of significantly changed transcripts in response to COP1 suppression in all 3 cell models**. Venn diagram representing overlap of nominally ($p < 0.05$) changed transcripts. Generated with the Transcriptome Analysis Console.

HepG2

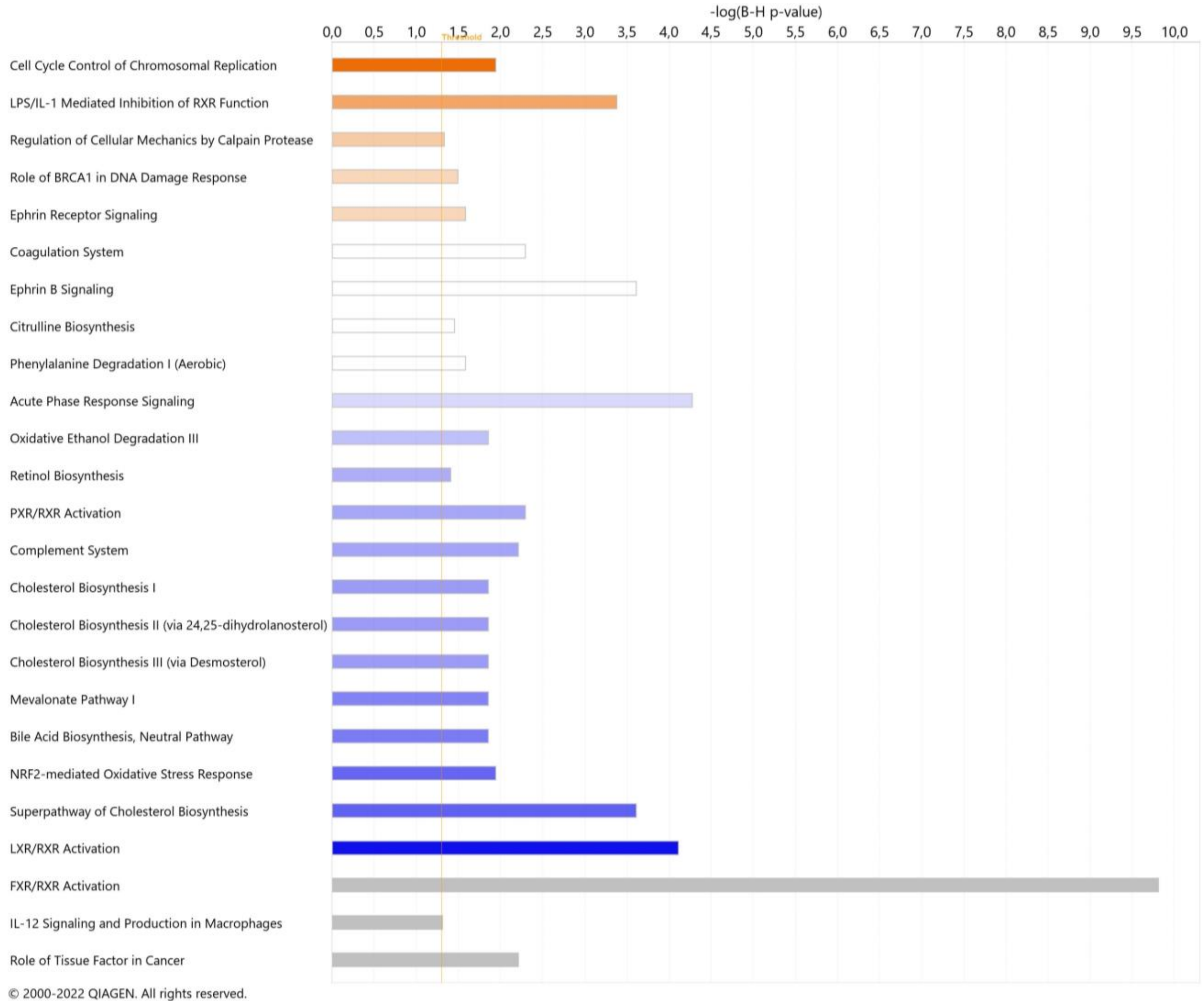


HuH-7

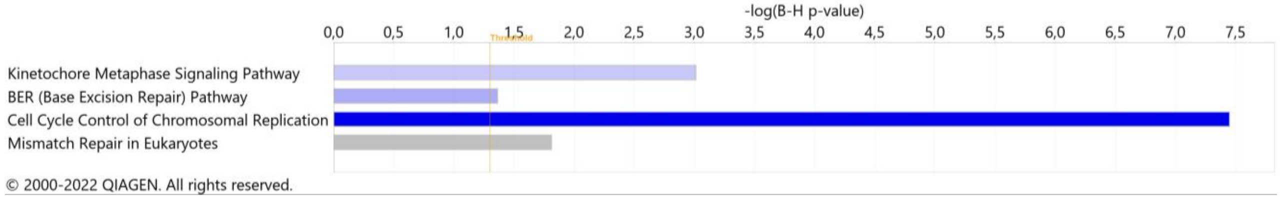


**Figure S4. Canonical pathways impacted by COP1si in HepG2 (left) and HuH-7 (right).** Organized by decreasing Z score (top to bottom). Only B-H significant hits are shown. Orange, positive Z-score (activation); blue, negative Z-score (inhibition); grey, direction uncertain. Darker color, higher Z-score (absolute value).

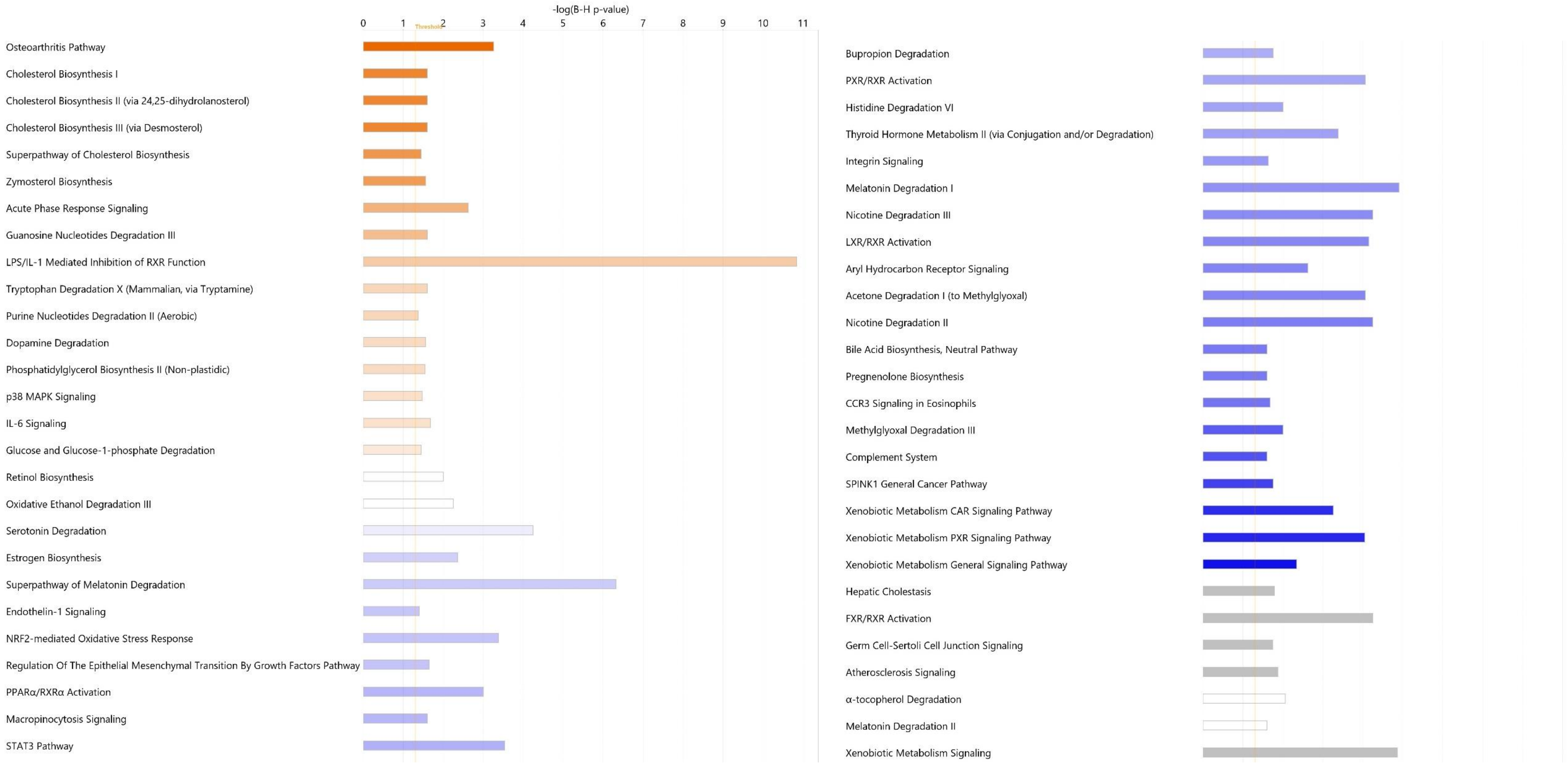


**Figure S5. Canonical pathways impacted by COP1si in hepatocytes.** Organized by decreasing Z score (left to right). Only BH significant hits are shown. Orange, positive Z-score (activation); blue, negative Z-score (inhibition); grey, direction uncertain. Darker color, higher Z-score (absolute value).

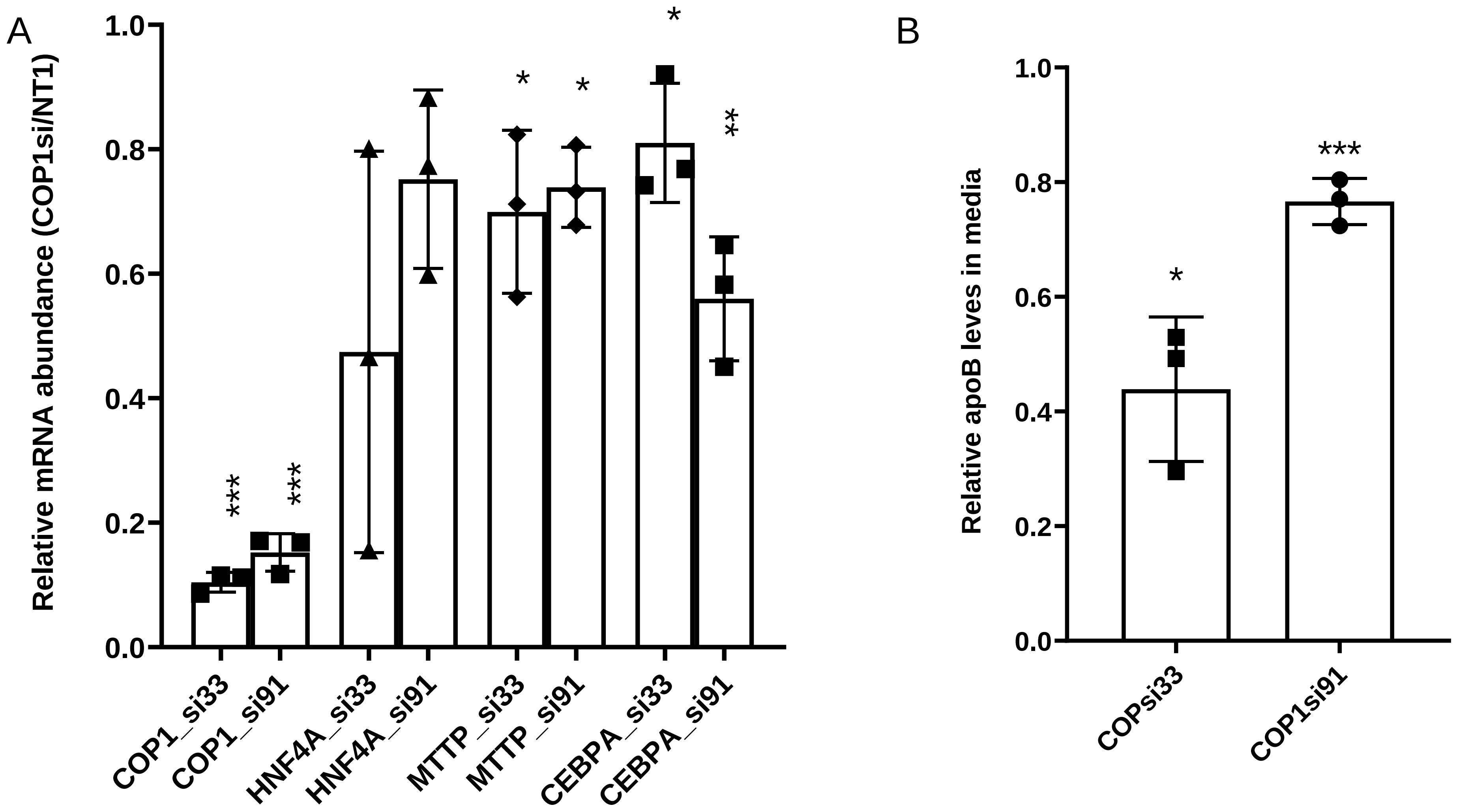


**Figure S6. COP1 suppression results in reduced expression of major liver genes and apoB secretion**. A, Hepatocytes were transfected with control or two cognate COP1 siRNAs (si33 and si91). RNA abundance was determined by qRT-PCR. Values were normalized internally to PPIA levels and are expressed relative to the corresponding NT1 values. B, Relative apolipoprotein B (apoB) concentration in the media. ApoB concentration was determined by ELISA, divided by the corresponding total RNA concentration (as a proxy of cell number). Values are expressed relative to the matching control (NT1) value. Results from three biological replicates (average and SD) are shown. Statistical significance was determined using a paired t-test vs the NT values. *, p<0.05;** p<0.01; ***,<0.001.

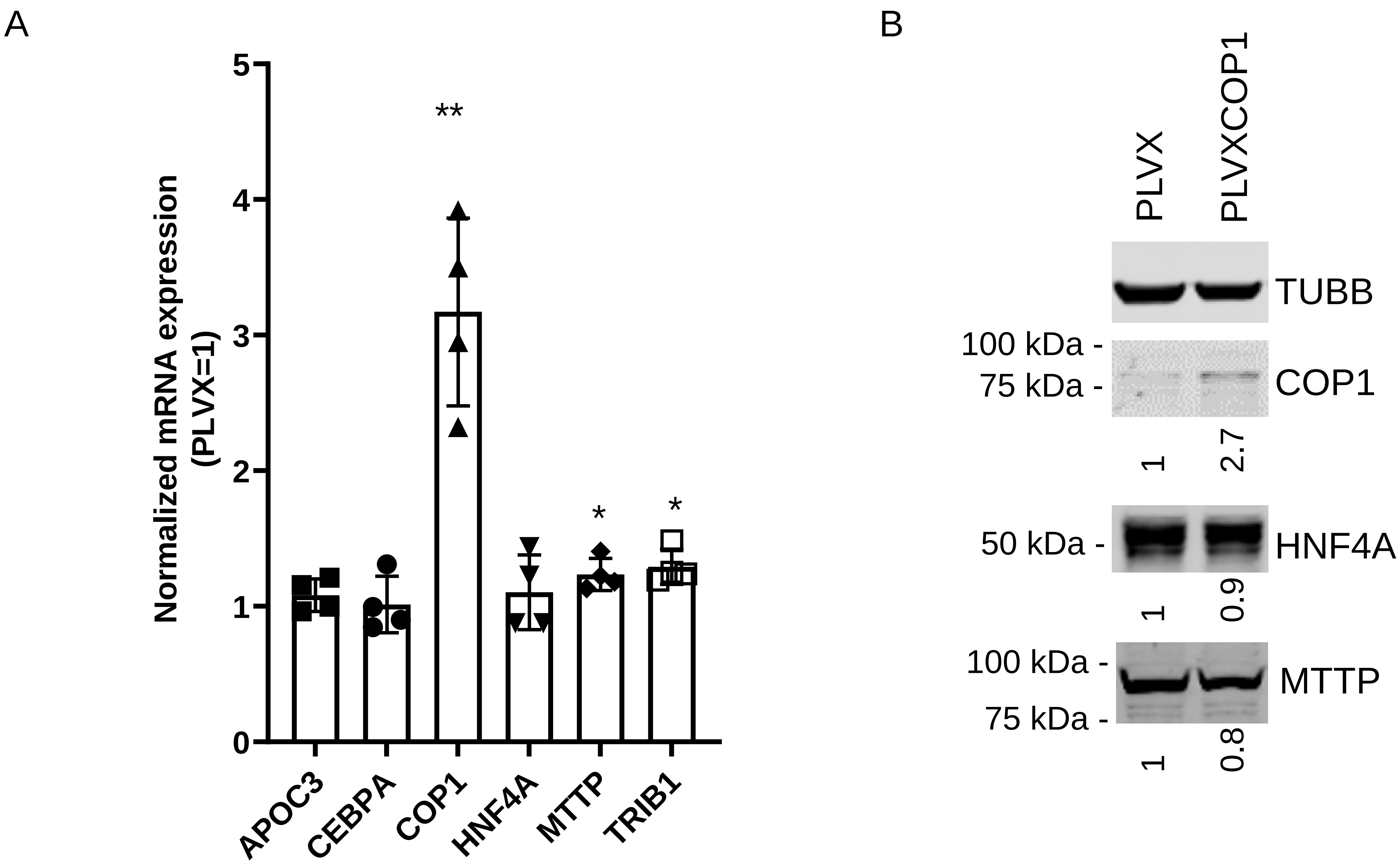


**Figure S7. Impact of COP1 overexpression on a panel of key liver transcripts**. A, A pool of HepG2 cells, stably transduced with COP1 were examined over 4 passages for the expression of a selection of transcripts by qRT-PCR. Results are expressed relative to the corresponding PLVX value. Results from four biological replicates (average and SD) are shown. Statistical significance was using Student's t-test. B, Western blot representative of 3 biological repeats. Values were normalized to TUBB (Tubulin beta chain) and are expressed relative to PLVX. Only COP1 protein abundance was significantly affected (3.2-fold, p=0.002, Student's t-test).

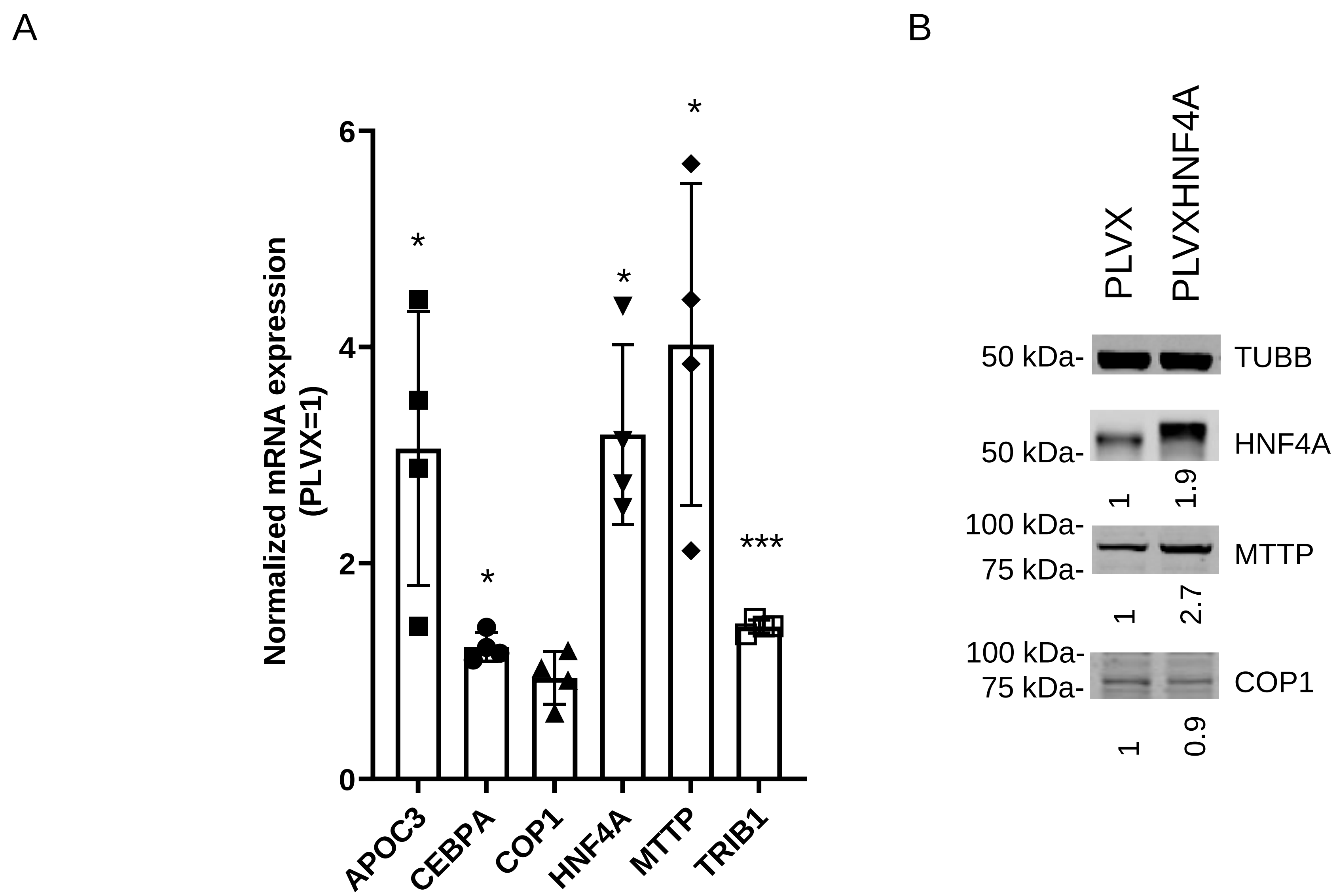


**Figure S8. Impact of HNF4A overexpression on a panel of key liver transcripts**. Stable pools of HepG2 cells, transduced with viral particles expressing either HNF4A or vector alone, were selected with puromycin and analyzed 10 to 14 days post infection for RNA and protein contents. A, qRT-PCR analyses of transcripts of interest. Values are normalized to the matching control (PLVX only) values. Student's (paired) *t*-test was performed comparing the HNF4 and PLVX values. Results from three biological replicates (average and SD) are shown. *, $p<0.05$;***,$<0.001$. B, Western blot analysis of protein lysates. Values were normalized to TUBB (Tubulin beta chain) and are expressed relative to PLVX. Representative of 3 experimental repeats.

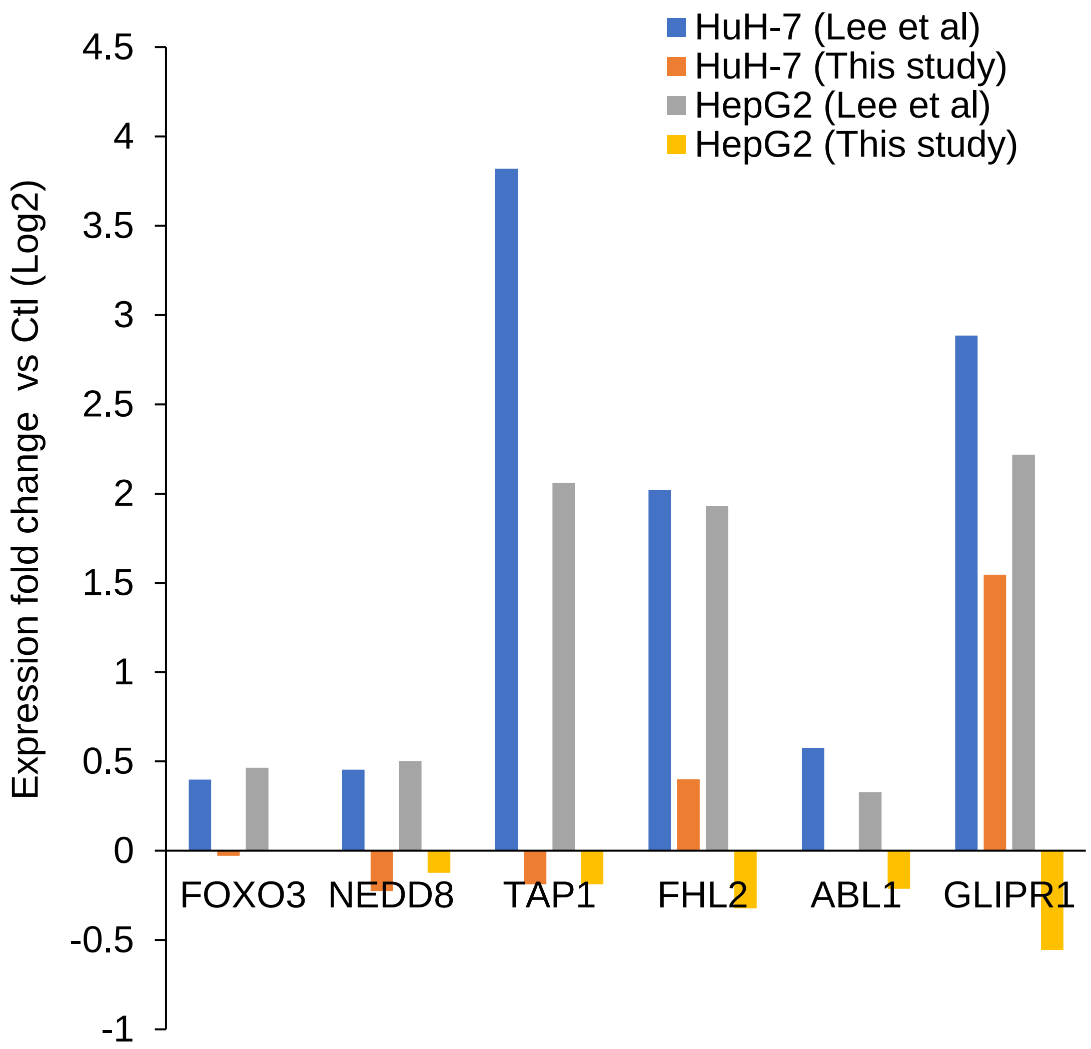


**Figure S9. Impact of COP1 silencing on a subset of genes linked to p53 regulation**. Transcript abundance changes as a result of COP1 silencing. Gene selection is by Lee et al. Values from both studies are shown.
