## Supplementary tables for "Constitutive Photomorphogenesis Protein 1 homolog (COP1) sustains nuclear factor-4 alpha function in human hepatocyte models"

**Enrichment analysis by Overerepresentation analysis (ORA) and Gene Set Enrichment analyses (GSEA). NCBI Entrez Genes were mapped via WebGestalt. GSEA was performed using the complete transcriptome fold-change dataset. Nominally changed transcripts were used for ORA. Default parameters were used. FDR significant hits (q < 0.05) hits are shown**

| | |
|---|---|
| **Table S1** | Over-representation of HepG2 and HuH-7 shared hits. |
| **Table S2** | ORA analyses of nominally changed transcripts in HepG2, HuH7 and Primary Hepatocytes |
| **Table S3** | ORA analyses using 3 randomized ID inputs |
| **Table S4** | Over-representation analysis of (nominally) upregulated or downregulated transcripts. |
| **Table S5** | Gene Set Enrichment analyses of transcriptome impacts of COP1 suppression in HuH-7, HepG2 and Hepatocytes |
| **Table S6** | Ingenuity Canonical Pathways identified in COP1 suppressed cells |
| **Table S7** | Upstream regulators predicted in at least one cell type by IPA upstream regulators |
| **Table S8** | Over-representation analyses of COP1 and HNF4A targeted hits |
| **Table S9** | Comparison of Gene list from Lee et al (Suppl Table 2) with values from current study. |

**Table S1. ORA React, KEGG and GO of HepG2 and HuH-7 shared hits.** Nominally significant hits common to HepG2 and HuH-7 cells were analyzed by ORA. Analysis performed via WebGestalt. Only FDR significant hits are shown. Size, Gene Set Size; Expect: Expected value by chance (number of hits mapped to both input and geneset); Ratio: Enrichment ratio (number of actual hits mapped to input and geneset overlap vs number of hits predicted by chance). P value, nominal pvalue of overlap. FDR, False Discovery Rate.

| GO (Biological_Process_noRedundant) | | | | | | |
|---|---|---|---|---|---|---|
| Gene Set | Description | Size | Expect | Ratio | P Value | FDR |
| GO:0006260 | DNA replication | 259 | 4.5 | 6.4 | 1.1E-15 | 9.4E-13 |
| GO:0044843 | cell cycle G1/S phase transitio | 236 | 4.1 | 4.9 | 5.3E-09 | 2.3E-06 |
| GO:0044772 | mitotic cell cycle phase transit | 464 | 8.1 | 3.3 | 3.8E-08 | 1.1E-05 |
| GO:0071103 | DNA conformation change | 230 | 4.0 | 4.0 | 2.7E-06 | 5.7E-04 |
| GO:0006301 | postreplication repair | 47 | 0.8 | 8.6 | 1.5E-05 | 2.6E-03 |
| GO:0000075 | cell cycle checkpoint | 211 | 3.7 | 3.5 | 8.2E-05 | 1.2E-02 |
| GO:0010948 | negative regulation of cell cyc | 266 | 4.6 | 3.0 | 2.3E-04 | 2.8E-02 |
| GO:0003170 | heart valve development | 52 | 0.9 | 6.6 | 2.7E-04 | 2.9E-02 |
| GO:0071897 | DNA biosynthetic process | 184 | 3.2 | 3.4 | 3.8E-04 | 3.0E-02 |
| GO:0002065 | columnar/cuboidal epithelial ( | 102 | 1.8 | 4.5 | 4.0E-04 | 3.0E-02 |
| GO:0042769 | DNA damage response, detec | 37 | 0.6 | 7.8 | 4.2E-04 | 3.0E-02 |
| GO:0031589 | cell-substrate adhesion | 316 | 5.5 | 2.7 | 4.3E-04 | 3.0E-02 |
| GO:0022616 | DNA strand elongation | 23 | 0.4 | 10.0 | 6.1E-04 | 4.0E-02 |
| GO:0048285 | organelle fission | 444 | 7.7 | 2.3 | 7.7E-04 | 4.4E-02 |
| GO:1901342 | regulation of vasculature deve | 300 | 5.2 | 2.7 | 7.8E-04 | 4.4E-02 |
| KEGG | | | | | | |
| Gene Set | Description | Size | Expect | Ratio | P Value | FDR |
| hsa04110 | Cell cycle | 119 | 2.2 | 6.5 | 2.3E-08 | 7.6E-06 |
| hsa03030 | DNA replication | 34 | 0.6 | 13.0 | 1.1E-07 | 1.8E-05 |
| REACTOME | | | | | | |
| Gene Set | Description | Size | Expect | Ratio | P Value | FDR |
| R-HSA-176974 | Unwinding of DNA | 12 | 0.2 | 41.3 | 3.7E-14 | 6.4E-11 |
| R-HSA-69190 | DNA strand elongation | 31 | 0.6 | 19.5 | 3.3E-12 | 2.8E-09 |
| R-HSA-176187 | Activation of ATR in response | 37 | 0.7 | 16.4 | 3.0E-11 | 1.7E-08 |
| R-HSA-69306 | DNA Replication | 124 | 2.3 | 7.5 | 7.8E-11 | 3.3E-08 |
| R-HSA-69239 | Synthesis of DNA | 116 | 2.1 | 7.6 | 2.6E-10 | 8.9E-08 |
| R-HSA-69242 | S Phase | 155 | 2.8 | 6.4 | 3.5E-10 | 9.9E-08 |
| R-HSA-1640170 | Cell Cycle | 609 | 11.1 | 3.2 | 8.3E-10 | 1.8E-07 |
| R-HSA-453279 | Mitotic G1-G1/S phases | 143 | 2.6 | 6.5 | 7.6E-10 | 1.8E-07 |
| R-HSA-69206 | G1/S Transition | 128 | 2.3 | 6.9 | 1.2E-09 | 2.2E-07 |
| R-HSA-68962 | Activation of the pre-replicati | 33 | 0.6 | 15.0 | 4.7E-09 | 8.0E-07 |
| R-HSA-69620 | Cell Cycle Checkpoints | 280 | 5.1 | 4.3 | 6.6E-09 | 1.0E-06 |
| R-HSA-69278 | Cell Cycle, Mitotic | 507 | 9.2 | 3.1 | 3.2E-08 | 4.6E-06 |
| R-HSA-539107 | Activation of E2F1 target gene | 28 | 0.5 | 13.8 | 5.0E-07 | 6.1E-05 |
| R-HSA-69205 | G1/S-Specific Transcription | 28 | 0.5 | 13.8 | 5.0E-07 | 6.1E-05 |
| R-HSA-69002 | DNA Replication Pre-Initiation | 84 | 1.5 | 6.5 | 2.7E-06 | 3.0E-04 |
| R-HSA-68867 | Assembly of the pre-replicativ | 67 | 1.2 | 7.4 | 3.1E-06 | 3.3E-04 |

| | | | | | | |
|---|---|---|---|---|---|---|
| R-HSA-69052 | Switching of origins to a post- | 88 | 1.6 | 6.3 | 4.1E-06 | 4.1E-04 |
| R-HSA-68949 | Orc1 removal from chromatin | 70 | 1.3 | 7.1 | 4.5E-06 | 4.2E-04 |
| R-HSA-1538133 | G0 and Early G1 | 27 | 0.5 | 12.2 | 7.2E-06 | 6.1E-04 |
| R-HSA-69481 | G2/M Checkpoints | 161 | 2.9 | 4.4 | 7.1E-06 | 6.1E-04 |
| R-HSA-73893 | DNA Damage Bypass | 48 | 0.9 | 8.0 | 2.3E-05 | 1.9E-03 |
| R-HSA-68689 | CDC6 association with the OR | 11 | 0.2 | 20.0 | 3.2E-05 | 2.4E-03 |
| R-HSA-73894 | DNA Repair | 302 | 5.5 | 2.9 | 1.2E-04 | 9.0E-03 |
| R-HSA-5685942 | HDR through Homologous Re | 64 | 1.2 | 6.0 | 1.5E-04 | 1.1E-02 |
| R-HSA-110314 | Recognition of DNA damage b | 30 | 0.5 | 9.2 | 1.9E-04 | 1.3E-02 |
| R-HSA-5656169 | Termination of translesion DN | 31 | 0.6 | 8.9 | 2.2E-04 | 1.4E-02 |
| R-HSA-1362277 | Transcription of E2F targets u | 19 | 0.3 | 11.6 | 3.3E-04 | 2.1E-02 |
| R-HSA-5693567 | HDR through Homologous Re | 123 | 2.2 | 4.0 | 4.0E-04 | 2.4E-02 |
| R-HSA-110313 | Translesion synthesis by Y fam | 38 | 0.7 | 7.2 | 5.8E-04 | 3.2E-02 |
| R-HSA-5693538 | Homology Directed Repair | 129 | 2.3 | 3.8 | 5.6E-04 | 3.2E-02 |
| R-HSA-5693616 | Presynaptic phase of homolog | 38 | 0.7 | 7.2 | 5.8E-04 | 3.2E-02 |
| R-HSA-5693579 | Homologous DNA Pairing and | 41 | 0.7 | 6.7 | 8.3E-04 | 4.4E-02 |

**Table S2. ORA analyses of nominally changed transcripts in HepG2, HuH7 and Primary Hepatocytes**. Over-representation analysis using Gene Ontology ("Biological_Process_noRedundant"), as well as KEGG and Reactome pathways datasets. Listed by increasing FDR values. Hits common to more than one cell type are highlighted in red.

GO

HuH-7

| Gene Set | Description | FDR |
|---|---|---|
| GO:0006260 | DNA replication | 3.7E-10 |
| GO:0044772 | mitotic cell cycle phase transition | 4.9E-06 |
| GO:0000075 | cell cycle checkpoint | 1.1E-05 |
| GO:0044843 | cell cycle G1/S phase transition | 2.7E-05 |
| GO:0071103 | DNA conformation change | 2.1E-04 |
| GO:0022616 | DNA strand elongation | 5.1E-04 |
| GO:1904888 | cranial skeletal system development | 1.2E-03 |
| GO:0006302 | double-strand break repair | 1.2E-02 |
| GO:0001525 | angiogenesis | 1.2E-02 |

HepG2

| Gene Set | Description | FDR |
|---|---|---|
| GO:0044282 | small molecule catabolic process | 4.5E-10 |
| GO:0006631 | fatty acid metabolic process | 2.2E-06 |
| GO:0006260 | DNA replication | 1.7E-05 |
| GO:0006732 | coenzyme metabolic process | 2.2E-04 |
| GO:1901615 | organic hydroxy compound metabolic p | 4.4E-04 |
| GO:0002576 | platelet degranulation | 4.4E-04 |
| GO:0002526 | acute inflammatory response | 6.5E-04 |
| GO:0072376 | protein activation cascade | 7.6E-04 |
| GO:0051604 | protein maturation | 8.1E-04 |
| GO:0019216 | regulation of lipid metabolic process | 1.1E-03 |
| GO:0006790 | sulfur compound metabolic process | 1.1E-03 |
| GO:0016053 | organic acid biosynthetic process | 1.2E-03 |
| GO:0055088 | lipid homeostasis | 2.5E-03 |
| GO:0008202 | steroid metabolic process | 2.7E-03 |
| GO:0006520 | cellular amino acid metabolic process | 3.6E-03 |
| GO:2000147 | positive regulation of cell motility | 4.2E-03 |
| GO:0051701 | interaction with host | 6.8E-03 |
| GO:0006979 | response to oxidative stress | 8.2E-03 |
| GO:0031589 | cell-substrate adhesion | 1.1E-02 |
| GO:0001525 | angiogenesis | 1.3E-02 |
| GO:0050817 | coagulation | 1.4E-02 |
| GO:0048732 | gland development | 1.4E-02 |
| GO:0019058 | viral life cycle | 1.4E-02 |

| | | |
|---|---|---|
| GO:0014812 | muscle cell migration | 1.5E-02 |
| GO:0061008 | hepaticobiliary system development | 1.5E-02 |
| GO:0009636 | response to toxic substance | 1.6E-02 |
| GO:0006638 | neutral lipid metabolic process | 1.7E-02 |
| GO:0051188 | cofactor biosynthetic process | 1.8E-02 |
| GO:0010038 | response to metal ion | 1.8E-02 |
| GO:0046434 | organophosphate catabolic process | 1.8E-02 |
| GO:0042558 | pteridine-containing compound metabo | 1.8E-02 |
| GO:0048545 | response to steroid hormone | 1.9E-02 |
| GO:0033865 | nucleoside bisphosphate metabolic pro | 2.0E-02 |
| GO:0042737 | drug catabolic process | 2.0E-02 |
| GO:0001101 | response to acid chemical | 2.0E-02 |
| GO:0016042 | lipid catabolic process | 2.0E-02 |
| GO:0050727 | regulation of inflammatory response | 2.1E-02 |
| GO:0006090 | pyruvate metabolic process | 2.1E-02 |
| GO:0034404 | nucleobase-containing small molecule | 2.1E-02 |
| GO:0044772 | mitotic cell cycle phase transition | 2.1E-02 |
| GO:0015849 | organic acid transport | 2.8E-02 |
| GO:1901342 | regulation of vasculature development | 3.8E-02 |
| GO:0032103 | positive regulation of response to exter | 3.8E-02 |
| GO:1903034 | regulation of response to wounding | 4.5E-02 |

### Hepatocytes

| Gene Set | Description | FDR |
|---|---|---|
| GO:0006631 | fatty acid metabolic process | 1.1E-06 |
| GO:0044282 | small molecule catabolic process | 7.8E-06 |
| GO:0015711 | organic anion transport | 8.0E-06 |
| GO:1901615 | organic hydroxy compound metabolic p | 8.0E-06 |
| GO:0010876 | lipid localization | 1.5E-05 |
| GO:0015849 | organic acid transport | 1.5E-05 |
| GO:0008202 | steroid metabolic process | 4.6E-05 |
| GO:0055088 | lipid homeostasis | 9.3E-05 |
| GO:1901568 | fatty acid derivative metabolic process | 1.2E-04 |
| GO:0050878 | regulation of body fluid levels | 2.0E-04 |
| GO:0050673 | epithelial cell proliferation | 3.9E-04 |
| GO:0007586 | digestion | 3.9E-04 |
| GO:0015850 | organic hydroxy compound transport | 4.1E-04 |
| GO:0016053 | organic acid biosynthetic process | 4.1E-04 |
| GO:0048732 | gland development | 4.8E-04 |
| GO:0006790 | sulfur compound metabolic process | 1.8E-03 |
| GO:0097164 | ammonium ion metabolic process | 2.0E-03 |
| GO:0062012 | regulation of small molecule metabolic | 4.2E-03 |
| GO:0010038 | response to metal ion | 4.7E-03 |
| GO:0002449 | lymphocyte mediated immunity | 4.7E-03 |
| GO:0006959 | humoral immune response | 4.7E-03 |

| | | |
|---|---|---|
| GO:0019216 | regulation of lipid metabolic process | 4.7E-03 |
| GO:0009410 | response to xenobiotic stimulus | 4.7E-03 |
| GO:0042445 | hormone metabolic process | 5.1E-03 |
| GO:0043491 | protein kinase B signaling | 5.1E-03 |
| GO:0042737 | drug catabolic process | 5.1E-03 |
| GO:0009112 | nucleobase metabolic process | 5.1E-03 |
| GO:0006575 | cellular modified amino acid metabolic | 6.4E-03 |
| GO:0006732 | coenzyme metabolic process | 7.6E-03 |
| GO:0043062 | extracellular structure organization | 9.4E-03 |
| GO:0032528 | microvillus organization | 1.1E-02 |
| GO:0070555 | response to interleukin-1 | 1.2E-02 |
| GO:0001655 | urogenital system development | 1.4E-02 |
| GO:0007015 | actin filament organization | 1.4E-02 |
| GO:0015748 | organophosphate ester transport | 1.4E-02 |
| GO:0014812 | muscle cell migration | 1.4E-02 |
| GO:0071241 | cellular response to inorganic substanc | 1.5E-02 |
| GO:0016042 | lipid catabolic process | 1.5E-02 |
| GO:0002576 | platelet degranulation | 1.5E-02 |
| GO:0009636 | response to toxic substance | 1.5E-02 |
| GO:0050817 | coagulation | 1.9E-02 |
| GO:0072376 | protein activation cascade | 2.1E-02 |
| GO:0070371 | ERK1 and ERK2 cascade | 2.1E-02 |
| GO:0003013 | circulatory system process | 2.9E-02 |
| GO:0031667 | response to nutrient levels | 3.8E-02 |
| GO:0072523 | purine-containing compound catabolic | 3.9E-02 |
| GO:0051188 | cofactor biosynthetic process | 4.5E-02 |
| GO:0090130 | tissue migration | 4.8E-02 |
| GO:0046486 | glycerolipid metabolic process | 4.9E-02 |

KEGG

HuH-7

| Gene Set | Description | FDR |
|---|---|---|
| hsa04110 | Cell cycle | 1.0E-07 |
| hsa03030 | DNA replication | 3.9E-06 |
| hsa03430 | Mismatch repair | 2.7E-02 |

HepG2

| Gene Set | Description | FDR |
|---|---|---|
| hsa01100 | Metabolic pathways | 5.7E-07 |
| hsa04610 | Complement and coagulation cascade | 5.0E-06 |
| hsa00100 | Steroid biosynthesis | 7.3E-04 |
| hsa03030 | DNA replication | 2.0E-02 |
| hsa00480 | Glutathione metabolism | 3.6E-02 |
| hsa04975 | Fat digestion and absorption | 3.9E-02 |

### Hepatocytes

| Gene Set | Description | FDR |
|---|---|---|
| hsa00982 | Drug metabolism | 8.6E-07 |
| hsa05204 | Chemical carcinogenesis | 2.8E-06 |
| hsa00830 | Retinol metabolism | 4.3E-06 |
| hsa00980 | Metabolism of xenobiotics by cytochror | 4.3E-06 |
| hsa00140 | Steroid hormone biosynthesis | 4.0E-05 |
| hsa00983 | Drug metabolism | 4.0E-05 |
| hsa00071 | Fatty acid degradation | 2.0E-03 |
| hsa00340 | Histidine metabolism | 2.0E-03 |
| hsa04978 | Mineral absorption | 3.0E-03 |
| hsa00053 | Ascorbate and aldarate metabolism | 4.5E-03 |
| hsa04610 | Complement and coagulation cascade | 7.1E-03 |
| hsa03320 | PPAR signaling pathway | 1.1E-02 |
| hsa00040 | Pentose and glucuronate interconversi | 1.1E-02 |
| hsa00380 | Tryptophan metabolism | 1.3E-02 |
| hsa01212 | Fatty acid metabolism | 2.5E-02 |
| hsa00410 | beta-Alanine metabolism | 2.5E-02 |
| hsa00480 | Glutathione metabolism | 2.5E-02 |
| hsa01100 | Metabolic pathways | 2.5E-02 |

REACTOME

### HuH-7

| Gene Set | Description | FDR |
|---|---|---|
| R-HSA-68962 | Activation of the pre-replicative comple | 4.68E-11 |
| R-HSA-176187 | Activation of ATR in response to replica | 4.68E-11 |
| R-HSA-69306 | DNA Replication | 1.21E-09 |
| R-HSA-69190 | DNA strand elongation | 1.31E-08 |
| R-HSA-1640170 | Cell Cycle | 1.78E-08 |
| R-HSA-69278 | Cell Cycle, Mitotic | 1.84E-07 |
| R-HSA-176974 | Unwinding of DNA | 1.84E-07 |
| R-HSA-453279 | Mitotic G1-G1/S phases | 1.84E-07 |
| R-HSA-69620 | Cell Cycle Checkpoints | 1.44E-06 |
| R-HSA-69206 | G1/S Transition | 1.54E-06 |
| R-HSA-69239 | Synthesis of DNA | 1.75E-06 |
| R-HSA-69242 | S Phase | 4.43E-06 |
| R-HSA-69002 | DNA Replication Pre-Initiation | 2.48E-05 |
| R-HSA-69481 | G2/M Checkpoints | 2.35E-03 |
| R-HSA-1538133 | G0 and Early G1 | 3.58E-03 |
| R-HSA-73886 | Chromosome Maintenance | 3.58E-03 |
| R-HSA-539107 | Activation of E2F1 target genes at G1/S | 4.25E-03 |
| R-HSA-69205 | G1/S-Specific Transcription | 4.25E-03 |
| R-HSA-5693538 | Homology Directed Repair | 4.25E-03 |
| R-HSA-174417 | Telomere C-strand (Lagging Strand) Sy | 4.25E-03 |

| | | |
|---|---|---|
| R-HSA-180786 | Extension of Telomeres | 5.49E-03 |
| R-HSA-68689 | CDC6 association with the ORC:origin | 7.94E-03 |
| R-HSA-5651801 | PCNA-Dependent Long Patch Base Ex | 8.17E-03 |
| R-HSA-73894 | DNA Repair | 2.15E-02 |
| R-HSA-72649 | Translation initiation complex formatior | 2.45E-02 |
| R-HSA-5693532 | DNA Double-Strand Break Repair | 2.50E-02 |
| R-HSA-156827 | L13a-mediated translational silencing c | 2.50E-02 |
| R-HSA-72662 | Activation of the mRNA upon binding o | 2.50E-02 |
| R-HSA-72702 | Ribosomal scanning and start codon re | 2.50E-02 |
| R-HSA-5693567 | HDR through Homologous Recombina | 2.50E-02 |
| R-HSA-5696397 | Gap-filling DNA repair synthesis and lig | 2.62E-02 |
| R-HSA-110314 | Recognition of DNA damage by PCNA | 2.74E-02 |
| R-HSA-69186 | Lagging Strand Synthesis | 2.79E-02 |
| R-HSA-72706 | GTP hydrolysis and joining of the 60S | 2.79E-02 |
| R-HSA-5685942 | HDR through Homologous Recombina | 3.01E-02 |
| R-HSA-110373 | Resolution of AP sites via the multiple- | 3.08E-02 |
| R-HSA-69052 | Switching of origins to a post-replicative | 3.43E-02 |
| R-HSA-73884 | Base Excision Repair | 3.43E-02 |
| R-HSA-73933 | Resolution of Abasic Sites (AP sites) | 3.43E-02 |
| R-HSA-5693616 | Presynaptic phase of homologous DNA | 3.43E-02 |

HepG2

| Gene Set | Description | FDR |
|---|---|---|
| R-HSA-1430728 | Metabolism | 9.7E-07 |
| R-HSA-176974 | Unwinding of DNA | 8.4E-05 |
| R-HSA-69190 | DNA strand elongation | 1.1E-04 |
| R-HSA-114608 | Platelet degranulation | 7.4E-04 |
| R-HSA-1640170 | Cell Cycle | 1.3E-03 |
| R-HSA-539107 | Activation of E2F1 target genes at G1/S | 1.3E-03 |
| R-HSA-69205 | G1/S-Specific Transcription | 1.3E-03 |
| R-HSA-76002 | Platelet activation, signaling and aggre | 1.3E-03 |
| R-HSA-76005 | Response to elevated platelet cytosolic | 1.3E-03 |
| R-HSA-69278 | Cell Cycle, Mitotic | 1.4E-03 |
| R-HSA-109582 | Hemostasis | 1.5E-03 |
| R-HSA-191273 | Cholesterol biosynthesis | 2.5E-03 |
| R-HSA-977606 | Regulation of Complement cascade | 2.5E-03 |
| R-HSA-166658 | Complement cascade | 3.3E-03 |
| R-HSA-6788656 | Histidine, lysine, phenylalanine, tyrosin | 5.1E-03 |
| R-HSA-174577 | Activation of C3 and C5 | 5.8E-03 |
| R-HSA-211859 | Biological oxidations | 6.2E-03 |
| R-HSA-556833 | Metabolism of lipids | 1.1E-02 |
| R-HSA-8957275 | Post-translational protein phosphorylat | 1.2E-02 |
| R-HSA-8978868 | Fatty acid metabolism | 1.5E-02 |
| R-HSA-69620 | Cell Cycle Checkpoints | 1.5E-02 |
| R-HSA-75205 | Dissolution of Fibrin Clot | 2.8E-02 |
| R-HSA-1538133 | G0 and Early G1 | 3.5E-02 |

| | | |
|---|---|---|
| R-HSA-8957322 | Metabolism of steroids | 3.5E-02 |
| R-HSA-381426 | Regulation of Insulin-like Growth Facto | 4.4E-02 |
| R-HSA-174411 | Polymerase switching on the C-strand | 4.8E-02 |
| R-HSA-69091 | Polymerase switching | 4.8E-02 |
| R-HSA-69109 | Leading Strand Synthesis | 4.8E-02 |

### Hepatocytes

| Gene Set | Description | FDR |
|---|---|---|
| R-HSA-211859 | Biological oxidations | 6.3E-10 |
| R-HSA-1430728 | Metabolism | 3.2E-06 |
| R-HSA-156580 | Phase II - Conjugation of compounds | 7.6E-06 |
| R-HSA-211945 | Phase I - Functionalization of compour | 4.3E-04 |
| R-HSA-556833 | Metabolism of lipids | 9.5E-04 |
| R-HSA-3000171 | Non-integrin membrane-ECM interactic | 2.8E-03 |
| R-HSA-8963899 | Plasma lipoprotein remodeling | 2.8E-03 |
| R-HSA-211958 | Miscellaneous substrates | 4.2E-03 |
| R-HSA-211897 | Cytochrome P450 - arranged by substr | 6.4E-03 |
| R-HSA-211981 | Xenobiotics | 1.2E-02 |
| R-HSA-156588 | Glucuronidation | 1.2E-02 |
| R-HSA-5660526 | Response to metal ions | 1.2E-02 |
| R-HSA-8978868 | Fatty acid metabolism | 1.2E-02 |
| R-HSA-3000170 | Syndecan interactions | 1.2E-02 |
| R-HSA-5661231 | Metallothioneins bind metals | 1.2E-02 |
| R-HSA-211935 | Fatty acids | 1.8E-02 |
| R-HSA-2142753 | Arachidonic acid metabolism | 1.8E-02 |
| R-HSA-211979 | Eicosanoids | 2.1E-02 |
| R-HSA-174824 | Plasma lipoprotein assembly, remodeli | 2.1E-02 |
| R-HSA-8963889 | Assembly of active LPL and LIPC lipas | 2.7E-02 |

**Table S3. ORA analyses using 3 randomized ID inputs from .** Over-representation analysis using KEGG and Reactome terms was performed on 3 lists consisting 4262 randomized IDs from the HuGene 2.0 array. Listed by increasing FDR values. Hits shared by more than one list are highlighted in red (KEGG) or yellow (Reactome).

KEGG

Set 1

| Gene Set | Description | FDR |
|---|---|---|
| hsa04612 | Antigen processing and presentation | 4.6E-05 |
| hsa05332 | Graft-versus-host disease | 1.2E-03 |
| hsa05330 | Allograft rejection | 1.4E-03 |
| hsa04940 | Type I diabetes mellitus | 4.0E-03 |
| hsa05310 | Asthma | 1.0E-02 |
| hsa05145 | Toxoplasmosis | 1.0E-02 |
| hsa05320 | Autoimmune thyroid disease | 1.0E-02 |
| hsa04145 | Phagosome | 1.6E-02 |
| hsa05416 | Viral myocarditis | 1.8E-02 |
| hsa00500 | Starch and sucrose metabolism | 1.8E-02 |
| hsa05322 | Systemic lupus erythematosus | 7.5E-02 |
| hsa05150 | Staphylococcus aureus infection | 7.7E-02 |
| hsa05166 | Human T-cell leukemia virus 1 infection | 1.5E-01 |
| hsa05323 | Rheumatoid arthritis | 1.9E-01 |
| hsa04672 | Intestinal immune network for IgA production | 1.9E-01 |
| hsa04213 | Longevity regulating pathway | 2.5E-01 |
| hsa05168 | Herpes simplex infection | 3.1E-01 |
| hsa04914 | Progesterone-mediated oocyte maturation | 3.1E-01 |
| hsa04640 | Hematopoietic cell lineage | 3.2E-01 |
| hsa04218 | Cellular senescence | 3.3E-01 |

Set 2

| Gene Set | Description | FDR |
|---|---|---|
| hsa04612 | Antigen processing and presentation | 4.0E-04 |
| hsa05332 | Graft-versus-host disease | 4.0E-04 |
| hsa05330 | Allograft rejection | 2.9E-03 |
| hsa05416 | Viral myocarditis | 4.3E-03 |
| hsa04940 | Type I diabetes mellitus | 5.7E-03 |
| hsa05320 | Autoimmune thyroid disease | 1.7E-02 |
| hsa05310 | Asthma | 1.8E-02 |
| hsa04672 | Intestinal immune network for IgA production | 5.7E-02 |
| hsa04650 | Natural killer cell mediated cytotoxicity | 8.9E-02 |
| hsa05168 | Herpes simplex infection | 9.4E-02 |
| hsa05164 | Influenza A | 1.0E-01 |
| hsa04640 | Hematopoietic cell lineage | 1.0E-01 |
| hsa05166 | Human T-cell leukemia virus 1 infection | 2.0E-01 |

| | | |
|---|---|---|
| hsa01521 | EGFR tyrosine kinase inhibitor resistance | 2.1E-01 |
| hsa03015 | mRNA surveillance pathway | 2.1E-01 |
| hsa04145 | Phagosome | 2.3E-01 |
| hsa04064 | NF-kappa B signaling pathway | 2.6E-01 |
| hsa04977 | Vitamin digestion and absorption | 3.2E-01 |
| hsa05145 | Toxoplasmosis | 3.9E-01 |
| hsa05219 | Bladder cancer | 3.9E-01 |

Set 3

| Gene Set | Description | FDR |
|---|---|---|
| hsa04940 | Type I diabetes mellitus | 1.4E-02 |
| hsa03008 | Ribosome biogenesis in eukaryotes | 6.7E-02 |
| hsa05310 | Asthma | 7.1E-02 |
| hsa05322 | Systemic lupus erythematosus | 7.1E-02 |
| hsa05330 | Allograft rejection | 7.1E-02 |
| hsa05169 | Epstein-Barr virus infection | 2.9E-01 |
| hsa04714 | Thermogenesis | 2.9E-01 |
| hsa05332 | Graft-versus-host disease | 2.9E-01 |
| hsa05140 | Leishmaniasis | 3.6E-01 |
| hsa05012 | Parkinson disease | 3.6E-01 |
| hsa04926 | Relaxin signaling pathway | 5.2E-01 |
| hsa00190 | Oxidative phosphorylation | 5.2E-01 |
| hsa05145 | Toxoplasmosis | 5.6E-01 |
| hsa05321 | Inflammatory bowel disease (IBD) | 5.6E-01 |
| hsa04659 | Th17 cell differentiation | 5.6E-01 |
| hsa01523 | Antifolate resistance | 5.6E-01 |
| hsa04260 | Cardiac muscle contraction | 5.6E-01 |
| hsa04612 | Antigen processing and presentation | 5.9E-01 |
| hsa05320 | Autoimmune thyroid disease | 6.8E-01 |
| hsa05168 | Herpes simplex infection | 6.8E-01 |

REACTOME

set1

| Gene Set | Description | FDR |
|---|---|---|
| R-HSA-189085 | Digestion of dietary carbohydrate | 4.7E-01 |
| R-HSA-3371568 | Attenuation phase | 7.5E-01 |
| R-HSA-202430 | Translocation of ZAP-70 to Immunological synapse | 7.5E-01 |
| R-HSA-8935690 | Digestion | 7.5E-01 |
| R-HSA-3560782 | Diseases associated with glycosaminoglycan metabolism | 7.5E-01 |
| R-HSA-3371571 | HSF1-dependent transactivation | 7.5E-01 |
| R-HSA-202427 | Phosphorylation of CD3 and TCR zeta chains | 7.5E-01 |
| R-HSA-174577 | Activation of C3 and C5 | 7.5E-01 |
| R-HSA-3656225 | Defective CHST6 causes MCDC1 | 7.5E-01 |
| R-HSA-3656243 | Defective ST3GAL3 causes MCT12 and EIEE15 | 7.5E-01 |
| R-HSA-3656244 | Defective B4GALT1 causes B4GALT1-CDG (CDG-2d) | 7.5E-01 |
| R-HSA-389948 | PD-1 signaling | 7.7E-01 |

| | | |
|---|---|---|
| R-HSA-2129379 | Molecules associated with elastic fibres | 1.0E+00 |
| R-HSA-8963743 | Digestion and absorption | 1.0E+00 |
| R-HSA-2022854 | Keratan sulfate biosynthesis | 1.0E+00 |
| R-HSA-877300 | Interferon gamma signaling | 1.0E+00 |
| R-HSA-5603029 | IkBA variant leads to EDA-ID | 1.0E+00 |
| R-HSA-3371453 | Regulation of HSF1-mediated heat shock response | 1.0E+00 |
| R-HSA-446219 | Synthesis of substrates in N-glycan biosythesis | 1.0E+00 |
| R-HSA-446193 | Biosynthesis of the N-glycan precursor (dolichol lipid-linked ol | 1.0E+00 |

set2

| Gene Set | Description | FDR |
|---|---|---|
| R-HSA-1181150 | Signaling by NODAL | 7.8E-01 |
| R-HSA-1236975 | Antigen processing-Cross presentation | 1.0E+00 |
| R-HSA-1236977 | Endosomal/Vacuolar pathway | 1.0E+00 |
| R-HSA-1433617 | Regulation of signaling by NODAL | 1.0E+00 |
| R-HSA-1643685 | Disease | 1.0E+00 |
| R-HSA-202427 | Phosphorylation of CD3 and TCR zeta chains | 7.8E-01 |
| R-HSA-202430 | Translocation of ZAP-70 to Immunological synapse | 1.0E+00 |
| R-HSA-2187335 | The retinoid cycle in cones (daylight vision) | 5.8E-01 |
| R-HSA-2428928 | IRS-related events triggered by IGF1R | 1.0E+00 |
| R-HSA-352230 | Amino acid transport across the plasma membrane | 1.0E+00 |
| R-HSA-376176 | Signaling by ROBO receptors | 1.0E+00 |
| R-HSA-389948 | PD-1 signaling | 1.0E+00 |
| R-HSA-444257 | RSK activation | 1.0E+00 |
| R-HSA-5579029 | Metabolic disorders of biological oxidation enzymes | 7.8E-01 |
| R-HSA-5676594 | TNF receptor superfamily (TNFSF) members mediating non-ca | 7.8E-01 |
| R-HSA-877300 | Interferon gamma signaling | 7.8E-01 |
| R-HSA-8866910 | TFAP2 (AP-2) family regulates transcription of growth factors | 9.9E-01 |
| R-HSA-9010553 | Regulation of expression of SLITs and ROBOs | 1.0E+00 |
| R-HSA-927802 | Nonsense-Mediated Decay (NMD) | 7.8E-01 |
| R-HSA-975957 | Nonsense Mediated Decay (NMD) enhanced by the Exon Junc | 7.8E-01 |

set3

| Gene Set | Description | FDR |
|---|---|---|
| R-HSA-5689880 | Ub-specific processing proteases | 3.7E-02 |
| R-HSA-389948 | PD-1 signaling | 3.9E-02 |
| R-HSA-202424 | Downstream TCR signaling | 3.9E-02 |
| R-HSA-202403 | TCR signaling | 9.5E-02 |
| R-HSA-5688426 | Deubiquitination | 2.7E-01 |
| R-HSA-202427 | Phosphorylation of CD3 and TCR zeta chains | 2.7E-01 |
| R-HSA-388841 | Costimulation by the CD28 family | 2.7E-01 |
| R-HSA-202430 | Translocation of ZAP-70 to Immunological synapse | 4.5E-01 |
| R-HSA-392499 | Metabolism of proteins | 9.9E-01 |
| R-HSA-1169091 | Activation of NF-kappaB in B cells | 9.9E-01 |
| R-HSA-2871837 | FCERI mediated NF-kB activation | 9.9E-01 |
| R-HSA-1428517 | The citric acid (TCA) cycle and respiratory electron transport | 9.9E-01 |
| R-HSA-163200 | Respiratory electron transport, ATP synthesis by chemiosmoti | 1.0E+00 |

| | | |
|---|---|---|
| R-HSA-597592 | Post-translational protein modification | 1.0E+00 |
| R-HSA-2262752 | Cellular responses to stress | 1.0E+00 |
| R-HSA-1280218 | Adaptive Immune System | 1.0E+00 |
| R-HSA-202433 | Generation of second messenger molecules | 1.0E+00 |
| R-HSA-5676590 | NIK-->noncanonical NF-kB signaling | 1.0E+00 |
| R-HSA-5684264 | MAP3K8 (TPL2)-dependent MAPK1/3 activation | 1.0E+00 |
| R-HSA-977443 | GABA receptor activation | 1.0E+00 |

**Table S4. Over-representation analysis of (nominally) upregulated or downregulated transcripts.** Nominally significant transcripts were analyzed by ORA using Gene Ontology "Biological_Process_noRedundant" categories. Green highlights indicate shared downregulated Gene Sets. Pink highlights point to the extensive overlap between HuH-7 downregulated and HepG2 upregulated ontologies. No shared upregulated categories were identified. No upregulated HuH-7 gene sets passed 0.05 FDR significance (Top 10 nominally significant hits are shown).

### HepG2

downregulated

| Gene Set | Description | FDR |
|---|---|---|
| GO:0044282 | small molecule catabolic process | <2.2e-16 |
| GO:0006631 | fatty acid metabolic process | 9.4E-14 |
| GO:0006732 | coenzyme metabolic process | 9.0E-12 |
| GO:1901615 | organic hydroxy compound metabolic process | 3.1E-11 |
| GO:0008202 | steroid metabolic process | 4.5E-10 |
| GO:0072376 | protein activation cascade | 3.0E-09 |
| GO:0016053 | organic acid biosynthetic process | 7.2E-09 |
| GO:0019216 | regulation of lipid metabolic process | 2.1E-08 |
| GO:0006790 | sulfur compound metabolic process | 6.7E-08 |
| GO:0055088 | lipid homeostasis | 1.3E-07 |
| GO:0006091 | generation of precursor metabolites and energy | 1.5E-07 |
| GO:0006520 | cellular amino acid metabolic process | 5.3E-07 |
| GO:0051188 | cofactor biosynthetic process | 5.3E-07 |
| GO:0009259 | ribonucleotide metabolic process | 5.3E-07 |
| GO:0002526 | acute inflammatory response | 5.6E-07 |
| GO:0016042 | lipid catabolic process | 5.6E-07 |
| GO:0033865 | nucleoside bisphosphate metabolic process | 1.1E-06 |
| GO:0030258 | lipid modification | 3.8E-06 |
| GO:0010876 | lipid localization | 3.9E-06 |
| GO:0015711 | organic anion transport | 4.2E-06 |
| GO:0051604 | protein maturation | 1.1E-05 |
| GO:0006638 | neutral lipid metabolic process | 1.5E-05 |
| GO:0016999 | antibiotic metabolic process | 1.6E-05 |
| GO:0006090 | pyruvate metabolic process | 1.8E-05 |
| GO:0005996 | monosaccharide metabolic process | 3.2E-05 |
| GO:0015849 | organic acid transport | 3.3E-05 |
| GO:0042180 | cellular ketone metabolic process | 3.3E-05 |
| GO:0046390 | ribose phosphate biosynthetic process | 6.3E-05 |
| GO:0006575 | cellular modified amino acid metabolic process | 8.3E-05 |

| | | |
|---|---|---|
| GO:0007031 | peroxisome organization | 8.4E-05 |
| GO:0016052 | carbohydrate catabolic process | 9.5E-05 |
| GO:0062012 | regulation of small molecule metabolic process | 2.0E-04 |
| GO:0002576 | platelet degranulation | 2.1E-04 |
| GO:0043648 | dicarboxylic acid metabolic process | 2.5E-04 |
| GO:1901293 | nucleoside phosphate biosynthetic process | 3.2E-04 |
| GO:0097006 | regulation of plasma lipoprotein particle levels | 3.3E-04 |
| GO:0043574 | peroxisomal transport | 3.4E-04 |
| GO:0072524 | pyridine-containing compound metabolic process | 3.9E-04 |
| GO:0042737 | drug catabolic process | 5.3E-04 |
| GO:0071825 | protein-lipid complex subunit organization | 5.6E-04 |
| GO:0031667 | response to nutrient levels | 5.6E-04 |
| GO:0046939 | nucleotide phosphorylation | 1.1E-03 |
| GO:0034976 | response to endoplasmic reticulum stress | 2.0E-03 |
| GO:0051181 | cofactor transport | 3.3E-02 |

### upregulated

| Gene Set | Description | FDR |
|---|---|---|
| GO:0006260 | DNA replication | <2.2e-16 |
| GO:0044772 | mitotic cell cycle phase transition | 6.7E-06 |
| GO:2000147 | positive regulation of cell motility | 1.7E-05 |
| GO:0007059 | chromosome segregation | 2.2E-05 |
| GO:0051052 | regulation of DNA metabolic process | 2.2E-05 |
| GO:0000075 | cell cycle checkpoint | 2.3E-04 |
| GO:0071103 | DNA conformation change | 3.0E-04 |
| GO:0001667 | ameboidal-type cell migration | 3.0E-04 |
| GO:0032200 | telomere organization | 5.8E-04 |
| GO:0031032 | actomyosin structure organization | 1.1E-03 |
| GO:0071897 | DNA biosynthetic process | 1.2E-03 |
| GO:0045930 | negative regulation of mitotic cell cycle | 1.2E-03 |
| GO:0090130 | tissue migration | 1.3E-03 |
| GO:0014812 | muscle cell migration | 1.4E-03 |
| GO:1901987 | regulation of cell cycle phase transition | 1.4E-03 |
| GO:0048285 | organelle fission | 1.4E-03 |
| GO:0044843 | cell cycle G1/S phase transition | 1.4E-03 |
| GO:0031023 | microtubule organizing center organization | 2.3E-03 |
| GO:0031589 | cell-substrate adhesion | 2.3E-03 |
| GO:0060249 | anatomical structure homeostasis | 3.5E-03 |
| GO:0044839 | cell cycle G2/M phase transition | 3.7E-03 |
| GO:0006302 | double-strand break repair | 5.3E-03 |
| GO:0006310 | DNA recombination | 5.6E-03 |
| GO:0051302 | regulation of cell division | 5.8E-03 |
| GO:0009314 | response to radiation | 5.8E-03 |
| GO:0032970 | regulation of actin filament-based process | 5.8E-03 |
| GO:0022406 | membrane docking | 6.2E-03 |
| GO:0048771 | tissue remodeling | 9.5E-03 |

| | | |
|---|---|---|
| GO:0007229 | integrin-mediated signaling pathway | 9.6E-03 |
| GO:0038127 | ERBB signaling pathway | 1.1E-02 |
| GO:0071900 | regulation of protein serine/threonine kinase activity | 1.1E-02 |
| GO:0010948 | negative regulation of cell cycle process | 1.1E-02 |
| GO:0034330 | cell junction organization | 1.1E-02 |
| GO:0034502 | protein localization to chromosome | 1.1E-02 |
| GO:0003170 | heart valve development | 1.3E-02 |
| GO:0051493 | regulation of cytoskeleton organization | 1.3E-02 |
| GO:0022604 | regulation of cell morphogenesis | 1.3E-02 |
| GO:0022616 | DNA strand elongation | 1.3E-02 |
| GO:0007163 | establishment or maintenance of cell polarity | 1.4E-02 |
| GO:0003158 | endothelium development | 1.5E-02 |
| GO:0050867 | positive regulation of cell activation | 1.7E-02 |
| GO:0002064 | epithelial cell development | 1.7E-02 |
| GO:0043588 | skin development | 1.7E-02 |
| GO:0006333 | chromatin assembly or disassembly | 1.8E-02 |
| GO:0104004 | cellular response to environmental stimulus | 2.0E-02 |
| GO:0048008 | platelet-derived growth factor receptor signaling pathway | 2.9E-02 |
| GO:1902532 | negative regulation of intracellular signal transduction | 3.4E-02 |
| GO:0003007 | heart morphogenesis | 3.4E-02 |

### Primary hepatocytes

downregulated

| Gene Set | Description | FDR |
|---|---|---|
| GO:0044282 | small molecule catabolic process | 5.2E-08 |
| GO:0006631 | fatty acid metabolic process | 1.7E-07 |
| GO:0006575 | cellular modified amino acid metabolic process | 4.0E-05 |
| GO:0015711 | organic anion transport | 5.9E-05 |
| GO:0015849 | organic acid transport | 7.5E-05 |
| GO:0010876 | lipid localization | 9.2E-05 |
| GO:0016053 | organic acid biosynthetic process | 9.8E-05 |
| GO:0009410 | response to xenobiotic stimulus | 9.8E-05 |
| GO:0008202 | steroid metabolic process | 3.4E-04 |
| GO:0042180 | cellular ketone metabolic process | 5.3E-04 |
| GO:0055088 | lipid homeostasis | 6.5E-04 |
| GO:1901615 | organic hydroxy compound metabolic process | 1.6E-03 |
| GO:0015850 | organic hydroxy compound transport | 2.0E-03 |
| GO:0032528 | microvillus organization | 2.6E-03 |
| GO:1901568 | fatty acid derivative metabolic process | 2.7E-03 |
| GO:0006520 | cellular amino acid metabolic process | 2.8E-03 |
| GO:0016042 | lipid catabolic process | 4.1E-03 |
| GO:0097164 | ammonium ion metabolic process | 4.9E-03 |
| GO:0007015 | actin filament organization | 7.8E-03 |
| GO:0030258 | lipid modification | 8.7E-03 |
| GO:0007586 | digestion | 1.0E-02 |
| GO:0006732 | coenzyme metabolic process | 1.0E-02 |

| | | |
|---|---|---|
| GO:0015748 | organophosphate ester transport | 1.1E-02 |
| GO:0055081 | anion homeostasis | 1.2E-02 |
| GO:0042737 | drug catabolic process | 1.2E-02 |
| GO:0030856 | regulation of epithelial cell differentiation | 1.3E-02 |
| GO:0070371 | ERK1 and ERK2 cascade | 1.3E-02 |
| GO:0051051 | negative regulation of transport | 1.3E-02 |
| GO:0050878 | regulation of body fluid levels | 1.4E-02 |
| GO:0042445 | hormone metabolic process | 1.6E-02 |
| GO:0042692 | muscle cell differentiation | 1.9E-02 |
| GO:0043270 | positive regulation of ion transport | 2.0E-02 |
| GO:0016999 | antibiotic metabolic process | 2.1E-02 |
| GO:0051271 | negative regulation of cellular component movement | 2.2E-02 |
| GO:0010959 | regulation of metal ion transport | 2.3E-02 |
| GO:0019216 | regulation of lipid metabolic process | 2.3E-02 |
| GO:0062012 | regulation of small molecule metabolic process | 2.3E-02 |
| GO:0071604 | transforming growth factor beta production | 2.5E-02 |
| GO:0003012 | muscle system process | 3.0E-02 |
| GO:0043648 | dicarboxylic acid metabolic process | 3.3E-02 |
| GO:0048732 | gland development | 4.2E-02 |
| GO:0062012 | regulation of small molecule metabolic process | 3.4E-02 |
| GO:0010959 | regulation of metal ion transport | 3.4E-02 |
| GO:0003012 | muscle system process | 3.4E-02 |
| GO:0048863 | stem cell differentiation | 4.5E-02 |

upregulated

| Gene Set | Description | FDR |
|---|---|---|
| GO:0002449 | lymphocyte mediated immunity | 2.2E-02 |
| GO:0002526 | acute inflammatory response | 2.2E-02 |
| GO:0050673 | epithelial cell proliferation | 2.2E-02 |
| GO:0010038 | response to metal ion | 2.2E-02 |
| GO:0002440 | production of molecular mediator of immune response | 2.2E-02 |
| GO:0098542 | defense response to other organism | 2.2E-02 |
| GO:0009112 | nucleobase metabolic process | 2.3E-02 |
| GO:0043062 | extracellular structure organization | 2.3E-02 |
| GO:0050663 | cytokine secretion | 5.0E-02 |

### HuH-7

downregulated

| Gene Set | Description | FDR |
|---|---|---|
| GO:0006260 | DNA replication | <2.2e-16 |
| GO:0044772 | mitotic cell cycle phase transition | 1.9E-11 |
| GO:0044843 | cell cycle G1/S phase transition | 1.5E-08 |
| GO:0000075 | cell cycle checkpoint | 7.7E-07 |
| GO:0071103 | DNA conformation change | 1.3E-06 |
| GO:0006310 | DNA recombination | 1.9E-06 |
| GO:0006302 | double-strand break repair | 3.2E-06 |
| GO:0022616 | DNA strand elongation | 1.8E-05 |

| | | |
|---|---|---|
| GO:0007059 | chromosome segregation | 3.0E-04 |
| GO:0044839 | cell cycle G2/M phase transition | 8.6E-04 |
| GO:0045930 | negative regulation of mitotic cell cycle | 1.0E-03 |
| GO:0006301 | postreplication repair | 1.3E-03 |
| GO:0048285 | organelle fission | 1.4E-03 |
| GO:0032200 | telomere organization | 1.4E-03 |
| GO:0051052 | regulation of DNA metabolic process | 2.1E-03 |
| GO:0006575 | cellular modified amino acid metabolic process | 2.6E-03 |
| GO:0071897 | DNA biosynthetic process | 4.2E-03 |
| GO:0071824 | protein-DNA complex subunit organization | 4.4E-03 |
| GO:0051321 | meiotic cell cycle | 4.4E-03 |
| GO:0042769 | DNA damage response, detection of DNA damage | 4.9E-03 |
| GO:1901987 | regulation of cell cycle phase transition | 5.4E-03 |
| GO:0006298 | mismatch repair | 5.4E-03 |
| GO:0061641 | CENP-A containing chromatin organization | 5.4E-03 |
| GO:0036297 | interstrand cross-link repair | 5.8E-03 |
| GO:0010948 | negative regulation of cell cycle process | 1.0E-02 |
| GO:0006333 | chromatin assembly or disassembly | 1.0E-02 |
| GO:0009314 | response to radiation | 1.0E-02 |
| GO:1904888 | cranial skeletal system development | 2.4E-02 |
| GO:0006284 | base-excision repair | 4.0E-02 |
| GO:0007568 | aging | 5.0E-02 |

### upregulated

| Gene Set | Description | FDR |
|---|---|---|
| GO:0016049 | cell growth | 1.9E-01 |
| GO:0098742 | cell-cell adhesion via plasma-membrane adhesion molecules | 1.9E-01 |
| GO:0070972 | protein localization to endoplasmic reticulum | 1.9E-01 |
| GO:0006605 | protein targeting | 1.9E-01 |
| GO:0006413 | translational initiation | 3.0E-01 |
| GO:0090150 | establishment of protein localization to membrane | 3.0E-01 |
| GO:0018208 | peptidyl-proline modification | 3.0E-01 |
| GO:0046486 | glycerolipid metabolic process | 5.9E-01 |
| GO:1904888 | cranial skeletal system development | 5.9E-01 |
| GO:0001525 | angiogenesis | 6.0E-01 |

**Table S5. Gene Set Enrichment analyses of transcriptome impacts of COP1 suppression in HuH-7, HepG2 and Hepatocytes.** Transcripts were mapped to Gene Ontology (geneontology_Biological_Process_noRedundant) and KEGG categories. Arranged by Normalized Effect Size. Hits common to more than one cell type are highlighted in red.

GO

**HuH-7**

| Gene Set | Description | NES | FDR |
|---|---|---|---|
| GO:0006260 | DNA replication | -4.4341 | 0 |
| GO:0044772 | mitotic cell cycle phase transition | -3.9099 | 0 |
| GO:0044839 | cell cycle G2/M phase transition | -3.519 | 0 |
| GO:0044843 | cell cycle G1/S phase transition | -3.382 | 0 |
| GO:1901987 | regulation of cell cycle phase transition | -3.2648 | 0 |
| GO:0007059 | chromosome segregation | -3.2585 | 0 |
| GO:0000075 | cell cycle checkpoint | -3.2432 | 0 |
| GO:0048285 | organelle fission | -3.1988 | 0 |
| GO:0071103 | DNA conformation change | -3.101 | 0 |
| GO:0010948 | negative regulation of cell cycle process | -3.0814 | 0 |
| GO:0061641 | CENP-A containing chromatin organizatic | -3.0146 | 0 |
| GO:0045930 | negative regulation of mitotic cell cycle | -3.0021 | 0 |
| GO:0051321 | meiotic cell cycle | -2.9098 | 0 |
| GO:0006302 | double-strand break repair | -2.792 | 6.1E-05 |
| GO:0022616 | DNA strand elongation | -2.7916 | 5.7E-05 |
| GO:0006310 | DNA recombination | -2.7283 | 5.4E-05 |
| GO:0006333 | chromatin assembly or disassembly | -2.6492 | 1.0E-04 |
| GO:0071824 | protein-DNA complex subunit organizatio | -2.5293 | 5.7E-04 |
| GO:0034502 | protein localization to chromosome | -2.4145 | 1.4E-03 |
| GO:0051052 | regulation of DNA metabolic process | -2.3984 | 1.8E-03 |
| GO:0070988 | demethylation | 2.2281 | 3.8E-02 |
| GO:0070972 | protein localization to endoplasmic reticul | 2.2503 | 3.6E-02 |
| GO:0032941 | secretion by tissue | 2.2572 | 4.2E-02 |
| GO:0098742 | cell-cell adhesion via plasma-membrane | 2.2906 | 4.2E-02 |
| GO:0044262 | cellular carbohydrate metabolic process | 2.3058 | 4.9E-02 |

**HepG2**

| Gene Set | Description | NES | FDR |
|---|---|---|---|
| GO:0044282 | small molecule catabolic process | -3.9555 | 0 |
| GO:0006631 | fatty acid metabolic process | -3.7902 | 0 |
| GO:0008202 | steroid metabolic process | -3.6309 | 0 |
| GO:0072376 | protein activation cascade | -3.6281 | 0 |
| GO:0055088 | lipid homeostasis | -3.4617 | 0 |
| GO:1901615 | organic hydroxy compound metabolic pro | -3.2933 | 0 |
| GO:0006091 | generation of precursor metabolites and | -3.2223 | 0 |
| GO:0016042 | lipid catabolic process | -3.2098 | 0 |
| GO:0006732 | coenzyme metabolic process | -3.142 | 0 |

| | | | |
|---|---|---|---|
| GO:0016053 | organic acid biosynthetic process | -3.1208 | 0 |
| GO:0010876 | lipid localization | -3.0669 | 0 |
| GO:0006638 | neutral lipid metabolic process | -3.0618 | 0 |
| GO:0016999 | antibiotic metabolic process | -2.98 | 0 |
| GO:0042180 | cellular ketone metabolic process | -2.9127 | 0 |
| GO:0006520 | cellular amino acid metabolic process | -2.8963 | 0 |
| GO:0002526 | acute inflammatory response | -2.8891 | 0 |
| GO:0015711 | organic anion transport | -2.8479 | 0 |
| GO:0033865 | nucleoside bisphosphate metabolic proce | -2.8237 | 0 |
| GO:0006575 | cellular modified amino acid metabolic pr | -2.7705 | 0 |
| GO:0015748 | organophosphate ester transport | -2.7459 | 0 |
| GO:0071825 | protein-lipid complex subunit organizatior | -2.6738 | 8.1E-05 |
| GO:0043574 | peroxisomal transport | -2.6361 | 1.1E-04 |
| GO:0009259 | ribonucleotide metabolic process | -2.5247 | 5.3E-04 |
| GO:0034976 | response to endoplasmic reticulum stress | -2.5032 | 5.8E-04 |
| GO:0042737 | drug catabolic process | -2.4131 | 1.3E-03 |
| GO:0070085 | glycosylation | -2.3514 | 2.4E-03 |
| GO:0009141 | nucleoside triphosphate metabolic proces | -2.2785 | 3.2E-03 |
| GO:0140053 | mitochondrial gene expression | -2.2455 | 3.6E-03 |
| GO:0010257 | NADH dehydrogenase complex assembly | -2.1179 | 8.2E-03 |
| GO:0044872 | lipoprotein localization | -2.1151 | 8.2E-03 |
| GO:0035456 | response to interferon-beta | -2.0008 | 1.6E-02 |
| GO:0072337 | modified amino acid transport | -1.933 | 2.3E-02 |
| GO:0006081 | cellular aldehyde metabolic process | -1.9249 | 2.4E-02 |
| GO:0046486 | glycerolipid metabolic process | -1.8947 | 2.7E-02 |
| GO:1903513 | endoplasmic reticulum to cytosol transpor | -1.8833 | 2.8E-02 |
| GO:0009743 | response to carbohydrate | -1.8483 | 3.3E-02 |
| GO:0001101 | response to acid chemical | -1.8227 | 3.7E-02 |
| GO:0046939 | nucleotide phosphorylation | -1.8 | 4.2E-02 |
| GO:0014075 | response to amine | 1.8837 | 4.3E-02 |
| GO:0050867 | positive regulation of cell activation | 1.8942 | 4.2E-02 |
| GO:0050906 | detection of stimulus involved in sensory | 1.911 | 3.9E-02 |
| GO:0007606 | sensory perception of chemical stimulus | 1.9188 | 3.8E-02 |
| GO:0003158 | endothelium development | 1.9657 | 3.1E-02 |
| GO:0045930 | negative regulation of mitotic cell cycle | 2.0053 | 2.6E-02 |
| GO:1902850 | microtubule cytoskeleton organization inv | 2.0089 | 2.5E-02 |
| GO:0021915 | neural tube development | 2.018 | 2.5E-02 |
| GO:0051052 | regulation of DNA metabolic process | 2.0768 | 2.0E-02 |
| GO:0048285 | organelle fission | 2.2032 | 1.2E-02 |
| GO:0001667 | ameboidal-type cell migration | 2.2073 | 1.2E-02 |
| GO:0051493 | regulation of cytoskeleton organization | 2.2178 | 1.1E-02 |
| GO:0071824 | protein-DNA complex subunit organizatio | 2.2369 | 1.0E-02 |
| GO:0090130 | tissue migration | 2.2429 | 1.0E-02 |
| GO:0042770 | signal transduction in response to DNA d | 2.2735 | 8.1E-03 |
| GO:0032970 | regulation of actin filament-based process | 2.3616 | 4.1E-03 |
| GO:0006302 | double-strand break repair | 2.3697 | 4.2E-03 |
| GO:0036297 | interstrand cross-link repair | 2.37 | 4.5E-03 |

| | | | |
|---|---|---|---|
| GO:2000147 | positive regulation of cell motility | 2.4025 | 3.6E-03 |
| GO:0031032 | actomyosin structure organization | 2.4038 | 3.8E-03 |
| GO:0061641 | CENP-A containing chromatin organizatic | 2.4162 | 3.7E-03 |
| GO:0032200 | telomere organization | 2.4563 | 3.2E-03 |
| GO:0071103 | DNA conformation change | 2.4564 | 3.5E-03 |
| GO:0006333 | chromatin assembly or disassembly | 2.474 | 3.3E-03 |
| GO:0000075 | cell cycle checkpoint | 2.4782 | 3.7E-03 |
| GO:0044772 | mitotic cell cycle phase transition | 2.5351 | 2.9E-03 |
| GO:0008544 | epidermis development | 2.5401 | 3.4E-03 |
| GO:0043588 | skin development | 2.5598 | 3.6E-03 |
| GO:0044843 | cell cycle G1/S phase transition | 2.6259 | 2.7E-03 |
| GO:0022616 | DNA strand elongation | 2.6869 | 1.8E-03 |
| GO:0007059 | chromosome segregation | 2.8819 | 5.3E-04 |
| GO:0006260 | DNA replication | 3.884 | 0.0E+00 |

**Hepatocytes**

| Gene Set | Description | NES | FDR |
|---|---|---|---|
| GO:0044282 | small molecule catabolic process | -2.5264 | 2.4E-02 |
| GO:0060840 | artery development | -2.4412 | 2.8E-02 |
| GO:0042180 | cellular ketone metabolic process | -2.3657 | 3.6E-02 |
| GO:0009410 | response to xenobiotic stimulus | -2.3339 | 3.4E-02 |
| GO:0032528 | microvillus organization | -2.3277 | 2.8E-02 |
| GO:0055081 | anion homeostasis | -2.2824 | 3.5E-02 |
| GO:0055088 | lipid homeostasis | -2.279 | 3.0E-02 |
| GO:0008202 | steroid metabolic process | -2.242 | 3.3E-02 |
| GO:0003012 | muscle system process | -2.0668 | 4.8E-02 |
| GO:0015849 | organic acid transport | -2.0603 | 4.8E-02 |
| GO:0009112 | nucleobase metabolic process | 2.2286 | 4.0E-02 |
| GO:0018208 | peptidyl-proline modification | 2.2422 | 4.9E-02 |
| GO:0006413 | translational initiation | 2.3506 | 2.9E-02 |
| GO:0070972 | protein localization to endoplasmic reticul | 3.4975 | 0.0E+00 |

KEGG

**HuH-7**

| Gene Set | Description | NES | FDR |
|---|---|---|---|
| hsa04110 | Cell cycle | -4.1587 | 0 |
| hsa03030 | DNA replication | -4.0391 | 0 |
| hsa03430 | Mismatch repair | -2.837 | 0 |
| hsa03050 | Proteasome | -2.8138 | 0 |
| hsa04914 | Progesterone-mediated oocyte maturatio | -2.6523 | 9.3E-04 |
| hsa03460 | Fanconi anemia pathway | -2.6116 | 9.7E-04 |
| hsa05322 | Systemic lupus erythematosus | -2.5167 | 1.7E-03 |
| hsa04114 | Oocyte meiosis | -2.4172 | 4.1E-03 |
| hsa03008 | Ribosome biogenesis in eukaryotes | -2.4085 | 4.3E-03 |
| hsa05169 | Epstein-Barr virus infection | -2.4005 | 3.9E-03 |

| | | | |
|---|---|---|---|
| hsa00480 | Glutathione metabolism | -2.3955 | 3.6E-03 |
| hsa03440 | Homologous recombination | -2.3754 | 3.7E-03 |
| hsa05416 | Viral myocarditis | -2.3409 | 4.6E-03 |
| hsa04721 | Synaptic vesicle cycle | -2.2871 | 5.9E-03 |
| hsa00980 | Metabolism of xenobiotics by cytochrome | -2.2784 | 5.8E-03 |
| hsa05204 | Chemical carcinogenesis | -2.1655 | 1.3E-02 |
| hsa04145 | Phagosome | -2.1559 | 1.3E-02 |
| hsa00190 | Oxidative phosphorylation | -2.0401 | 2.7E-02 |
| hsa05162 | Measles | -2.0083 | 3.2E-02 |
| hsa05203 | Viral carcinogenesis | -1.9927 | 3.4E-02 |
| hsa05110 | Vibrio cholerae infection | -1.9627 | 3.9E-02 |
| hsa05020 | Prion diseases | -1.9603 | 3.8E-02 |
| hsa04218 | Cellular senescence | -1.9399 | 4.2E-02 |
| hsa05034 | Alcoholism | -1.9267 | 4.4E-02 |
| hsa05330 | Allograft rejection | -1.9037 | 4.9E-02 |

**HepG2**

| Gene Set | Description | NES | FDR |
|---|---|---|---|
| hsa04610 | Complement and coagulation cascades | -3.7898 | 0 |
| hsa04975 | Fat digestion and absorption | -3.4699 | 0 |
| hsa03320 | PPAR signaling pathway | -3.4138 | 0 |
| hsa01100 | Metabolic pathways | -3.4032 | 0 |
| hsa04979 | Cholesterol metabolism | -3.1544 | 0 |
| hsa00100 | Steroid biosynthesis | -3.0108 | 0 |
| hsa01200 | Carbon metabolism | -2.8472 | 1.7E-04 |
| hsa00010 | Glycolysis / Gluconeogenesis | -2.8184 | 1.5E-04 |
| hsa00120 | Primary bile acid biosynthesis | -2.7835 | 1.3E-04 |
| hsa05204 | Chemical carcinogenesis | -2.7736 | 1.2E-04 |
| hsa00071 | Fatty acid degradation | -2.7688 | 1.1E-04 |
| hsa04146 | Peroxisome | -2.7052 | 2.0E-04 |
| hsa01212 | Fatty acid metabolism | -2.6842 | 1.8E-04 |
| hsa00980 | Metabolism of xenobiotics by cytochrome | -2.6762 | 2.5E-04 |
| hsa00190 | Oxidative phosphorylation | -2.6735 | 2.4E-04 |
| hsa01230 | Biosynthesis of amino acids | -2.6268 | 3.0E-04 |
| hsa05150 | Staphylococcus aureus infection | -2.5803 | 4.9E-04 |
| hsa04932 | Non-alcoholic fatty liver disease (NAFLD) | -2.5645 | 4.6E-04 |
| hsa00053 | Ascorbate and aldarate metabolism | -2.5204 | 6.9E-04 |
| hsa04152 | AMPK signaling pathway | -2.519 | 6.5E-04 |
| hsa04976 | Bile secretion | -2.4775 | 8.6E-04 |
| hsa04141 | Protein processing in endoplasmic reticul | -2.3154 | 2.2E-03 |
| hsa05012 | Parkinson disease | -2.258 | 3.4E-03 |
| hsa03050 | Proteasome | -2.2135 | 4.4E-03 |
| hsa03010 | Ribosome | -2.1588 | 5.6E-03 |
| hsa04714 | Thermogenesis | -2.1283 | 6.6E-03 |
| hsa04931 | Insulin resistance | -2.0972 | 8.1E-03 |
| hsa00052 | Galactose metabolism | -2.072 | 9.2E-03 |

| | | | |
|---|---|---|---|
| hsa00770 | Pantothenate and CoA biosynthesis | -2.0178 | 1.3E-02 |
| hsa00360 | Phenylalanine metabolism | -1.8007 | 4.5E-02 |
| hsa00510 | N-Glycan biosynthesis | -1.7803 | 4.6E-02 |
| hsa04810 | Regulation of actin cytoskeleton | 1.915 | 4.9E-02 |
| hsa04330 | Notch signaling pathway | 1.9599 | 3.9E-02 |
| hsa04392 | Hippo signaling pathway | 1.9612 | 4.1E-02 |
| hsa04740 | Olfactory transduction | 1.9686 | 4.2E-02 |
| hsa03420 | Nucleotide excision repair | 2.0662 | 2.4E-02 |
| hsa00531 | Glycosaminoglycan degradation | 2.0743 | 2.5E-02 |
| hsa04514 | Cell adhesion molecules (CAMs) | 2.0766 | 2.6E-02 |
| hsa03440 | Homologous recombination | 2.0983 | 2.5E-02 |
| hsa05202 | Transcriptional misregulation in cancer | 2.1005 | 2.7E-02 |
| hsa03430 | Mismatch repair | 2.1514 | 2.2E-02 |
| hsa05130 | Pathogenic Escherichia coli infection | 2.2067 | 1.7E-02 |
| hsa04218 | Cellular senescence | 2.2606 | 1.3E-02 |
| hsa04064 | NF-kappa B signaling pathway | 2.2664 | 1.5E-02 |
| hsa03460 | Fanconi anemia pathway | 2.4478 | 4.5E-03 |
| hsa03410 | Base excision repair | 2.4884 | 4.4E-03 |
| hsa04110 | Cell cycle | 2.6386 | 2.0E-03 |
| hsa03030 | DNA replication | 2.9101 | 8.2E-04 |

**Hepatocytes**

| Gene Set | Description | NES | FDR |
|---|---|---|---|
| hsa00980 | Metabolism of xenobiotics by cytochrome | -3.4527 | 0 |
| hsa00982 | Drug metabolism | -3.4458 | 0 |
| hsa00040 | Pentose and glucuronate interconversion | -3.3805 | 0 |
| hsa05204 | Chemical carcinogenesis | -3.3029 | 0 |
| hsa00053 | Ascorbate and aldarate metabolism | -3.0858 | 0 |
| hsa00983 | Drug metabolism | -3.039 | 0 |
| hsa00830 | Retinol metabolism | -2.9012 | 0 |
| hsa00140 | Steroid hormone biosynthesis | -2.689 | 3.3E-04 |
| hsa05206 | MicroRNAs in cancer | -2.657 | 4.4E-04 |
| hsa04928 | Parathyroid hormone synthesis, secretior | -2.2627 | 1.2E-02 |
| hsa04976 | Bile secretion | -2.1737 | 2.0E-02 |
| hsa04270 | Vascular smooth muscle contraction | -2.1682 | 1.9E-02 |
| hsa04964 | Proximal tubule bicarbonate reclamation | -2.1256 | 2.4E-02 |
| hsa00480 | Glutathione metabolism | -2.122 | 2.2E-02 |
| hsa04919 | Thyroid hormone signaling pathway | -2.1104 | 2.2E-02 |
| hsa04530 | Tight junction | -2.0869 | 2.5E-02 |
| hsa04915 | Estrogen signaling pathway | -2.0549 | 2.9E-02 |
| hsa00860 | Porphyrin and chlorophyll metabolism | -2.0045 | 4.0E-02 |
| hsa05219 | Bladder cancer | -2.0031 | 3.8E-02 |
| hsa03320 | PPAR signaling pathway | -1.9866 | 4.0E-02 |
| hsa00590 | Arachidonic acid metabolism | -1.9855 | 3.9E-02 |
| hsa01524 | Platinum drug resistance | -1.9552 | 4.5E-02 |
| hsa04024 | cAMP signaling pathway | -1.9547 | 4.3E-02 |

| | | | |
|---|---|---|---|
| hsa00531 | Glycosaminoglycan degradation | 2.1481 | 4.6E-02 |
| hsa04978 | Mineral absorption | 2.6509 | 1.4E-03 |
| hsa03010 | Ribosome | 3.4594 | 0.00 |

**Table S6. Ingenuity Canonical Pathways identified in COP1 suppressed cells** . Only nominally significant hits (green) in at least one cell type are shown. Values are organized as increasing Z-scores, using Hepatocyte values. N/A: no Z-score available for the pathway.

| | Hepatocytes | HepG2 | HuH7 |
|---|---|---|---|
| Xenobiotic Metabolism General Signaling Pathway | -2.921 | -3.78 | -0.333 |
| Xenobiotic Metabolism PXR Signaling Pathway | -2.655 | -5.425 | -0.5 |
| Xenobiotic Metabolism CAR Signaling Pathway | -2.596 | -4.439 | -1 |
| SPINK1 General Cancer Pathway | -2.324 | -2.111 | -1.342 |
| Complement System | -2.121 | -1.941 | N/A |
| Methylglyoxal Degradation III | -2 | -2 | N/A |
| CCR3 Signaling in Eosinophils | -1.698 | 0.832 | 0.447 |
| Bile Acid Biosynthesis, Neutral Pathway | -1.633 | -2.828 | N/A |
| Pregnenolone Biosynthesis | -1.633 | N/A | N/A |
| Nicotine Degradation II | -1.606 | -1.291 | N/A |
| Acetone Degradation I (to Methylglyoxal) | -1.508 | -0.333 | N/A |
| Aryl Hydrocarbon Receptor Signaling | -1.5 | 0.728 | -1.387 |
| LXR/RXR Activation | -1.461 | -5.096 | 0 |
| Nicotine Degradation III | -1.414 | -1.069 | N/A |
| Melatonin Degradation I | -1.342 | -1.291 | -1 |
| Integrin Signaling | -1.257 | 3.703 | 0.258 |
| Thyroid Hormone Metabolism II (via Conjugation and/or Degrada | -1.155 | -1.89 | N/A |
| Histidine Degradation VI | -1.134 | -1 | N/A |
| PXR/RXR Activation | -1.091 | -1.877 | -1.265 |
| Bupropion Degradation | -1 | -0.333 | N/A |
| STAT3 Pathway | -0.962 | 0.6 | -0.302 |
| Macropinocytosis Signaling | -0.905 | 2.309 | N/A |
| PPARα/RXRα Activation | -0.898 | -1.89 | -0.905 |
| Regulation Of The Epithelial Mesenchymal Transition By Growth | -0.73 | 1.768 | 0.229 |
| NRF2-mediated Oxidative Stress Response | -0.728 | -3.266 | 0.378 |
| Endothelin-1 Signaling | -0.707 | 0.192 | -0.775 |
| Superpathway of Melatonin Degradation | -0.626 | -1.698 | -0.447 |
| Estrogen Biosynthesis | -0.577 | -0.577 | 0 |
| Ephrin B Signaling | -0.302 | 0 | N/A |
| Serotonin Degradation | -0.229 | -3.051 | N/A |
| Ephrin Receptor Signaling | -0.209 | 1 | 1.265 |
| Kinetochore Metaphase Signaling Pathway | 0 | 1.387 | -1 |
| Oxidative Ethanol Degradation III | 0 | -1.342 | N/A |
| Regulation of Cellular Mechanics by Calpain Protease | 0 | 1.291 | 0.816 |
| Retinol Biosynthesis | 0 | -1.732 | N/A |
| Glucose and Glucose-1-phosphate Degradation | 0.447 | N/A | N/A |
| IL-6 Signaling | 0.6 | -0.408 | 0.378 |
| p38 MAPK Signaling | 0.626 | 0.229 | -0.5 |
| Dopamine Degradation | 0.707 | -2.236 | N/A |
| Phosphatidylglycerol Biosynthesis II (Non-plastidic) | 0.707 | 1 | N/A |
| Purine Nucleotides Degradation II (Aerobic) | 0.816 | N/A | N/A |

| | | | |
|---|---|---|---|
| Tryptophan Degradation X (Mammalian, via Tryptamine) | 0.816 | -2.236 | N/A |
| LPS/IL-1 Mediated Inhibition of RXR Function | 1.061 | 2.2 | 0.333 |
| BER (Base Excision Repair) Pathway | 1.342 | 2.496 | -1.508 |
| Guanosine Nucleotides Degradation III | 1.342 | N/A | N/A |
| Coagulation System | 1.414 | 0 | -1 |
| Acute Phase Response Signaling | 1.569 | -0.822 | 0.832 |
| Zymosterol Biosynthesis | 2 | N/A | N/A |
| Superpathway of Cholesterol Biosynthesis | 2.121 | -3.357 | N/A |
| Cholesterol Biosynthesis I | 2.449 | -2.121 | N/A |
| Cholesterol Biosynthesis II (via 24,25-dihydrolanosterol) | 2.449 | -2.121 | N/A |
| Cholesterol Biosynthesis III (via Desmosterol) | 2.449 | -2.121 | N/A |
| Osteoarthritis Pathway | 3.042 | 2.469 | -1.414 |
| α-tocopherol Degradation | N/A | N/A | N/A |
| Atherosclerosis Signaling | N/A | N/A | N/A |
| Cell Cycle Control of Chromosomal Replication | N/A | 3.578 | -4.583 |
| Citrulline Biosynthesis | N/A | 0 | N/A |
| FXR/RXR Activation | N/A | N/A | N/A |
| Germ Cell-Sertoli Cell Junction Signaling | N/A | N/A | N/A |
| Hepatic Cholestasis | N/A | N/A | N/A |
| IL-12 Signaling and Production in Macrophages | N/A | N/A | N/A |
| Melatonin Degradation II | N/A | N/A | N/A |
| Mevalonate Pathway I | N/A | -2.646 | N/A |
| Mismatch Repair in Eukaryotes | N/A | N/A | N/A |
| Phenylalanine Degradation I (Aerobic) | N/A | 0 | N/A |
| Role of BRCA1 in DNA Damage Response | N/A | 1 | -1.508 |
| Role of Tissue Factor in Cancer | N/A | N/A | N/A |
| Xenobiotic Metabolism Signaling | N/A | N/A | N/A |

**Table S7. Upstream regulators predicted in at least one cell type by IPA upstream regulators**. Bias-corrected Z-scores are shown incrementally, using hepatocyte values. Negative values, predicted inhibition; positive values, predicted activation. N/A, no Z-score available.

| | Hepatocytes | HepG2 | HuH7 |
|---|---|---|---|
| HNF1A | -3.15 | -5.388 | 1.538 |
| estrogen receptor | -3.035 | -1.966 | N/A |
| Growth hormone | -2.976 | -1.114 | N/A |
| MRTFB | -2.913 | 1.73 | N/A |
| ADRB | -2.733 | 2.014 | N/A |
| NOX4 | -2.601 | -1.075 | N/A |
| PPARGC1A | -2.572 | -4.296 | -0.388 |
| OGA | -2.536 | -0.126 | N/A |
| F2 | -2.523 | 2.173 | N/A |
| SNAI2 | -2.496 | 0.329 | N/A |
| Tgf beta | -2.474 | 0.877 | N/A |
| HNF4alpha dimer | -2.429 | -2.583 | N/A |
| HSPA5 | -2.418 | -1.114 | N/A |
| BRD4 | -2.395 | 1.343 | -1.515 |
| SRF | -2.378 | 1.947 | -2.563 |
| E2F1 | -2.288 | 3.318 | -4.71 |
| RASSF1 | -2.253 | -0.211 | N/A |
| MRTFA | -2.236 | 1.693 | -1.308 |
| TFAM | -2.236 | -1.667 | N/A |
| PROM1 | -2.236 | 1.334 | N/A |
| MOGAT1 | -2.213 | -2.213 | N/A |
| ACOX1 | -2.162 | 0.071 | 0.258 |
| IL10RA | -2.147 | 0.215 | -1.789 |
| PPARG | -2.129 | -4.328 | 0.063 |
| GSTO1 | -2.124 | -0.759 | N/A |
| Alpha catenin | -2.117 | -1.809 | N/A |
| RUNX1 | -2.071 | -0.936 | N/A |
| EDN1 | -2.057 | 2.789 | N/A |
| NR5A1 | -2.049 | -1.246 | -0.113 |
| TGFB1 | -2.04 | 5.721 | 1.675 |
| ERG | -2.028 | 3.334 | N/A |
| PKD1 | -2.024 | -2.805 | -2.901 |
| RAC1 | -1.945 | 2.16 | N/A |
| FAS | -1.82 | 1.331 | -2.919 |
| PXR ligand-PXR-Retinoic acid-RXRA | -1.798 | -2.112 | -0.489 |
| PNPLA2 | -1.794 | -2.854 | N/A |
| INSIG1 | -1.777 | 2.388 | N/A |
| TGFB3 | -1.759 | 2.892 | -0.307 |
| CYP1B1 | -1.755 | N/A | -2.219 |
| NFE2L2 | -1.736 | -4.51 | N/A |

| | | | |
|---|---|---|---|
| EDN3 | -1.664 | 2.377 | -0.152 |
| HNF4A | -1.662 | -6.356 | 0.792 |
| GATA1 | -1.57 | N/A | 2.421 |
| BMP4 | -1.566 | 0.278 | 2.248 |
| IGFBP2 | -1.535 | 2.412 | -0.758 |
| SPARC | -1.513 | -3.343 | 1.245 |
| NFYA | -1.405 | -2.561 | N/A |
| RXRA | -1.358 | -2.107 | N/A |
| PRKCD | -1.313 | 1.35 | 2.113 |
| TP53 | -1.242 | 1.182 | 3.445 |
| PPARD | -1.173 | -2.878 | N/A |
| N-cor | -1.108 | 2.219 | N/A |
| RBL1 | -1.048 | -3.482 | 2.723 |
| ERBB4 | -1.033 | -0.363 | -2.781 |
| PI3K (family) | -0.819 | -0.901 | -2.908 |
| LPCAT3 | -0.816 | -2 | N/A |
| YAP1 | -0.812 | 3.7 | -0.751 |
| EGLN | -0.806 | 2.881 | -2.798 |
| FOXA1 | -0.762 | -2.047 | -1.074 |
| CCND1 | -0.75 | 4.397 | -4.599 |
| NTRK2 | -0.714 | 2.065 | -1.474 |
| Immunoglobulin | -0.705 | 0.585 | -2.061 |
| IRF2 | -0.675 | -2.095 | 1.682 |
| ERK1/2 | -0.673 | 3.051 | 0.112 |
| VCAN | -0.647 | -0.626 | -3.448 |
| E2F3 | -0.62 | 3.687 | -3.576 |
| INSR | -0.572 | -3.708 | N/A |
| HNF1B | -0.57 | -3.127 | N/A |
| KDM1A | -0.561 | 2.179 | -2.45 |
| GLI1 | -0.529 | -1.601 | -3.684 |
| FN1 | -0.468 | 2.51 | N/A |
| RELA | -0.461 | 2.309 | N/A |
| let-7 | -0.441 | -3.197 | 3.917 |
| KLF5 | -0.441 | 2.282 | -1.76 |
| ESR1 | -0.402 | 3.03 | -1.693 |
| ERK | -0.401 | 3.305 | N/A |
| VEGFA | -0.388 | 2.026 | -2.222 |
| THRB | -0.347 | -2.043 | N/A |
| mir-21 | -0.332 | -2.03 | 1.919 |
| BTG2 | -0.328 | -2.54 | 0.238 |
| ERBB2 | -0.305 | 3.579 | -6.116 |
| NCOA2 | -0.284 | -2.07 | N/A |
| VDR | -0.277 | 2.404 | -1.55 |
| Gcg | -0.258 | -1.604 | 2.236 |
| HGF | -0.213 | 3.723 | -1.841 |
| SNAI1 | -0.191 | 1.424 | -2.003 |
| miR-155-5p (miRNAs w/seed UAAUGCU | -0.15 | -2.587 | 1.53 |

| | | | |
|---|---|---|---|
| IL2 | -0.121 | 2.919 | -2.15 |
| BSG | -0.109 | 2.175 | -0.579 |
| FGF2 | -0.088 | 2.133 | 1.077 |
| miR-199a-5p (and other miRNAs w/seec | -0.057 | -2.428 | N/A |
| POR | 0.07 | 2.686 | 0.724 |
| FXR ligand-FXR-Retinoic acid-RXR? | 0.098 | -2.447 | N/A |
| CD36 | 0.108 | 1.74 | 2.095 |
| EZH2 | 0.122 | 2.147 | N/A |
| S100A9 | 0.142 | 1.092 | -3.309 |
| IRF1 | 0.145 | -2.302 | -1.942 |
| PRL | 0.16 | -1.264 | -2.663 |
| HDAC1 | 0.206 | 0.457 | 2.998 |
| CDKN2A | 0.215 | -3.29 | 4.203 |
| CSF2 | 0.216 | 3.361 | -2.182 |
| IRF3 | 0.241 | -0.509 | -2.659 |
| FOXO4 | 0.269 | -0.226 | 2.09 |
| IFNB1 | 0.339 | -2.403 | -0.876 |
| SMARCB1 | 0.346 | -2.18 | 3.053 |
| MGP | 0.378 | 2.354 | N/A |
| EFNA1 | 0.385 | -2.034 | N/A |
| SP110 | 0.392 | -2.661 | N/A |
| mir-8 | 0.413 | -3.113 | 1.526 |
| NCOR1 | 0.426 | 2.35 | 1.863 |
| EHMT1 | 0.426 | -1.301 | 2.111 |
| RPTOR | 0.445 | -2.548 | N/A |
| TAB1 | 0.454 | 2.216 | 0.945 |
| PDLIM2 | 0.469 | -0.871 | -2.309 |
| S100A8 | 0.478 | 1.37 | -2.719 |
| Vegf | 0.491 | 3.789 | -2.442 |
| SYVN1 | 0.508 | 2.057 | N/A |
| JUN | 0.527 | 2.384 | N/A |
| IFN Beta | 0.533 | -1.505 | -2.248 |
| SOX17 | 0.563 | -2.155 | N/A |
| RORA | 0.567 | 0.279 | -2.17 |
| RAF1 | 0.622 | 3.092 | -1.172 |
| WT1 | 0.728 | 1.75 | 2.151 |
| MAP3K7 | 0.816 | 2.117 | -1.019 |
| STAT1/3/5 dimer | 0.816 | 2.236 | N/A |
| FOSL1 | 0.827 | 4.16 | N/A |
| TGFA | 0.893 | 2.542 | N/A |
| ESR2 | 0.9 | 2.452 | -1.305 |
| SLC27A2 | 0.91 | 3.707 | N/A |
| SOX4 | 0.966 | 3.998 | N/A |
| CCL5 | 1.017 | 0.388 | -2.386 |
| CEBPB | 1.05 | 2.598 | -4.085 |
| SREBF1 | 1.093 | -3.883 | N/A |
| STAT3 | 1.176 | 0.004 | 2.516 |

| | | | |
|---|---|---|---|
| PDGF BB | 1.179 | 2.189 | N/A |
| RABL6 | 1.192 | 2.683 | -3.889 |
| RUNX2 | 1.212 | 0.321 | 2.54 |
| Mek | 1.282 | 1.539 | -2.036 |
| DDX58 | 1.29 | -2.795 | -1.893 |
| NGFR | 1.359 | -2.06 | 2.193 |
| ELOVL3 | 1.387 | 2.478 | N/A |
| SASH1 | 1.414 | N/A | -2.333 |
| HMOX1 | 1.426 | -2.462 | 2.939 |
| MAPK7 | 1.482 | -3.193 | N/A |
| MAP2K1/2 | 1.53 | 0.488 | 2.415 |
| KLF6 | 1.615 | 2.33 | N/A |
| MAVS | 1.621 | -2.675 | -2.201 |
| GSR | 1.667 | 3.357 | N/A |
| CEBPA | 1.68 | -2.782 | 1.303 |
| TXNRD1 | 1.706 | 3.503 | N/A |
| EGF | 1.754 | 2.165 | N/A |
| IL17A | 1.763 | 2.199 | N/A |
| NUPR1 | 1.889 | -1.714 | 5.288 |
| Nr1h | 1.918 | -2.765 | N/A |
| MAP2K5 | 1.947 | -3.095 | -0.342 |
| IL1A | 1.997 | 2.816 | N/A |
| TLR3 | 2.015 | -0.156 | N/A |
| NR3C1 | 2.066 | -1.007 | 1.928 |
| GPER1 | 2.074 | -0.428 | N/A |
| INPP5D | 2.1 | N/A | N/A |
| TNF | 2.109 | 2.855 | 1.085 |
| LUCAT1 | 2.121 | -0.535 | 1.89 |
| HFE | 2.132 | -2.219 | N/A |
| SCAP | 2.145 | -4.227 | N/A |
| CSF1 | 2.191 | -1.064 | N/A |
| GATA2 | 2.206 | N/A | N/A |
| NCOA1 | 2.275 | -1.303 | N/A |
| TARDBP | 2.36 | -2.412 | N/A |
| F3 | 2.421 | 2.67 | -0.816 |
| GCK | 2.425 | 1.117 | N/A |
| CSF | 2.517 | -0.218 | N/A |
| IL1 | 2.541 | 2.743 | 0.645 |
| SH3TC2 | 2.646 | -2.673 | N/A |
| BHLHE40 | 2.68 | -0.2 | 0.104 |
| SREBF2 | 2.745 | -3.521 | N/A |
| IL6 | 3.472 | 0.342 | 2.57 |
| IL1B | 3.553 | 1.373 | 0.888 |
| E2f | N/A | 4.41 | -4.185 |
| TBX2 | N/A | 4.168 | -3.897 |
| CDKN1A | N/A | -2.875 | 4.399 |
| Rb | N/A | -3.455 | 3.395 |

| | | | |
|---|---|---|---|
| RB1 | N/A | -3.18 | 2.99 |
| EP400 | N/A | 2.985 | -3.162 |
| MAP4K4 | N/A | 4.034 | -2 |
| IFNL1 | N/A | -2.506 | -3.242 |
| BNIP3L | N/A | -2.412 | 2.72 |
| E2F6 | N/A | -1.897 | 3.051 |
| FOXM1 | N/A | 3.493 | -1.405 |
| E2F2 | N/A | 2.644 | -1.667 |
| ERN1 | N/A | -1.715 | 2.433 |
| SIX2 | N/A | -1.789 | 2.121 |
| ACSS2 | N/A | -3.827 | N/A |
| EIF4G1 | N/A | 2.353 | -1.342 |
| DYRK1A | N/A | -2.279 | 1.353 |
| NKX2-2-AS1 | N/A | -3.606 | N/A |
| PPARGC1B | N/A | -3.523 | N/A |
| BACH1 | N/A | 3.498 | N/A |
| MFSD2A | N/A | 3.426 | N/A |
| PTN | N/A | -3.418 | N/A |
| TRIB3 | N/A | 1.039 | 2.177 |
| ATP7B | N/A | -3.207 | N/A |
| AIRE | N/A | -1 | 2 |
| H2AZ1 | N/A | 0.956 | -2 |
| COMMD1 | N/A | 2.891 | N/A |
| IGF2 | N/A | -2.333 | -0.555 |
| CITED2 | N/A | -2.011 | 0.842 |
| XDH | N/A | 2.793 | N/A |
| ATF4 | N/A | -2.747 | N/A |
| let-7a-5p (and other miRNAs w/seed GA | N/A | -2.659 | N/A |
| MIF | N/A | 2.633 | N/A |
| Irgm1 | N/A | -0.316 | 2.274 |
| ELAVL1 | N/A | -0.36 | -2.179 |
| ZFP36 | N/A | -2.483 | N/A |
| IFNL4 | N/A | N/A | -2.421 |
| ETS1 | N/A | 2.363 | N/A |
| C4BP | N/A | -2.236 | N/A |
| TLR7 | N/A | 2.236 | N/A |
| CHCHD5 | N/A | -2.236 | N/A |
| AGO2 | N/A | 2.233 | N/A |
| TGIF2 | N/A | 2.224 | N/A |
| SOD1 | N/A | -2.2 | N/A |
| CXCR4 | N/A | 2.113 | N/A |
| Laminin (complex) | N/A | 2.096 | N/A |
| ALDH3A2 | N/A | 2 | N/A |
| PPP2R2A | N/A | 2 | N/A |

**Table S8. Over-representation analysis of COP1 and HNF4A targeted hits.** Analysis performed via WebGestalt, mapping shared FDR significant hits to enriched KEGG, GO and reactome gene sets. Only FDR significant hits are shown. Values are organized by decreasing enrichments ratios, to highlight relative biological contribution. Size, Gene Set Size; Expect: Expected value by chance (number of hits mapped to both input and geneset); Ratio: Enrichment ratio (number of actual hits mapped to input and geneset overlap vs number of hits predicted by chance). P value, nominal pvalue of overlap. FDR, False Discovery Rate.

| GO (Biological_Process_noRedundant) | | | | | | |
|---|---|---|---|---|---|---|
| Gene Set | Description | Size | Expect | Ratio | P Value | FDR |
| GO:0044282 | small molecule catabolic process | 401 | 5.40 | 4.45 | 8.8E-10 | 7.4E-07 |
| GO:0015711 | organic anion transport | 431 | 5.80 | 3.79 | 8.1E-08 | 3.5E-05 |
| GO:0055088 | lipid homeostasis | 116 | 1.56 | 7.05 | 4.5E-07 | 1.3E-04 |
| GO:0010876 | lipid localization | 343 | 4.62 | 3.90 | 8.9E-07 | 1.9E-04 |
| GO:0070482 | response to oxygen levels | 321 | 4.32 | 3.94 | 1.6E-06 | 2.7E-04 |
| GO:0016042 | lipid catabolic process | 295 | 3.97 | 4.03 | 2.4E-06 | 3.4E-04 |
| GO:0072376 | protein activation cascade | 90 | 1.21 | 7.43 | 3.3E-06 | 3.8E-04 |
| GO:0002526 | acute inflammatory response | 143 | 1.92 | 5.72 | 3.6E-06 | 3.8E-04 |
| GO:0001101 | response to acid chemical | 309 | 4.16 | 3.85 | 4.4E-06 | 4.1E-04 |
| GO:0043627 | response to estrogen | 71 | 0.96 | 8.37 | 4.8E-06 | 4.1E-04 |
| GO:0008202 | steroid metabolic process | 288 | 3.88 | 3.87 | 8.2E-06 | 6.3E-04 |
| GO:0006631 | fatty acid metabolic process | 329 | 4.43 | 3.61 | 9.7E-06 | 6.8E-04 |
| GO:0042180 | cellular ketone metabolic process | 170 | 2.29 | 4.81 | 1.9E-05 | 1.2E-03 |
| GO:1901615 | organic hydroxy compound metabolic process | 474 | 6.38 | 2.98 | 2.2E-05 | 1.3E-03 |
| GO:0043574 | peroxisomal transport | 66 | 0.89 | 7.88 | 2.8E-05 | 1.6E-03 |
| GO:0019216 | regulation of lipid metabolic process | 360 | 4.84 | 3.30 | 2.9E-05 | 1.6E-03 |
| GO:0006979 | response to oxidative stress | 406 | 5.46 | 3.11 | 3.5E-05 | 1.7E-03 |
| GO:0051348 | negative regulation of transferase activity | 251 | 3.38 | 3.85 | 3.5E-05 | 1.7E-03 |
| GO:0006732 | coenzyme metabolic process | 336 | 4.52 | 3.32 | 5.0E-05 | 2.2E-03 |
| GO:0006520 | cellular amino acid metabolic process | 300 | 4.04 | 3.47 | 5.5E-05 | 2.3E-03 |
| GO:0016053 | organic acid biosynthetic process | 380 | 5.11 | 3.13 | 5.6E-05 | 2.3E-03 |
| GO:0015850 | organic hydroxy compound transport | 227 | 3.05 | 3.93 | 5.8E-05 | 2.3E-03 |
| GO:0043062 | extracellular structure organization | 385 | 5.18 | 3.09 | 6.5E-05 | 2.3E-03 |
| GO:0009636 | response to toxic substance | 470 | 6.33 | 2.85 | 6.6E-05 | 2.3E-03 |

| | | | | | | |
|---|---|---|---|---|---|---|
| GO:0007031 | peroxisome organization | 77 | 1.04 | 6.76 | 7.8E-05 | 2.6E-03 |
| GO:0097006 | regulation of plasma lipoprotein particle levels | 78 | 1.05 | 6.67 | 8.4E-05 | 2.8E-03 |
| GO:0006790 | sulfur compound metabolic process | 333 | 4.48 | 3.12 | 1.7E-04 | 5.3E-03 |
| GO:0015849 | organic acid transport | 296 | 3.98 | 3.26 | 1.9E-04 | 5.7E-03 |
| GO:0010038 | response to metal ion | 339 | 4.56 | 3.07 | 2.0E-04 | 5.9E-03 |
| GO:0046677 | response to antibiotic | 299 | 4.02 | 3.23 | 2.1E-04 | 5.9E-03 |
| GO:0051604 | protein maturation | 307 | 4.13 | 3.15 | 2.7E-04 | 7.3E-03 |
| GO:0002791 | regulation of peptide secretion | 439 | 5.91 | 2.71 | 3.0E-04 | 7.8E-03 |
| GO:0071825 | protein-lipid complex subunit organization | 45 | 0.61 | 8.26 | 3.3E-04 | 8.5E-03 |
| GO:0061008 | hepaticobiliary system development | 133 | 1.79 | 4.47 | 4.3E-04 | 1.1E-02 |
| GO:0062012 | regulation of small molecule metabolic process | 324 | 4.36 | 2.98 | 4.5E-04 | 1.1E-02 |
| GO:0048732 | gland development | 414 | 5.57 | 2.69 | 4.9E-04 | 1.1E-02 |
| GO:0071706 | tumor necrosis factor superfamily cytokine produc | 136 | 1.83 | 4.37 | 5.0E-04 | 1.1E-02 |
| GO:0034367 | protein-containing complex remodeling | 28 | 0.38 | 10.62 | 5.1E-04 | 1.1E-02 |
| GO:0009743 | response to carbohydrate | 213 | 2.87 | 3.49 | 6.2E-04 | 1.4E-02 |
| GO:0005996 | monosaccharide metabolic process | 257 | 3.46 | 3.18 | 7.3E-04 | 1.5E-02 |
| GO:0016052 | carbohydrate catabolic process | 150 | 2.02 | 3.96 | 9.6E-04 | 1.8E-02 |
| GO:0033500 | carbohydrate homeostasis | 222 | 2.99 | 3.01 | 3.1E-03 | 4.5E-02 |
| GO:0061458 | reproductive system development | 406 | 5.46 | 2.38 | 3.4E-03 | 4.7E-02 |
| GO:0006979 | response to oxidative stress | 406 | 5.46 | 3.11 | 3.5E-05 | 1.7E-03 |
| GO:0043062 | extracellular structure organization | 385 | 5.18 | 3.09 | 6.5E-05 | 2.3E-03 |
| GO:0010038 | response to metal ion | 339 | 4.56 | 3.07 | 2.0E-04 | 5.9E-03 |
| GO:0033500 | carbohydrate homeostasis | 222 | 2.99 | 3.01 | 3.1E-03 | 4.5E-02 |
| GO:0030258 | lipid modification | 274 | 3.69 | 2.98 | 1.2E-03 | 2.1E-02 |
| GO:0062012 | regulation of small molecule metabolic process | 324 | 4.36 | 2.98 | 4.5E-04 | 1.1E-02 |
| GO:1901615 | organic hydroxy compound metabolic process | 474 | 6.38 | 2.98 | 2.2E-05 | 1.3E-03 |
| GO:0009914 | hormone transport | 308 | 4.15 | 2.90 | 9.6E-04 | 1.8E-02 |
| GO:0009636 | response to toxic substance | 470 | 6.33 | 2.85 | 6.6E-05 | 2.3E-03 |
| GO:0051188 | cofactor biosynthetic process | 264 | 3.55 | 2.81 | 3.1E-03 | 4.5E-02 |
| GO:0050727 | regulation of inflammatory response | 348 | 4.68 | 2.78 | 8.7E-04 | 1.8E-02 |
| GO:0002791 | regulation of peptide secretion | 439 | 5.91 | 2.71 | 3.0E-04 | 7.8E-03 |
| GO:0048732 | gland development | 414 | 5.57 | 2.69 | 4.9E-04 | 1.1E-02 |
| GO:0048545 | response to steroid hormone | 364 | 4.90 | 2.65 | 1.3E-03 | 2.2E-02 |
| GO:0042326 | negative regulation of phosphorylation | 395 | 5.32 | 2.63 | 9.3E-04 | 1.8E-02 |

| | | | | | | |
|---|---|---|---|---|---|---|
| GO:0052547 | regulation of peptidase activity | 398 | 5.36 | 2.61 | 1.0E-03 | 1.8E-02 |
| GO:0051047 | positive regulation of secretion | 388 | 5.22 | 2.49 | 2.3E-03 | 3.5E-02 |
| GO:0050878 | regulation of body fluid levels | 458 | 6.16 | 2.43 | 1.4E-03 | 2.3E-02 |
| GO:0061458 | reproductive system development | 406 | 5.46 | 2.38 | 3.4E-03 | 4.7E-02 |

REACTOME

| Gene Set | Description | Size | Expect | Ratio | P Value | FDR |
|---|---|---|---|---|---|---|
| R-HSA-1430728 | Metabolism | 1957 | 30.49 | 2.00 | 1.0E-08 | 1.7E-05 |
| R-HSA-192105 | Synthesis of bile acids and bile salts | 32 | 0.50 | 14.04 | 4.7E-07 | 4.0E-04 |
| R-HSA-193368 | Synthesis of bile acids and bile salts via 7alpha-hydrox | 22 | 0.34 | 17.50 | 7.9E-07 | 4.5E-04 |
| R-HSA-194068 | Bile acid and bile salt metabolism | 41 | 0.64 | 10.96 | 2.8E-06 | 1.2E-03 |
| R-HSA-977606 | Regulation of Complement cascade | 45 | 0.70 | 9.98 | 5.4E-06 | 1.8E-03 |
| R-HSA-166658 | Complement cascade | 56 | 0.87 | 8.02 | 2.4E-05 | 5.9E-03 |
| R-HSA-211945 | Phase I - Functionalization of compounds | 100 | 1.56 | 5.78 | 2.4E-05 | 5.9E-03 |
| R-HSA-9033241 | Peroxisomal protein import | 60 | 0.93 | 7.49 | 3.8E-05 | 8.0E-03 |
| R-HSA-1474228 | Degradation of the extracellular matrix | 136 | 2.12 | 4.72 | 5.0E-05 | 9.5E-03 |
| R-HSA-174577 | Activation of C3 and C5 | 8 | 0.12 | 24.07 | 2.0E-04 | 3.0E-02 |
| R-HSA-166665 | Terminal pathway of complement | 8 | 0.12 | 24.07 | 2.0E-04 | 3.0E-02 |
| R-HSA-211859 | Biological oxidations | 195 | 3.04 | 3.62 | 2.3E-04 | 3.2E-02 |
| R-HSA-8957275 | Post-translational protein phosphorylation | 106 | 1.65 | 4.84 | 2.4E-04 | 3.2E-02 |

KEGG

| Gene Set | Description | Size | Expect | Ratio | P Value | FDR |
|---|---|---|---|---|---|---|
| hsa04610 | Complement and coagulation cascades | 79 | 1.41 | 7.10 | 1.1E-06 | 3.7E-04 |
| hsa00120 | Primary bile acid biosynthesis | 16 | 0.29 | 17.53 | 6.2E-06 | 1.0E-03 |
| hsa03320 | PPAR signaling pathway | 67 | 1.19 | 5.86 | 1.7E-04 | 1.9E-02 |
| hsa04146 | Peroxisome | 80 | 1.43 | 4.91 | 5.2E-04 | 3.5E-02 |
| hsa04726 | Serotonergic synapse | 105 | 1.87 | 4.27 | 5.3E-04 | 3.5E-02 |
| hsa04950 | Maturity onset diabetes of the young | 25 | 0.45 | 8.98 | 9.1E-04 | 4.9E-02 |

**Table S9. Comparison of Gene list from Lee et al (Suppl Table 2) with values from current study**. 78 Gene list from Lee et al, describing genes impacted similarly in HuH-7 and HepG2 was converted to a 74 gene list via the Transcriptome Analysis Console. Values from Lee et al are available at the Gene Expression Omnibus dataset GSE21955. Upregulated and downregulated genes in response to COP1 suppression are highlighted in red and green respectively.

| **HepG2 (current)** | | | **HepG2 (Lee et al)** | | |
|---|---|---|---|---|---|
| Gene Symbol | Fold Chan | pValue | Gene Symbol | Fold Change (Log2) | pValue |
| AARS | -1.07 | 0.5923 | AARS | -1.024 | 0 |
| ABL1 | 1 | 0.3505 | ABL1 | -1.307 | 0.0004 |
| ALDH18A1 | -1.02 | 0.9913 | ALDH18A1 | -1.268 | 0.0006 |
| ARIH2 | -1.05 | 0.9615 | ARIH2 | -1.064 | 0.0018 |
| ARIH2 | -1.08 | 0.3728 | | | |
| ARIH2 | -1.01 | 0.4357 | | | |
| B4GALT5 | 1.59 | 0.0006 | B4GALT5 | -1.214 | 0.0014 |
| BOLA3 | -1.13 | 0.098 | BOLA3 | -1.746 | 0 |
| C6orf106 | -1.3 | 0.0247 | C6orf106 | -1.287 | 0 |
| C17orf58 | -1.08 | 0.2769 | C17orf58 | -1.515 | 0.0002 |
| CANX | -1.03 | 0.8011 | CANX | -3.516 | 0.0006 |
| CCNI | 1.13 | 0.2479 | CCNI | -1.688 | 0 |
| CETN2 | 1.04 | 0.736 | CETN2 | -1.173 | 0 |
| COMMD1 | -1.22 | 0.1158 | COMMD1 | -1.315 | 0 |
| COQ9 | -1.04 | 0.5042 | COQ9 | -1.49 | 0.0004 |
| COX7A2 | -1.15 | 0.1134 | COX7A2 | -2.216 | 0.0016 |
| CPD | -1.17 | 0.025 | CPD | -1.602 | 0.0012 |
| CTDSPL | 1.45 | 0.0016 | CTDSPL | -1.088 | 0.0004 |
| CTRC | 1.05 | 0.4831 | CTRC | 1.286 | 0.0037 |
| EPB41L4B | -1.11 | 0.2702 | EPB41L4B | -1.091 | 0 |
| ETV4 | -1.4 | 0.002 | ETV4 | -1.127 | 0 |
| FADD | -1.01 | 0.4299 | FADD | -1.836 | 0.0016 |
| FAM168B | 1.16 | 0.0884 | FAM168B | -1.086 | 0 |
| FBXW4 | -1 | 0.7858 | FBXW4 | -1.085 | 0 |
| FHL2 | 1.32 | 0.0024 | FHL2 | 1.036 | 0.0004 |

| | | | | | |
|---|---|---|---|---|---|
| FOXJ3 | 1.21 | 0.1518 | FOXJ3 | -1.707 | 0.0002 |
| FOXO3 | -1.02 | 0.3337 | FOXO3 | -1.109 | 0.0048 |
| FZD9 | 1.16 | 0.4699 | FZD9 | -1.455 | 0.0002 |
| GLIPR1 | 2.92 | 8.13E-07 | GLIPR1 | 1.86 | 0.0006 |
| GLUD1 | -1.23 | 0.0257 | GLUD1 | -1.716 | 0.0037 |
| HIST1H2AC | -1.12 | 0.1045 | HIST1H2AC | 1.192 | 0 |
| IDH1 | -1.5 | 0.0003 | IDH1 | -1.515 | 0 |
| LEPR; LEPROT | 1.02 | 0.8324 | LEPROT | -1.82 | 0 |
| LETM1 | -1.16 | 0.1599 | LETM1 | -1.942 | 0 |
| MDP1; NEDD8-MDI | 1 | 0.9445 | NEDD8 | -1.108 | 0 |
| MORC2 | 1.09 | 0.5599 | MORC2 | -1.324 | 0 |
| MRPL18 | 1.09 | 0.2511 | MRPL18 | -2.069 | 0 |
| NETO2 | 1.68 | 0.0003 | NETO2 | -3.022 | 0.0006 |
| NFIX | -1.05 | 0.7755 | NFIX | -1.425 | 0 |
| NQO1 | -1.71 | 4.02E-05 | NQO1 | -2.131 | 0 |
| NUDT2 | -1.01 | 0.4154 | NUDT2 | -1.186 | 0 |
| PLEKHB2 | 1 | 0.888 | PLEKHB2 | 1.044 | 0 |
| POLR2C | -1.01 | 0.3243 | POLR2C | -1.612 | 0.0029 |
| RCC2 | 1.17 | 0.035 | RCC2 | -2.127 | 0 |
| RFWD2 | -2.71 | 1.52E-05 | RFWD2 | -2.034 | 0 |
| RPAIN | 1.06 | 0.8175 | RPAIN | -1.116 | 0.0006 |
| RPUSD3 | 1.02 | 0.9947 | RPUSD3 | -1.192 | 0.0002 |
| SCMH1 | -1.03 | 0.6343 | SCMH1 | -1.061 | 0.0018 |
| SDC1 | 1.62 | 0.0018 | SDC1 | -1.255 | 0.0004 |
| SDC4 | 1.5 | 0.0033 | SDC4 | 1.044 | 0 |
| SLC31A2 | 1.04 | 0.6018 | SLC31A2 | 1.047 | 0.0006 |
| SOCS2 | 1.12 | 0.2457 | SOCS2 | 1.195 | 0 |
| SPOCK2 | -1.39 | 0.0145 | SPOCK2 | -2.314 | 0.0002 |
| ST6GALNAC6 | -1.21 | 0.0412 | ST6GALNAC6 | -1.19 | 0 |
| SURF4 | -1.05 | 0.2975 | SURF4 | -1.914 | 0 |
| TALDO1 | -1.34 | 0.013 | TALDO1 | -1.642 | 0.0018 |
| TAP1 | -1.14 | 0.2619 | TAP1 | 1.054 | 0 |
| TFPI | 1.09 | 0.2803 | TFPI | -1.798 | 0.0018 |
| THAP11 | 1.04 | 0.9195 | THAP11 | -1.541 | 0 |

| | | | | | |
|---|---|---|---|---|---|
| TIMM23 | -1.07 | 0.3857 | TIMM23 | -1.639 | 0.0008 |
| TIMM23B; AGAP6; ' | -1.29 | 0.0346 | | | |
| TIMP2; CEP295NL | 1.35 | 0.0047 | TIMP2 | 1.107 | 0 |
| TMCO3 | -1.3 | 0.0462 | TMCO3 | -1.24 | 0.004 |
| TMEM30A; COX7A2 | 1.13 | 0.0875 | | | |
| TNFRSF21 | -1.18 | 0.0422 | TNFRSF21 | -1.871 | 0 |
| TRIM35 | 1 | 0.8196 | TRIM35 | 1.117 | 0 |
| TUFT1 | -1.27 | 0.0157 | TUFT1 | 1.004 | 0 |
| UBE2G2 | -1.08 | 0.3353 | UBE2G2 | -2.013 | 0.0012 |
| UBE2Q1 | 1.04 | 0.4841 | UBE2Q1 | -1.897 | 0 |
| UBE2Z | 1.15 | 0.1203 | UBE2Z | -1.279 | 0.0008 |
| UBFD1 | -1.1 | 0.1842 | UBFD1 | -1.081 | 0.0004 |
| UBIAD1 | 1.07 | 0.2348 | UBIAD1 | -1.617 | 0 |
| ZHX3 | 1.01 | 0.3078 | ZHX3 | -1.072 | 0.0012 |
| ZNF358 | 1.01 | 0.2765 | ZNF358 | -1.14 | 0.0018 |

| **HuH7 (current)** | | | **Huh7 (Lee et al)** | | |
|---|---|---|---|---|---|
| Gene Symbol | Fold Chang | pValue | | Fold Change (Log2) | pValue |
| AARS | 1.04 | 0.8703 | AARS | -1.061 | 0 |
| ABL1 | -1.16 | 0.2362 | ABL1 | -1.006 | 0 |
| ALDH18A1 | 1.3 | 0.0277 | ALDH18A1 | -1.526 | 0.0007 |
| ARIH2 | -1.23 | 0.05 | ARIH2 | -1.112 | 0 |
| ARIH2 | 1.01 | 0.9433 | | | |
| ARIH2 | 1.09 | 0.4479 | | | |
| B4GALT5 | 1.14 | 0.8097 | B4GALT5 | -1.115 | 0.0086 |
| BOLA3 | -1.26 | 0.0952 | BOLA3 | -1.991 | 0.0092 |
| C6orf106 | 1.14 | 0.4155 | C6orf106 | -1.423 | 0.0078 |
| C17orf58 | -1.1 | 0.6015 | C17orf58 | -1.207 | 0 |
| CANX | 1.02 | 0.871 | CANX | -3.083 | 0 |
| CCNI | -1.06 | 0.5705 | CCNI | -1.819 | 0 |
| CETN2 | -1.05 | 0.5515 | CETN2 | -1.16 | 0.0006 |
| COMMD1 | -1.21 | 0.1006 | COMMD1 | -2.195 | 0.0028 |
| COQ9 | 1.06 | 0.5114 | COQ9 | -2.017 | 0 |

| | | | | | |
|---|---|---|---|---|---|
| COX7A2 | -1.07 | 0.6507 | COX7A2 | -2.826 | 0 |
| CPD | 1.12 | 0.3535 | CPD | -1.321 | 0.005 |
| CTDSPL | 1.17 | 0.2574 | CTDSPL | -2.911 | 0 |
| CTRC | -1.17 | 0.2777 | CTRC | -1.373 | 0.001 |
| EPB41L4B | -1.05 | 0.9708 | EPB41L4B | -2.199 | 0 |
| ETV4 | -1.52 | 0.0081 | ETV4 | -3.142 | 0 |
| FADD | 1.07 | 0.9915 | FADD | -1.684 | 0.0061 |
| FAM168B | 1.42 | 0.04 | FAM168B | -1.118 | 0 |
| FBXW4 | -1.13 | 0.8449 | FBXW4 | -1.03 | 0.0035 |
| FHL2 | -1.25 | 0.1981 | FHL2 | 1.138 | 0 |
| FOXJ3 | -1.04 | 0.3585 | FOXJ3 | -1.335 | 0 |
| FOXO3 | -1 | 0.9975 | FOXO3 | -1.372 | 0.0082 |
| FZD9 | 1.16 | 0.4633 | FZD9 | -3.333 | 0 |
| GLIPR1 | -1.47 | 0.2894 | GLIPR1 | 3.002 | 0.0003 |
| GLUD1 | 1.09 | 0.4917 | GLUD1 | -1.5 | 0.0097 |
| HIST1H2AC | 1.27 | 0.0658 | HIST1H2AC | 1.366 | 0.0089 |
| IDH1 | 1 | 0.9332 | IDH1 | -1.017 | 0.008 |
| LEPR; LEPROT | 1.1 | 0.8202 | LEPROT | -1.98 | 0.0014 |
| LETM1 | -1.17 | 0.4937 | LETM1 | -1.72 | 0.0003 |
| MDP1; NEDD8-MDP1; | -1.17 | 0.335 | NEDD8 | -1.146 | 0.0062 |
| MORC2 | 1.22 | 0.4537 | MORC2 | -1.156 | 0.009 |
| MRPL18 | -1.1 | 0.1999 | MRPL18 | -2.199 | 0.0049 |
| NETO2 | 1.12 | 0.8749 | NETO2 | -1.392 | 0.0025 |
| NFIX | -1.18 | 0.1033 | NFIX | -2.388 | 0 |
| NQO1 | -1.14 | 0.5158 | NQO1 | -2.418 | 0 |
| NUDT2 | 1.2 | 0.0478 | NUDT2 | -1.64 | 0 |
| PLEKHB2 | -1.14 | 0.0672 | PLEKHB2 | 1.278 | 0.0066 |
| POLR2C | 1.08 | 0.8576 | POLR2C | -1.091 | 0 |
| RCC2 | 1.33 | 0.1616 | RCC2 | -1.419 | 0.0019 |
| RFWD2 | -3.35 | 0.001 | RFWD2 | -2.695 | 0 |
| RPAIN | 1.13 | 0.2309 | RPAIN | -1.602 | 0.0015 |
| RPUSD3 | 1.1 | 0.6669 | RPUSD3 | -1.501 | 0.0025 |
| SCMH1 | -1.08 | 0.4546 | SCMH1 | -1.029 | 0.0068 |
| SDC1 | 1.15 | 0.5305 | SDC1 | -2.118 | 0 |

| | | | | | |
|---|---|---|---|---|---|
| SDC4 | 1.29 | 0.5247 | SDC4 | 1.189 | 0.0059 |
| SLC31A2 | -1.12 | 0.2158 | SLC31A2 | 1.794 | 0 |
| SOCS2 | 1.17 | 0.5212 | SOCS2 | 2.182 | 0.0008 |
| SPOCK2 | 1.07 | 0.5896 | SPOCK2 | -2.23 | 0 |
| ST6GALNAC6 | 1.25 | 0.0905 | ST6GALNAC6 | -1.538 | 0.0036 |
| SURF4 | 1.2 | 0.1395 | SURF4 | -1.886 | 0 |
| TALDO1 | -1.18 | 0.0887 | TALDO1 | -1.491 | 0.0023 |
| TAP1 | -1.14 | 0.6264 | TAP1 | 2 | 0 |
| TFPI | 1.24 | 0.3774 | TFPI | -1.28 | 0.0085 |
| THAP11 | -1.13 | 0.6441 | THAP11 | -1.442 | 0 |
| TIMM23 | -1.11 | 0.394 | TIMM23 | -1.399 | 0.0072 |
| TIMM23B; AGAP6; TIM | 1.17 | 0.3508 | | | |
| TIMP2; CEP295NL | 1.03 | 0.5754 | TIMP2 | 1.103 | 0.0004 |
| TMCO3 | 1.02 | 0.8982 | TMCO3 | -1.705 | 0 |
| TMEM30A; COX7A2 | 1 | 0.8953 | | | |
| TNFRSF21 | 1.16 | 0.4267 | TNFRSF21 | -1.276 | 0 |
| TRIM35 | 1.06 | 0.5854 | TRIM35 | 1.074 | 0.0028 |
| TUFT1 | 1.02 | 0.8536 | TUFT1 | 1.868 | 0 |
| UBE2G2 | -1.09 | 0.2776 | UBE2G2 | -1.727 | 0.002 |
| UBE2Q1 | -1.09 | 0.3168 | UBE2Q1 | -1.702 | 0 |
| UBE2Z | 1.08 | 0.4197 | UBE2Z | -1.468 | 0.0032 |
| UBFD1 | 1.2 | 0.2846 | UBFD1 | -1.449 | 0.0021 |
| UBIAD1 | 1.12 | 0.4943 | UBIAD1 | -2.501 | 0.0002 |
| ZHX3 | -1.04 | 0.6264 | ZHX3 | -1.383 | 0 |
| ZNF358 | 1 | 0.8263 | ZNF358 | -2.328 | 0.0019 |
