## Supplementary Material and Methods for "Constitutive Photomorphogenesis Protein 1 homolog (COP1) sustains nuclear factor-4 alpha function in human hepatocyte models"

**Supplementary materials**

siRNA (Silencer Select grade) were obtained from Life Technologies (Ambion):

HNF4A siRNA: s6698

control siRNA: NT #1, 4390844

MTTP siRNA: s9053

COP1 siRNA:

COP1si: s34632

COP1si-33: s34633

COP1si-91: s59691

Antibodies

COP1: A300-894A (Bethyl Laboratories)

MTTP: N-17 (SC-33116, Santa Cruz Biotechnology Inc)

HNF4A: C11F12 (Cell Signaling Technology)

TUBB: GTX11307 (GeneTex)

PCR primers used (forward, reverse)

APOC3

CTGCCCCTGTAGGTTGCTTA

GCAGCTTCTTGTCCAGCTTT

CEBPA

TGGACAAGAACAGCAACGAG

AGGCACCGGAATCTCCTAGT

COP1

TGTAGGCCTGGCTTCCAATG

CCTCCAGCACACAGCACTAA

HNF4A

GCGGAAGAACCACATGTACTC

GGCTGCTGTCCTCATAGCTT

MTTP

GTAGTCCCCGTTCGGCATCT

AGGGGTGGCTGCGATTAAGG

PPIA

ACCGTGTTCTTCGACATTGC

TTCTGTGAAAGCAGGAACCC

TRIB1

TTCAAGCAGATTGTCTCCGC

AGTGGTGTTGAGGATCTCAG

Recombinant proteins (expressed from PLVX)

HNF4A-HA sequence. HA tag in red.

MRLSKTLVDMDMADYSAALDPAYTTLEFENVQVLTMGNDTSPSEGTNLNAPNSLGVS
ALCAICGDRATGKHYGASSCDGCKGFFRRSVRKNHMYSCRFSRQCVVDKDKRNQCR
YCRLKKCFRAGMKKEAVQNERDRISTRRSSYEDSSLPSINALLQAEVLSRQITSPVSGI
NGDIRAKKIASIADVCESMKEQLLVLVEWAKYIPAFCELPLDDQVALLRAHAGEHLLLGA
TKRSMVFKDVLLLGNDYIVPRHCPELAEMSRVSIRILDELVLPFQELQIDDNEYAYLKAII
FFDPDAKGLSDPGKIKRLRSQVQVSLEDYINDRQYDSRGRFGELLLLLPTLQSITWQMI
EQIQFIKLFGMAKIDNLLQEMLLGGSPSDAPHAHHPLHPHLMQEHMGTNVIVANTMPT
HLSNGQMCEWPRPRGQAATPETPQPSPPGGSGSEPYKLLPGAVATIVKPLSAIPQPTI
TKQEVIGGAAGYPYDVPDYA

flagCOP1 sequence. Flag tag in red.

MDYKDDDDKAMSGSRQAGSGSAGTSPGSSAASSVTSASSSLSSSPSPPSVAVSAAA
LVSGGVAQAAGSGGLGGPVRPVLVAPAVSGSGGGAVSTGLSRHSCAARPSAGVGGS
SSSLGSGSRKRPLLAPLCNGLINSYEDKSNDFVCPICFDMIEEAYMTKCGHSFCYKCIH
QSLEDNNRCPKCNYVVDNIDHLYPNFLVNELILKQKQRFEEKRFKLDHSVSSTNGHRW
QIFQDWLGTDQDNLDLANVNLMLELLVQKKKQLEAESHAAQLQILMEFLKVARRNKRE
QLEQIQKELSVLEEDIKRVEEMSGLYSPVSEDSTVPQFEAPSPSHSSIIDSTEYSQPPG
FSGSSQTKKQPWYNSTLASRRKRLTAHFEDLEQCYFSTRMSRISDDSRTASQLDEFQ

ECLSKFTRYNSVRPLATLSYASDLYNGSSIVSSIEFDRDCDYFAIAGVTKKIKVYEYDTVI
QDAVDIHYPENEMTCNSKISCISWSSYHKNLLASSDYEGTVILWDGFTGQRSKVYQEH
EKRCWSVDFNLMDPKLLASGSDDAKVKLWSTNLDNSVASIEAKANVCCVKFSPSSRY
HLAFGCADHCVHYYDLRNTKQPIMVFKGHRKAVSYAKFVSGEEIVSASTDSQLKLWNV
GKPYCLRSFKGHINEKNFVGLASNGDYIACGSENNSLYLYYKGLSKTLLTFKFDTVKSV
LDKDRKEDDTNEFVSAVCWRALPDGESNVLIAANSQGTIKVLELV
